# Conserved cell types with divergent features between human and mouse cortex

**DOI:** 10.1101/384826

**Authors:** Rebecca D Hodge, Trygve E Bakken, Jeremy A Miller, Kimberly A Smith, Eliza R Barkan, Lucas T Graybuck, Jennie L Close, Brian Long, Osnat Penn, Zizhen Yao, Jeroen Eggermont, Thomas Hollt, Boaz P Levi, Soraya I Shehata, Brian Aevermann, Allison Beller, Darren Bertagnolli, Krissy Brouner, Tamara Casper, Charles Cobbs, Rachel Dalley, Nick Dee, Song-Lin Ding, Richard G Ellenbogen, Olivia Fong, Emma Garren, Jeff Goldy, Ryder P Gwinn, Daniel Hirschstein, C Dirk Keene, Mohamed Keshk, Andrew L Ko, Kanan Lathia, Ahmed Mahfouz, Zoe Maltzer, Medea McGraw, Thuc Nghi Nguyen, Julie Nyhus, Jeffrey G Ojemann, Aaron Oldre, Sheana Parry, Shannon Reynolds, Christine Rimorin, Nadiya V Shapovalova, Saroja Somasundaram, Aaron Szafer, Elliot R Thomsen, Michael Tieu, Richard H Scheuermann, Rafael Yuste, Susan M Sunkin, Boudewijn Lelieveldt, David Feng, Lydia Ng, Amy Bernard, Michael Hawrylycz, John W. Phillips, Bosiljka Tasic, Hongkui Zeng, Allan R Jones, Christof Koch, Ed S Lein

## Abstract

Elucidating the cellular architecture of the human neocortex is central to understanding our cognitive abilities and susceptibility to disease. Here we applied single nucleus RNA-sequencing to perform a comprehensive analysis of cell types in the middle temporal gyrus of human cerebral cortex. We identify a highly diverse set of excitatory and inhibitory neuronal types that are mostly sparse, with excitatory types being less layer-restricted than expected. Comparison to a similar mouse cortex single cell RNA-sequencing dataset revealed a surprisingly well-conserved cellular architecture that enables matching of homologous types and predictions of human cell type properties. Despite this general conservation, we also find extensive differences between homologous human and mouse cell types, including dramatic alterations in proportions, laminar distributions, gene expression, and morphology. These species-specific features emphasize the importance of directly studying human brain.

## Introduction

The cerebral cortex, responsible for most of our higher cognitive abilities, is the most complex structure known to biology and is comprised of approximately 16 billion neurons and 61 billion non-neuronal cells organized into approximately 200 distinct anatomical or functional regions^1,2,3,4^. The human cortex is greatly expanded relative to the mouse, the dominant model organism in basic and translational research, with a 1200-fold increase in cortical neurons compared to only a 60-fold increase in sub-cortical neurons (excluding cerebellum) ^5,6^. The general principles of neocortical development and the basic multilayered cellular cytoarchitecture of the neocortex appear relatively conserved across mammals ^7,8^. However, whether the cellular and circuit architecture of cortex is fundamentally conserved across mammals, with a massive evolutionary areal expansion of a canonical columnar architecture in human, or is qualitatively and quantitatively specialized in human, remains an open question long debated in the field ^9,10^. Addressing this question has been challenging due to a lack of tools to broadly characterize cell type diversity in complex brain regions, particularly in human brain tissues.

Prior studies have described differences in the cellular makeup of the cortex in human and specialized features of specific cell types ^11,12,13,14,15,16,17^, although the literature is remarkably limited. For example, the supragranular layers of cortex, involved in cortico-cortical communication, are differentially expanded in mammalian evolution ^18^. Furthermore, certain cell types show highly specialized features in human and non-human primate compared to mouse, such as the interlaminar astrocytes^17^, and the recently described rosehip cell ^19^, a type of inhibitory interneuron in cortical layer 1 with distinctive morpho-electrical properties. All of these cellular properties are a function of the genes that are actively used in each cell type, and transcriptomic methods provide a powerful method to understand the molecular underpinnings of cellular phenotypes as well as a means for mechanistic understanding of species-specialized phenotypes. Indeed, a number of studies have shown significant differences in transcriptional regulation between mouse, non-human primate and human, including many genes associated with neuronal structure and function ^20,21,22,23^.

Dramatic advances in single cell transcriptional profiling present a new approach for large-scale comprehensive molecular classification of cell types in complex tissues, and a metric for comparative analyses. The power of these methods is fueling ambitious new efforts to understand the complete cellular makeup of the mouse brain ^24^ and the even the whole human body ^25^. Recent applications of single cell RNA-sequencing (scRNA-seq) methods in mouse cortex have demonstrated robust transcriptional signatures of neuronal and non-neuronal cell types ^26,27,28^, and suggest the presence of approximately 100 neuronal and non-neuronal cell types in any given cortical area. Similar application of scRNA-seq to human brain has been challenging due to the difficulty in dissociating intact cells from densely interconnected human tissue ^29^. In contrast, single nucleus RNA-sequencing (snRNA-seq) methods allow for transcriptional profiling of intact neuronal nuclei that are relatively easy to isolate and enable use of frozen postmortem specimens from human brain repositories ^30,31,32^. Importantly, it was recently shown that single nuclei contain sufficient gene expression information to distinguish closely related subtypes of cells at a similar resolution to scRNA-seq ^33,34^, demonstrating that snRNA-seq is a viable method for surveying cell types that can be compared to scRNA-seq data. Early applications of snRNA-seq to human cortex demonstrated the feasibility of the approach but have not provided depth of coverage sufficient to achieve similar resolution to mouse studies ^35^.

The current study aimed to establish a robust methodology for relatively unbiased cell type classification in human brain using snRNA-seq, and to perform the first comprehensive comparative analysis of cortical cell types to understand conserved and divergent features of human and mouse cerebral cortex. We first describe the cellular landscape of the human cortex, and then demonstrate a similar degree of cellular diversity between human and mouse and a conserved set of homologous cell types and subclasses. In contrast, we present evidence for extensive differences between homologous types, including evolutionary changes in relative proportions, laminar distributions, subtype diversity, gene expression and other cellular phenotypes.

## Results

### Transcriptomic taxonomy of cell types

A robust snRNA-seq methodology was established to analyze transcriptomically defined cell types in human cortex. We focused on the middle temporal gyrus (MTG), with samples largely derived from high-quality postmortem brain specimens. This region is frequently available through epilepsy surgery resections, permitting a comparison of postmortem versus acute neurosurgical tissues, as well as allowing future correlation with *in vitro* slice physiology experiments in MTG. Frozen tissue blocks were thawed, vibratome sectioned, and stained with fluorescent Nissl dye. Individual cortical layers were microdissected, tissues were homogenized to release nuclei, and nuclei were stained with an antibody against NeuN to differentiate neuronal (NeuN-positive) and non-neuronal (NeuN-negative) nuclei. Single nuclei were collected via fluorescence-activated cell sorting (FACS) (Fig. 1A, Extended Data Figure 1A, **Methods**). We sorted ~90% NeuN-positive and ~10% NeuN-negative nuclei across all cortical layers to enrich for neurons. The final dataset contained less than the targeted 10% non-neuronal nuclei because nearly 50% of NeuN-negative nuclei failed quality control criteria, potentially due to the lower RNA content of glia compared to neurons (**Methods**)^27^. SMART-Seqv4 (Takara Bio USA Inc.) was used to reverse transcribe mRNA and amplify cDNA. Sequencing libraries were generated using Nextera XT (Illumina), which were sequenced on a HiSeq 2500 at a median depth of 2.6 +/- 0.5 million reads/nucleus. Nuclei were collected from 8 total human tissue donors (4 male, 4 female; 4 postmortem, 4 neurosurgical) ranging in age from 24 to 66 years (Extended Data Table 1). 15,206 nuclei were collected from postmortem tissue donors with no history of neuropathology or neuropsychiatric disorders, and 722 nuclei came from apparently histologically normal MTG distal to pathological tissue that was removed during surgical resections to treat epilepsy (**Methods**).

**Figure 1.**
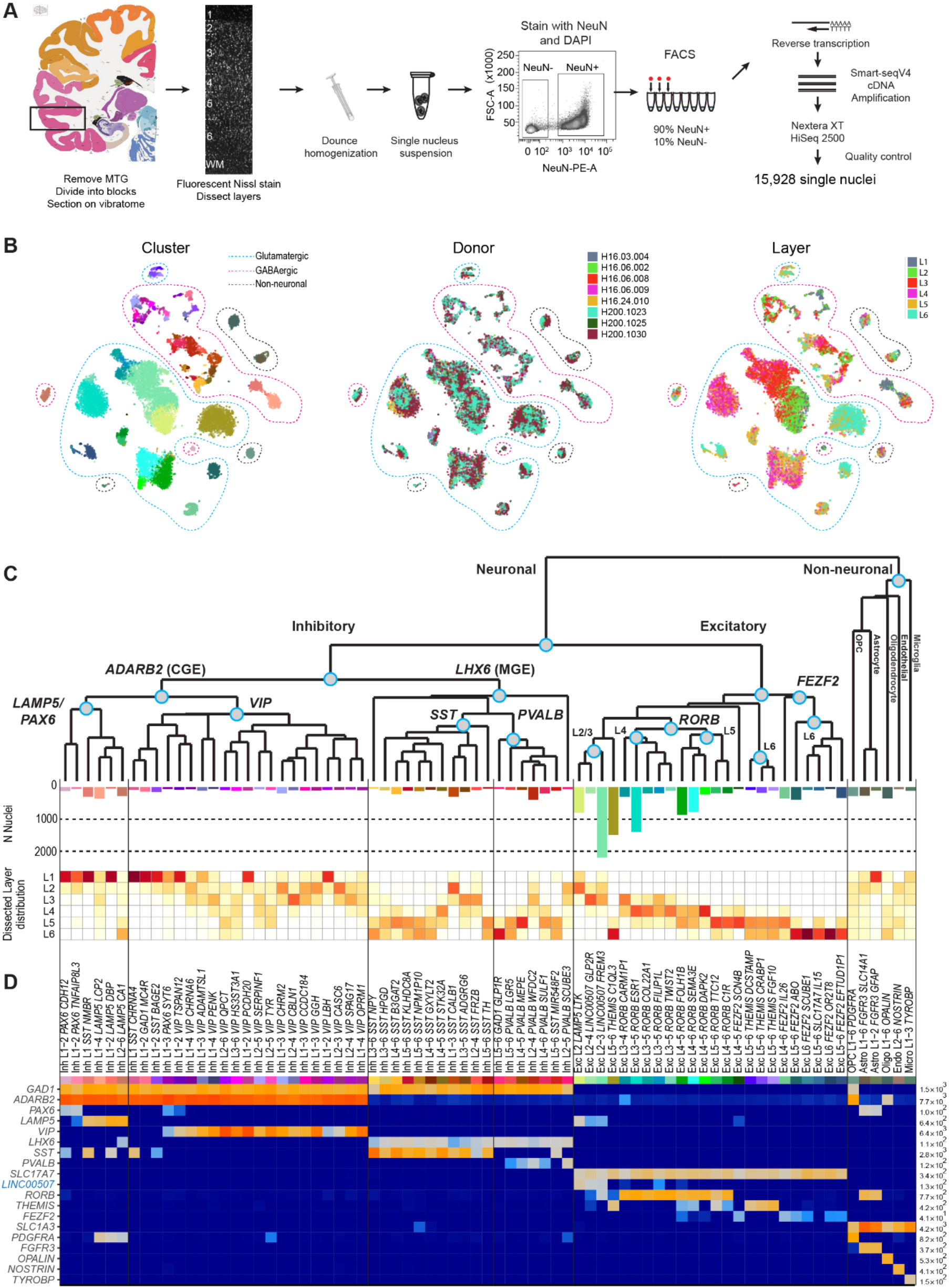
Cell type taxonomy in human middle temporal gyrus (MTG) **(A)** Schematic diagram illustrating nuclei isolation from frozen MTG specimens by vibratome sectioning, fluorescent Nissl staining and dissection of specific cortical layers. Single neuronal (NeuN+) and non-neuronal (NeuN-) nuclei were collected by fluorescence-activated cell sorting (FACS), and RNA-sequencing of single nuclei used SMART-seqv4, Nextera XT, and HiSeq2500 sequencing. **(B)** Overview of transcriptomic cell type clusters visualized using t-distributed stochastic neighbor embedding (t-SNE). On the left t-SNE map, each dot corresponding to one of 15,928 nuclei has a cell-type specific color that is used throughout the remainder of the manuscript. In the middle, donor metadata is overlaid on the t-SNE map to illustrate the contribution of nuclei from different individuals to each cluster. In the list of specimens, H16.03.004-H16.06.009 are neurosurgical tissue donors and H16.24.010-h200.1030 are postmortem donors. On the right, layer metadata is overlaid on the t-SNE map to illustrate the laminar composition of each cluster. **(C)** Hierarchical taxonomy of cell types based on median cluster expression consisting of 69 neuronal (45 inhibitory, 24 excitatory) and 6 non-neuronal transcriptomic cells types. Major cell classes are labeled at branch points in the dendrogram. The bar plot below the dendrogram represents the number of nuclei within each cluster. The laminar distributions of clusters are shown in the plot that follows. For each cluster, the proportion of nuclei in each layer is depicted using a scale from white (low) to dark red (high). **(D)** Heatmap showing the expression of cell class marker genes (blue, non-coding) across clusters. Maximum expression values for each gene are listed on the far-right hand side. Gene expression values are quantified as counts per million of intronic plus exonic reads and displayed on a log10 scale.

To evenly survey cell type diversity across cortical layers, nuclei were sampled based on the relative proportion of neurons in each layer reported in human temporal cortex^36^. Based on Monte Carlo simulations, we estimated that 14,000 neuronal nuclei were needed to target types as rare as 0.2% of the total neuron population (**Methods**). Using an initial subset of RNA-seq data, we observed more transcriptomic diversity in layers 1, 5, and 6 than in other layers so additional neuronal nuclei were sampled from those layers. In total, 15,928 nuclei passed quality control criteria and were split into three broad classes of cells (10,708 excitatory neurons, 4297 inhibitory neurons, and 923 non-neuronal cells) based on NeuN staining and cell class marker gene expression (**Methods**).

Nuclei from each broad class were iteratively clustered as described in^33^. Briefly, high variance genes were identified while accounting for gene dropouts, expression dimensionality was reduced with principal components analysis (PCA), and nuclei were clustered using Jaccard-Louvain community detection (**Methods**). On average, neuronal nuclei were larger than non-neuronal nuclei (Extended Data Fig. 1B), and median gene detection (Extended Data Fig. 1C,D) was correspondingly higher for neurons (9046 genes) than for non-neuronal cells (6432 genes), as previously reported for mouse^26,27,28^. Transcriptomic cell types were largely conserved across diverse individuals and tissue types (postmortem, neurosurgical), since all curated clusters contained nuclei derived from multiple donors, and nuclei from postmortem and neurosurgical tissue types clustered together (Fig. 1B, Extended Data Fig. 2A). However, a small, but consistent expression signature related to tissue type was apparent; for example, nuclei derived from neurosurgical tissues exhibited higher expression of some activity related genes (Extended Data Fig. 2). 325 nuclei were assigned to donor-specific or outlier clusters that contained marginal quality nuclei and were excluded from further analysis (**Methods**).

This analysis method defined 75 transcriptomically distinct cell types, including 45 inhibitory neuron types that express the canonical GABAergic interneuron marker *GAD1*, 24 excitatory neuron types that express the vesicular glutamate transporter *SLC17A7*, and 6 non-neuronal types that express the glutamate transporter *SLC1A3* (Fig. 1C, D). As expected based on prior studies^26,27,28,31^, the hierarchical relationships among types roughly mirrors the developmental origin of different cell types. We refer to the cell type clusters as cell *types*, intermediate order nodes as *subclasses*, and higher order nodes such as the interneurons derived from the caudal ganglionic eminence (CGE) as *classes*, and the broadest divisions such as excitatory neurons as *major classes*. Neuronal types split into two major classes representing cortical plate-derived glutamatergic excitatory neurons (n=10,525 nuclei) and ganglionic eminence-derived GABAergic inhibitory neurons (n=4164 nuclei). Non-neuronal types (n=914 nuclei) formed a separate main branch based on differential expression of many genes (Fig. 1C). We developed a principled nomenclature for clusters based on: 1) major cell class, 2) layer enrichment (including layers containing at least 10% of nuclei in that cluster), 3) a subclass marker gene (maximal expression of 14 manually-curated genes), and 4) a cluster-specific marker gene (maximal detection difference compared to all other clusters) (Fig. 1D, Extended Data Fig. 3, **Methods**). For example, the left-most inhibitory neuron type in Figure 1D, found in samples dissected from layers 1 and 2, and expressing the subclass marker *PAX6* and the specific marker *CDH12*, is named Inh L1-2 *PAX CDH12*. Additionally, we generated a searchable semantic representation of these cell type clusters that incorporates this accumulated knowledge about marker gene expression, layer enrichment, specimen source, and parent cell class to link them to existing anatomical and cell type ontologies^37^ (**Supplementary Data**). We find broad correspondence to an earlier study^31^, but identify many additional types of excitatory and inhibitory neurons due to increased sampling and/or methodological differences (Extended Data Fig. 4). The majority of cell types were rare (<100 nuclei per cluster, <0.7% of cortical neurons), including almost all interneuron types and deep layer excitatory neuron types. In contrast, the excitatory neurons of superficial layers 2-4 were dominated by a small number of relatively abundant types (>500 nuclei per cluster, >3.5% of neurons) (Fig. 1C). Both excitatory types and many interneuron types were restricted to a few layers, whereas non-neuronal nuclei were distributed across all layers, with the notable exception of one astrocyte type (Fig. 1C).

### Excitatory neurons often span multiple layers

The 24 transcriptionally distinct excitatory neuron types broadly segregated by layer and expressed known laminar markers (Fig. 2A-C). In general, excitatory types were most similar to other types in the same or adjacent layers. Transcriptomic similarity by proximity for cortical layers has been described before, and interpreted as a developmental imprint of the inside-out generation of cortical layers^38^. Complex relationships between clusters are represented as constellation diagrams (Fig. 2A, **Methods**)^26^, where the circles represent core cells that were most transcriptionally similar to the cluster to which they were originally assigned, and indicate the size (proportional to circle area) and average laminar position of each cell type. The thickness of lines between cell clusters represents their similarity based on the number of nuclei whose assignment to a cluster switched upon reassignment (intermediate cells, **Methods**). This similarity by proximity is also apparent in the hierarchical dendrogram structure of cluster similarity in Figure 2B. One exception is the layer 5 Exc L5-6 *THEMIS C1QL3* type, which has a transcriptional signature similar to layer 2 and 3 types as well as several deep layer cell types (Fig. 2A, B). Two types, Exc L4-5 *FEZF2 SCN4B* and Exc L4-6 *FEZF2 IL26*, were so distinct that they occupied separate branches on the dendrogram and did not connect via intermediate cells to any other type (Fig. 2A, B).

**Figure 2.**
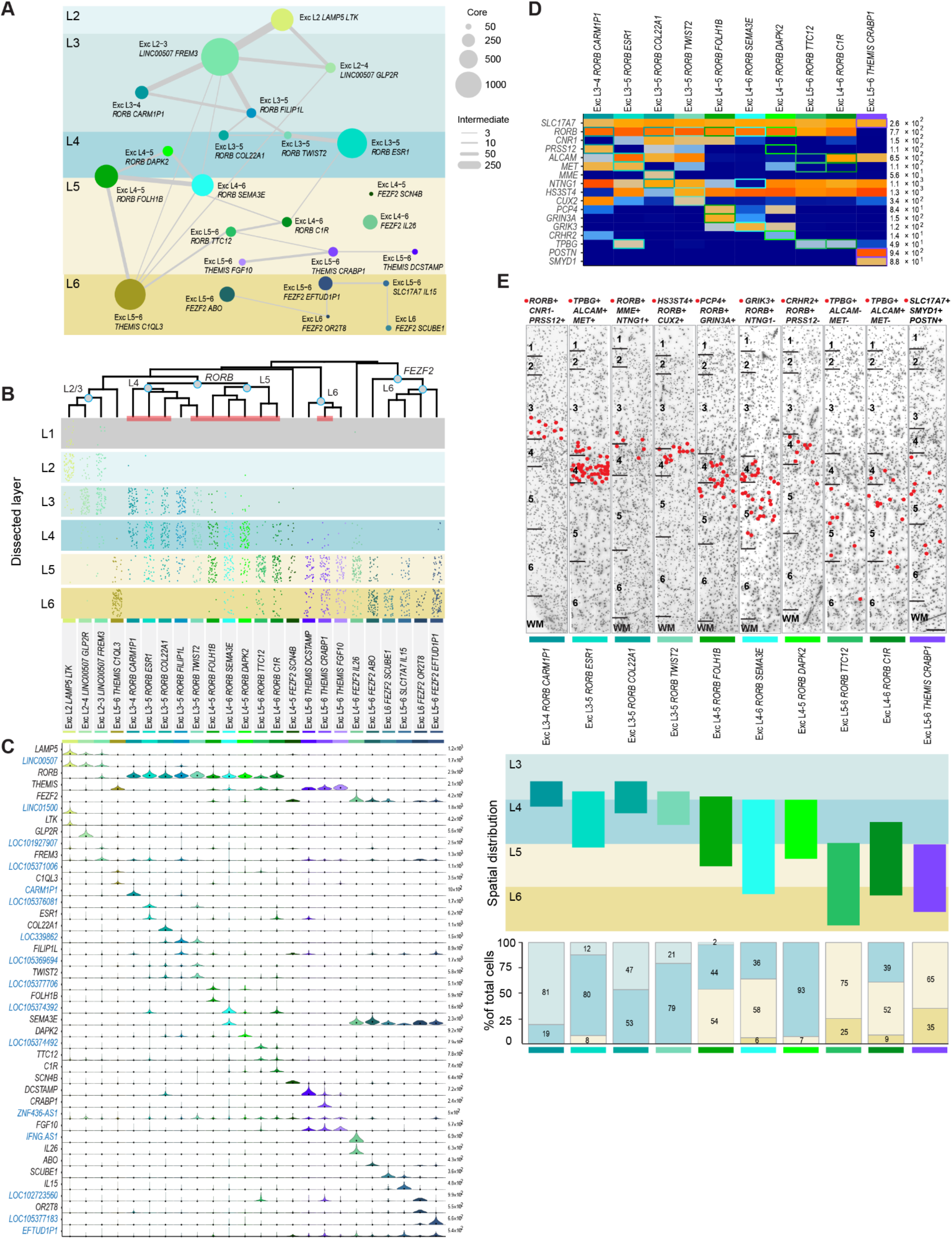
Excitatory neuron diversity and marker gene expression. **(A)** Constellation diagram for excitatory cell types. The number of cells that could be unambiguously assigned to each cluster (core cells) is represented by disc area and the number of cells with uncertain membership between each pair of clusters (intermediate cells) is represented by line thickness. **(B)** Dendrogram illustrating overall gene expression similarity between cell types. Layer distributions of cell types are shown as dot plots where each dot represents a single nucleus from a layer-specific dissection. Note that incidental capture of some layer 2 excitatory neurons occurred in layer 1 dissections and is reflected in the dot plots. Clusters marked by a red bar at the base of the dendrogram are examined using fluorescent in situ hybridization (FISH) in (D-E). **(C)** Violin plot showing marker gene (blue, non-coding) expression distributions across clusters. Each row represents a gene, black dots show median gene expression within clusters, and the maximum expression value for each gene is shown on the right-hand side of each row. Gene expression values are shown on a linear scale. **(D)** Heatmap summarizing combinatoral 3-gene panels used for multiplex fluorescent in situ hybridization assays to explore the spatial distribution of 10 excitatory clusters. Gene combinations for each cluster are indicated by colored boxes on the heatmap. **(E)** Representative inverted images of DAPI-stained cortical columns spanning layers 1-6 for each marker gene panel. Red dots depict the locations of cells positive for the specific marker gene combinations for each cluster. Marker gene combinations are listed at the top of each image. Cluster names along with color coded cluster-specific bars are beneath each panel. Scale bar, 250μm. Below the DAPI images, a schematic diagram of the spatial distribution (i.e. the laminar extent) of each cluster examined. The schematic is based on the observed positions of labeled cells across n=3-4 sections per cell type and n=2-3 donors per cell type. Bar plots below summarize counts of the percentage of labeled cells per layer, expressed as a fraction of the total number of labeled cells for each type. Bars are color coded to represent different cortical layers using the scheme shown in (A). The cluster represented by each bar is indicated by the colored bar at the bottom of the plot. Cell counts are cumulative values from n=2-3 subjects for each cell type.

Each excitatory type showed selective expression of genes that can be used as cell type markers (Fig. 2C), although in general a small combinatorial profile (generally 2-3 genes per type) was necessary to distinguish each type from all other cortical cell types (Fig. 2D). The majority of these markers are novel as excitatory neuron markers, and belonged to diverse and functionally important gene families, such as BHLH transcription factors (*TWIST2*), collagens (*COL22A1*), and semaphorins (*SEMA3E*). Surprisingly, 16 out of 37 (41%) of these most specific marker genes were unannotated loci (LOCs), long non-coding RNAs (lincRNA), pseudogenes, and antisense transcripts. This may partially be a result of profiling nuclear RNA, as some of these transcripts have been shown to be enriched in the nucleus (Fig. 2C, Extended Data Figs. 3, 5)^39^.

Unexpectedly, most excitatory neuron types were present in multiple layers based on layer dissection information (Fig. 2B). Within the supragranular layers, three main types were enriched in layer 2 and 3 dissections. Additionally, ten RORB-expressing types were enriched in layer 3-6 dissections (Fig. 2B, C). Layers 5 and 6 contained 11 excitatory types: 4 types that expressed *THEMIS* (Thymocyte Selection Associated), 6 types that expressed *FEZF2*, and 1 type that expressed the cytokine *IL15* (Interleukin 15). The majority of these types were similarly represented in layer 5 and 6 dissections (Fig. 2B). To clarify whether this crossing of layer boundaries was an artifact of dissection or a feature of MTG organization, we investigated the layer distribution of 10 types using multiplex fluorescence *in situ* hybridization (FISH) with combinatorial gene panels designed to discriminate clusters (Fig. 2B, D, Extended Data Fig.6). *In situ* distributions largely validated snRNA-seq predictions (Fig. 2E). Three types were mainly localized to layer 3c and the upper part of layer 4, defined as the dense band of granule cells visible in Nissl stained sections (Fig 2E). Interestingly, one of these types (Exc L3-4 *RORB CARM1P1*) had large nuclei, suggesting that it may correspond to a subset of the giant pyramidal layer 3c neurons previously described in MTG^40^ (Fig. 2E, Extended Data Fig. 6). Two types were mostly restricted to layer 4 (Exc L3-5 *RORB ESR1*, Exc L4-5 *RORB DAPK2*), but the five other types examined all spanned multiple layers (Fig. 2E). Taken together, the snRNA-seq and *in situ* validation data indicate that transcriptomically defined excitatory neuron types are frequently not layer-specific, but rather spread across multiple anatomically defined layers.

### Heterogeneous expression within clusters

A major evolutionary feature of human cortical architecture is the expansion of supragranular layers compared to other mammals, and morphological and physiological properties of pyramidal neurons vary across layers 2 and 3 of human temporal cortex^40,41^. In that light, it was surprising to find only three main excitatory clusters in human cortical layers 2 and 3. However, one cluster was very large (Exc L2-3 *LINC00507 FREM3;* n=2284 nuclei) and spanned layers 2 and 3, posing the possibility that there is significant within-cluster heterogeneity. Indeed, we find continuous variation in gene expression in this cluster along the axis of cortical depth, illustrated well using two data visualization and mining tools built for this project to allow public access to this dataset. The Cytosplore MTG Viewer (https://viewer.cytosplore.org), is an extension of Cytosplore^42^, and presents a hierarchy of t-SNE maps of different subsets of MTG clusters^43^, with each map defined using informative marker genes (Fig. 3A). Layer dissection metadata overlaid onto the t-SNE map of Exc *L2-3 LINC00507 FREM3 revealed* that nuclei in this type were ordered by layer, with nuclei sampled from layers 2 and 3 occupying relatively distinct locations in t-SNE space. Selecting nuclei at both ends of the cluster gradient in t-SNE space and computing differential expression between these nuclei revealed a set of genes with variable expression across this cluster (Fig. 3A, **Supplementary Movie 1**).

**Figure 3.**
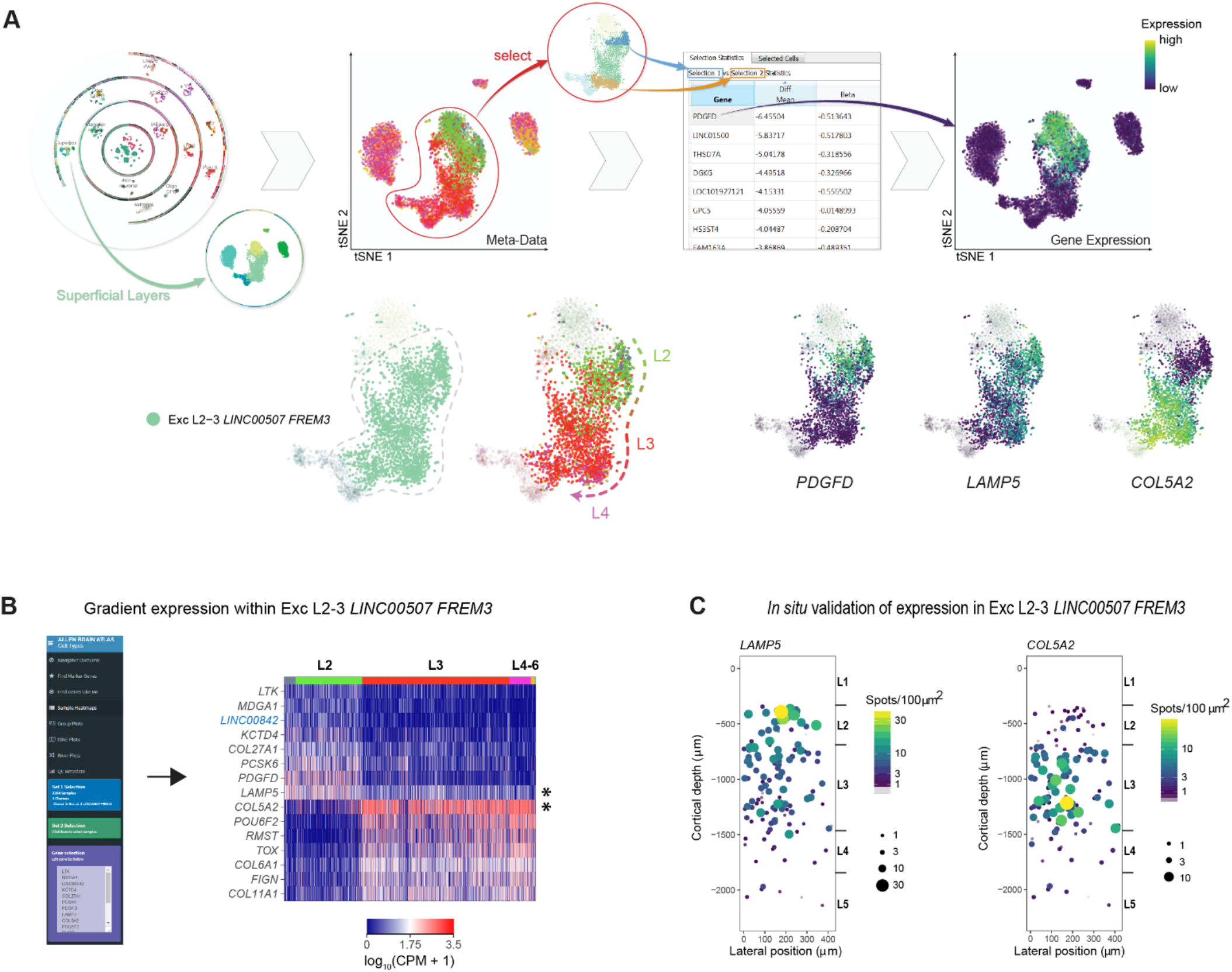
Gene expression heterogeneity within the Exc L2-3 *LINC00507 FREM3* cell type. **(A, B)** Transcriptomics data visualization tools for exploring gene expression gradients in human cortical neurons. **(A)** Cytosplore MTG Viewer. Top panels, left to right: the hierarchy viewer shows an overview of the t-SNE map of all clusters. Zooming in allows for visualization and selection of superficial layer excitatory neurons on the t-SNE map. Overlaying layer metadata on the t-SNE map shows that nuclei within the EXC L2-3 *LINC00507 FREM3* cell type are sorted by cortical layer. Differential expression analysis, computed by selecting nuclei on opposite ends of the cluster, reveals gene expression gradients organized along the layer structure of the cluster. Bottom panels, left to right: t-SNE map showing the EXC L2-3 *LINC00507 FREM3* cluster outlined by dashed gray line. Overlaying layer metadata on the cluster highlights its layer structure. Examples of genes that exhibit expression heterogeneity across the layer structure of the cluster are shown to the right. **(B)** RNA-Seq Data Navigator. Selection of the sample heatmaps option in the browser allows for visualization of gene expression patterns in the EXC L2-3 *LINC00507 FREM3* cluster. Each row in the heatmap represents a gene (blue, non-coding), and nuclei in the cluster are ordered by layer (colored bar at the top of the heatmap). The selected genes illustrate opposing gene expression gradients across the layer structure of the cluster. Genes marked with an asterisk were included in the validation experiments in (C). **(C)** Single molecule fluorescent in situ hybridization (smFISH) validation of gene expression heterogeneity. Panels show quantification of *LAMP5* (left) and *COL5A2* (right) expression in cells located in layers 2-3. Each circle represents a cell, the size of each circle is proportional to the number of smFISH spots per cell, and circles are color-coded per the scale shown to the right of each panel. Consistent with the RNA-seq data shown in panels A and B, smFISH analysis demonstrates that these genes exhibit opposing expression gradients across cortical layers 2 and 3.

Examining this set of variable genes within Exc L2-3 *LINC00507 FREM3* using the RNA-Seq Data Navigator (http://celltypes.brain-map.org/rnaseq/human) showed gradient expression between layers 2 and 3 (Fig. 3B). Finally, single molecule FISH confirmed gradient expression of *LAMP5* and *COL5A2* across layers 2 and 3 in cells mapping to this cluster (Fig. 3C,Extended Data Figs. 7, 8). These results illustrate that there is additional diversity in human supragranular pyramidal neurons manifested as continuous variation in gene expression as a function of cortical depth that likely correlates with anatomical and functional heterogeneity of those cells.

### Inhibitory neuron diversity

GABAergic inhibitory neurons split into two major branches, largely distinguished by expression of Adenosine Deaminase, RNA Specific B2 (*ADARB2*) and the transcription factor LIM Homeobox 6 (*LHX6*) (Fig. 4A-F). In mouse cortex, interneurons split into the same two major branches, also defined by expression of *Adarb2* and *Lhx6* and developmental origins in the caudal ganglionic eminence (CGE) and medial ganglionic eminence (MGE), respectively^26^. The *ADARB2* branch was further subdivided into the *LAMP5/PAX6* and *VIP* subclasses of interneurons, with likely developmental origins in the CGE. Surprisingly, the serotonin receptor subunit *HTR3A*, which marks CGE-derived interneurons in mouse^44^, was not a good marker of these types in human (Fig. 4E). The *LHX6* branch consisted of *PVALB* and *SST* subclasses of interneurons, likely originating in the medial ganglionic eminence MGE^45,46^. Consistent with mouse cortex^26^, the *ADARB2* branch showed a much higher degree of diversity in supragranular layers 1-3 compared to layers 4-6, whereas the opposite was true for the *LHX6* branch (Fig. 4A, B). As with the excitatory neuron taxonomy, many interneuron cluster specific markers were unannotated (LOC) genes, lincRNAs, pseudogenes, and antisense transcripts (Fig. 4E, F).

**Figure 4.**
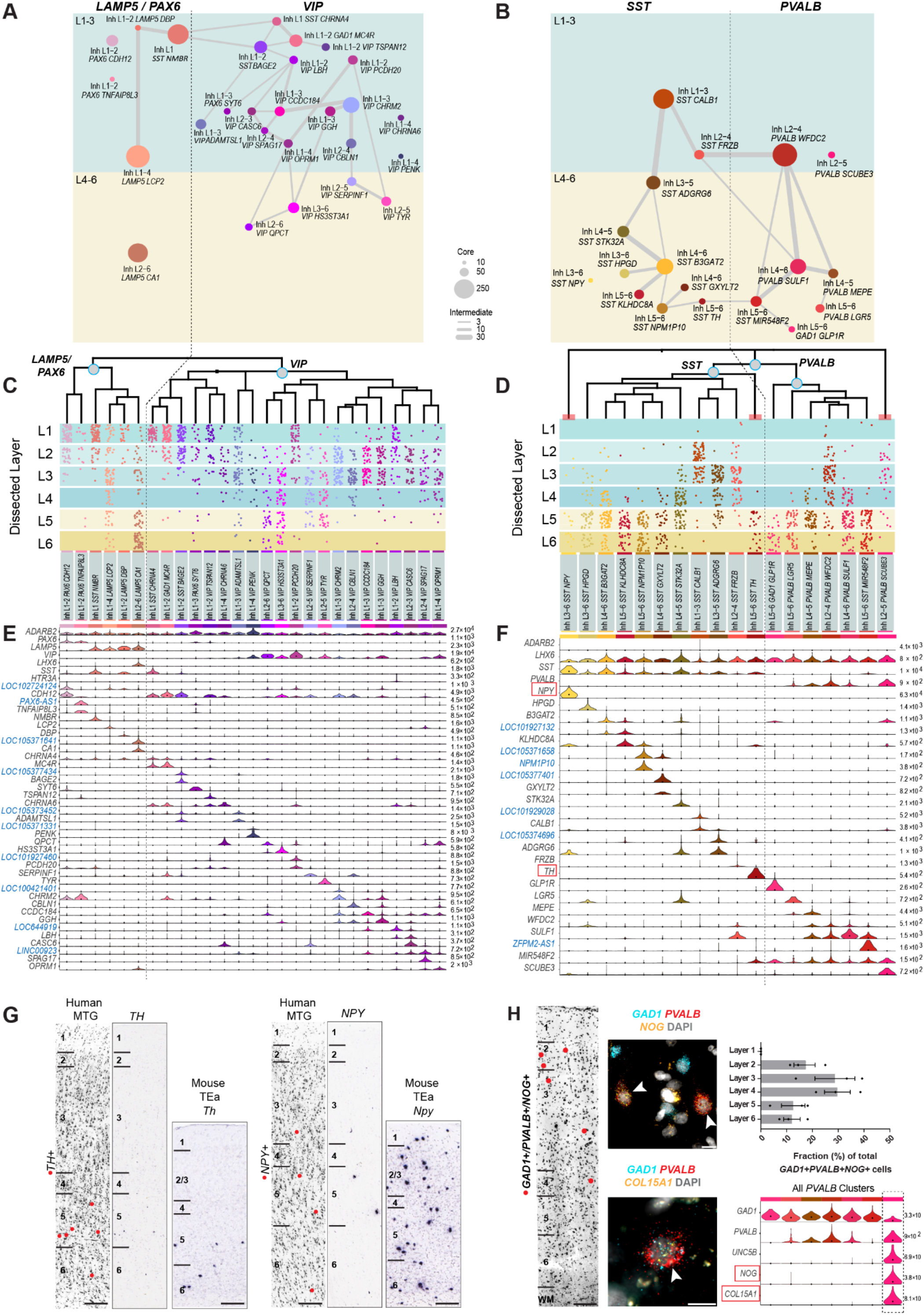
Inhibitory neuron diversity and marker gene expression. **(A, B)** Constellation diagrams for *LAMP5/PAX6* and *VIP* (A) and *SST/PVALB* (B) subclasses. The number of core cells within each cluster is represented by disc area and the number of intermediate cells by weighted lines. **(C, D)** Dendrograms illustrate gene expression similarity between cell types. Below each dendrogram, the spatial distribution of each type is shown. Each dot represents a single nucleus derived from a layer-specific dissection. Red bars at the base of the dendrogram in (D) indicate clusters examined using *in situ* hybridization (ISH) in (G-H). **(E, F)** Violin plots of marker gene expression distributions across clusters. Rows are genes (blue, non-coding transcripts), black dots in each violin represent median gene expression within clusters, and the maximum expression value for each gene is shown on the right-hand side of each row. Gene expression values are shown on a linear scale. Genes shown in **(G)** are outlined by red boxes in (F). **(G)** Chromogenic single gene ISH for *TH* (left), a marker of Inh L5-6 *SST TH*, and *NPY* (right), a marker of Inh L3-6 *SST NPY*, from the Allen Human Brain Atlas. Left columns show grayscale images of the Nissl stained section nearest the ISH stained section shown in the right panel for each gene. Red dots overlaid on the Nissl section show the laminar positions of cells positive for the gene assayed by ISH. Chromogenic ISH for *Th* and *Npy* in mouse temporal association cortex (TEa) from the Allen Mouse Brain Atlas are to the right of the human ISH images. Scale bars: human (250μm), mouse (100μm). **(H)** RNAscope mutiplex fluorescent ISH for markers of putative chandelier cell cluster Inh L2-5 *PVALB SCUBE3*. Left panel - representative inverted DAPI-stained cortical column with red dots marking the position of cells positive for the genes *GAD1, PVALB*, and *NOG* (scale bar, 250μm). Middle - images of cells positive for *GAD1, PVALB*, and the specific marker genes *NOG* (top, scale bar 10μm) and *COL15A1* (bottom, scale bar 10um). White arrows mark triple positive cells. Right - bar plot summarizes counts of *GAD1*+, *PVALB+, NOG+* cells across layers (expressed as percentage of total triple positive cells). Bars show the mean, error bars represent the standard error of the mean (SEM), and dots represent data points for individual specimens (n=3 subjects). Violin plot shows gene expression distributions across clusters in the *PVALB* subclass for the chandelier cell marker *UNC5B* and the Inh L2-5 *PVALB SCUBE3* cluster markers *NOG* and *COL15A1*.

The *LAMP5/PAX6* subclass of interneurons included 6 transcriptomic types, many of which were enriched in layers 1 and 2 (Fig. 4C). Several types coexpressed *SST* (Fig. 4E), consistent with previous reports demonstrating *SST* expression in layer 1 of human MTG^19^ and different from mouse *Lamp5* and *Pax6* interneurons^26,27^, which do not express *SST*. The Inh L1-4 *LAMP5 LCP2* type expressed marker genes of rosehip cells, a type of interneuron with characteristic large axonal boutons that we described in a previous study of layer 1 MTG interneurons^19^. With whole cortex coverage, it is clear that this type is not restricted to layer 1 but rather present across all cortical layers. Among *LAMP5/PAX6* types on the *ADARB2* (CGE-derived) branch, Inh L2-6 *LAMP5 CA1* cells uniquely expressed *LHX6*, suggesting possible developmental origins in the MGE, and appear similar to the *Lamp5 Lhx6* cells previously described in mouse cortex^26,27^.

*VIP* interneurons represented the most diverse subclass, containing 21 transcriptomic types (Fig. 4A), many of which were enriched in layers 2 and 3 (Fig. 4C). Several types in the *VIP* subclass (Inh L1 *SST CHRNA4* and Inh L1-2 *SST BAGE2*) appeared to be closely related to the L1 *SST NMBR* type of the *LAMP5/PAX6* subclass, as evidenced by intermediate cell connections between these types. Interestingly, these highly related types were all localized to layers 1 and 2. Furthermore, while both the Inh L1 *SST CHRNA4* and Inh L1-2 *SST BAGE2* were grouped into the *VIP* subclass, they appeared to lack expression of *VIP*. Rather, they expressed *SST*, consistent with expression of this gene in layer 1 and 2 interneurons as discussed above (Fig. 4A, C, E)^19^. The Inh L1-2 *GAD1 MC4R* type also lacked expression of *VIP* (Fig. 4E). Notably, this type specifically expresses the Melanocortin 4 Receptor, a gene linked to autosomal dominant obesity and previously shown to be expressed in a population of mouse hypothalamic neurons that regulate feeding behavior^48,49^.

The *SST* subclass consisted of 11 transcriptomic types, including one highly distinct type, Inh L3-6 *SST NPY*, that occupied its own discrete branch on the dendrogram and was not connected to other types in the *SST* constellation (Fig. 4B, D). Several *SST* types displayed laminar enrichments, with Inh L5-6 *SST TH* cells being a particularly restricted type, found only in layers 5 and 6. We further validated marker gene expression and the spatial distribution of the Inh L3-6 *SST NPY* and Inh L5-6 *SST TH* types using ISH from the Allen Human Brain Atlas (http://human.brain-map.org/; Fig. 4G). ISH for *TH* confirmed that expression of this gene is sparse and restricted to layers 5-6; interestingly, *Th* ISH in mouse temporal association area (TEa; the closest homolog to human MTG) showed similar sparse labeling restricted to layers 5 and 6, suggesting that this gene may mark similar cell types in human and mouse (http://mouse.brain-map.org/; Fig. 4G). In contrast, the well-known interneuron marker neuropeptide Y (*Npy*) was broadly expressed in a scattered pattern throughout all layers in mouse TEa, whereas, in human MTG, *NPY* labeled only a single interneuron type whose sparsity was confirmed by ISH (Fig. 4G), indicating that this heavily-studied marker labels a different cohort of cell types in human and mouse^50,51^.

The *PVALB* subclass comprised 7 clusters, including two types that were grouped into this branch but did not appear to express *PVALB* (Fig. 4F). One of these types, Inh L5-6 *SST MIR548F2*, had low expression of *SST*, whereas the other type, Inh L5-6 *GAD1 GLP1R*, did not express any canonical interneuron subclass markers. Intermediate cells connected the Inh L5-6 *SST MIR548F2* type in the *PVALB* constellation to the Inh L5-6 *SST TH* type in the *SST* constellation. Two other connections between the *SST* and *PVALB* constellations were apparent, both of which included the Inh L2-4 *SST FRZB* cluster (Fig. 4B). One highly distinctive *PVALB* type (Inh L2-5 *PVALB SCUBE3*) (Fig. 4B, D) likely corresponds to chandelier (axo-axonic) cells as it expresses *UNC5B*, a marker of chandelier (axo-axonic) cells in mouse^52^ (Fig. 4H). Multiplex FISH (RNAscope, **Methods**) validated expression of several novel marker genes (*NOG, COL15A1*, Fig. 4H) and showed enrichment of these cells mainly in layers 2-4, consistent with the pattern observed in the snRNA-seq data (Fig. 4D, H).

### Diverse morphology of astrocyte types

Although non-neuronal (NeuN-) cells were not sampled as deeply as neurons, all major glial types - astrocytes, oligodendrocytes, endothelial cells, and microglia - were identified (Fig. 5A). In contrast to studies of mouse cortex where non-neuronal cells were more extensively sampled or selectively targeted with Cre lines^26,28,53^, we did not find other types of immune or vascular cells. This decreased diversity is likely largely due to more limited non-neuronal sampling, but may also reflect the age of tissue analyzed. For example, previous reports showed that adult mouse cortex contains mainly oligodendrocyte progenitor cells (OPCs) and mature oligodendrocytes, but few immature and myelinating oligodendrocyte types^28,53^, similarly, we found only two oligodendrocyte types, one of which expressed markers of oligodendrocyte progenitor cells (OPCs) (e.g. *PDGFRA, OLIG2*) and another that expressed mature oligodendrocyte markers (e.g. *OPALIN, MAG*) (Fig. 5A, B).

**Figure 5.**
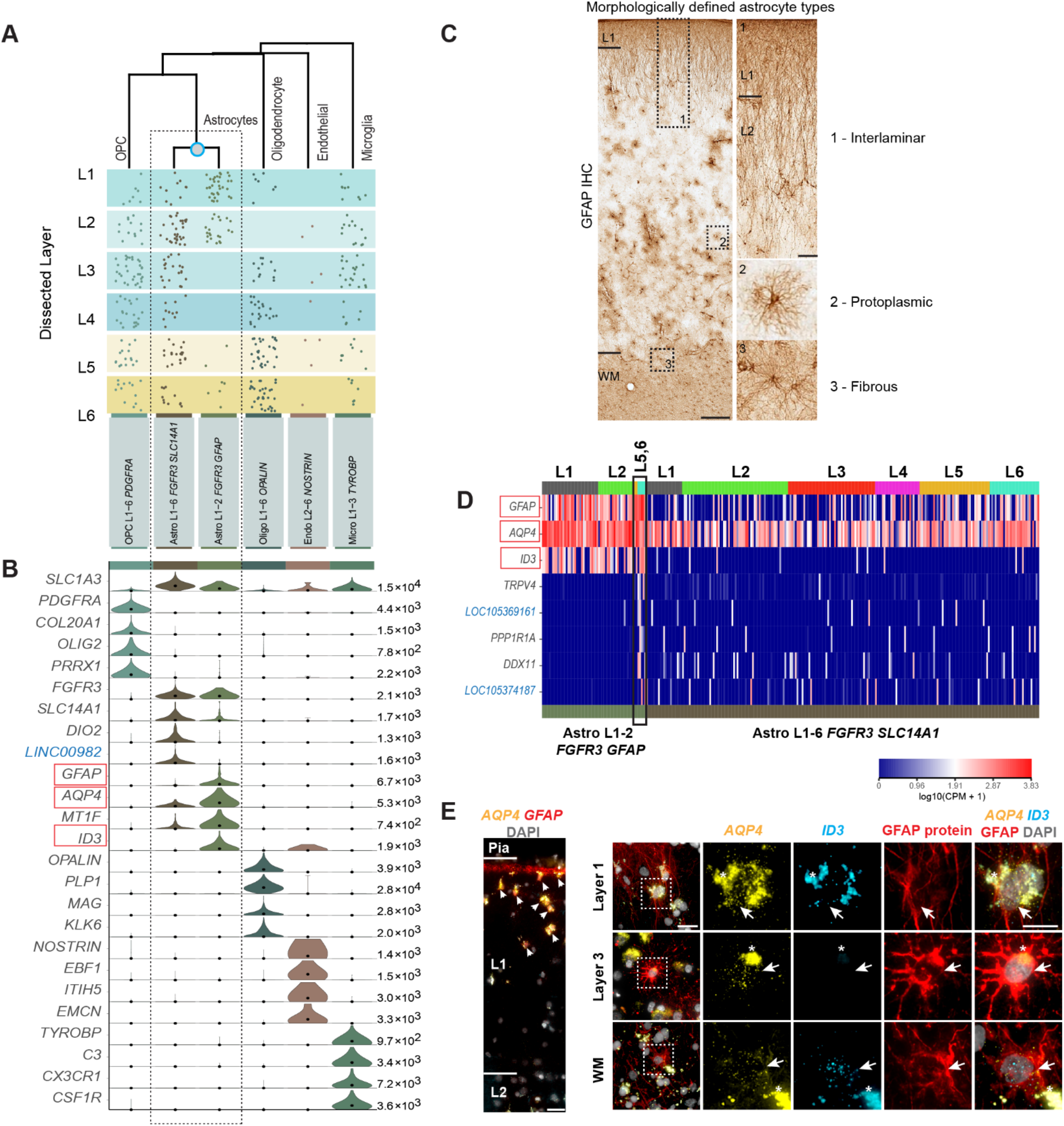
Non-neuronal cell type diversity and marker gene expression. **(A)** Dendrogram illustrating overall gene expression similarity between non-neuronal cell types, with the spatial distribution of types shown beneath the dendrogram. Each dot represents a single nucleus from a layer-specific dissection. **(B)** Violin plots show expression distributions of marker genes across clusters. Each row represents a gene (blue, non-coding), black dots represent median gene expression within clusters, and the maximum expression value for each gene is shown on the right-hand side of each row. Gene expression values are shown on a linear scale. **(C)** Immunohistochemistry (IHC) for GFAP in human MTG illustrates the features of morphologically-defined astrocyte types. Black boxes on the left panel indicate regions shown at higher magnification on the right. Scale bars: low mag (250μm), high mag (50μm). **(D)** Heatmap illustrating marker gene expression in the Astro L1-2 *FGFR3 GfAP* and Astro L1 —6 *FGFR3 SLC14A1* clusters. Each row is a gene, each column a single nucleus, and the heatmap is ordered per the layers that nuclei were dissected from. A minority of nuclei in the Astro L1—2 *FGFR3 GFAP* cluster came from deep layers (black box on heatmap) and express marker genes distinct from the other nuclei in the cluster. Red boxes in (B, D) are genes examined in (E). (E) RNAscope multiplex fluorescent *in situ* hybridization (FISH) for astrocyte markers. Left - expression of *AQP4* and *GFAP* in layer 1 (scale bar, 25μm). Cells expressing high levels of *AQP4* and *GFAP*, consistent with the Astro L1 —2 *FGFR3 GFAP* cluster, are localized to the top half of layer 1 (white arrowheads). Right - FISH for *AQP4* and *ID3* combined with GFAP immunohistochemistry. White box indicates area shown at higher magnification to the right. Scale bars: low mag (25μm), high mag (15μm). Asterisks mark lipofuscin autofluoresence. Top row: *AQP4* expressing cells in layer 1 coexpress *ID3* and have long, GFAP-labeled processes that span layer 1. Middle row: protoplasmic astrocyte located in layer 3 lacks expression of *ID3*, consistent with the Astro L1 —6 *FGFR3 SLC14A1* type. Bottom row: fibrous astrocyte at the white matter (WM)/layer 6 boundary triple positive for *AQP4, ID3*, and GFAP protein.

Astrocytes in human cortex are both functionally^54^ and morphologically^17^ specialized in comparison to rodent astrocytes, with distinct morphological types residing in different layers of human cortex (Fig. 5C). Interlaminar astrocytes, described only in primates to date, reside in layer 1 and extend long processes into lower layers, whereas protoplasmic astrocytes are found throughout cortical layers 2-6^17^ (Fig.5C). Similarly, we find two astrocyte clusters with different laminar distributions. Astro L1-2 *FGFR3 GFAP* originated mostly from layer 1 and 2 dissections, whereas the Astro L1-6 *FGFR3 SLC14A1* type was found in all layers (Fig.5A). The two astrocyte types we identified were distinguished by expression of the specific marker gene *ID3* along with higher expression of *GFAP* and *AQP4* in the Astro L1-2 *FGFR3 GFAP* type than in the Astro L1-6 *FGFR3 SLC14A1* type (Fig. 5B, D). To determine if these two transcriptomic types correspond to distinct morphological types, we labeled cells with a combination of multiplex FISH and immunohistochemistry for GFAP protein. Cells with high *GFAP* and *AQP4* expression, characteristic of the Astro L1-2 *FGFR3 GFAP* type and consistent with previous reports of interlaminar astrocytes^55^, were present predominantly in the upper half of layer 1 (Fig. 5E). Coexpression of *AQP4* and *ID3* was apparent in layer 1 cells that had extensive, long-ranging GFAP-positive processes characteristic of interlaminar astrocytes (Fig. 5E). In contrast, GFAP-positive cells with protoplasmic astrocyte morphology lacked expression of *ID3*, consistent with the Astro L1-6 *FGFR3 SLC14A1* type (Fig. 5E).

Interestingly, while most nuclei contributing to the Astro L1-6 *FGFR3 GFAP* cluster came from layer 1 and 2 dissections, seven nuclei were from layer 5 and 6 dissections and expressed *ID3* as well as a distinct set of marker genes (Fig. 5D). Based on their laminar origin, we hypothesized that these nuclei may correspond to fibrous astrocytes, which are enriched in white matter^17^ (Fig. 5C). Indeed, astrocytes at the border of layer 6 and the underlying white matter coexpressed *ID3* and *AQP4* and had relatively thick, straight GFAP-positive processes characteristic of fibrous astrocytes (Fig. 5E), suggesting that the Astro L1-6 *FGFR3 GFAP* cluster contains a mixture of two different morphological astrocyte types. Given that nuclei corresponding to fibrous astrocytes express distinct marker genes from interlaminar astrocytes (Fig. 5D), it is likely that fibrous astrocytes will form a separate transcriptomic type with increased sampling.

### Human and mouse cell type homology

Single cell transcriptomics not only provides a new method for comprehensive analysis of species-specific cellular diversity, but also a quantitative metric for comparative analysis between species. Furthermore, identification of homologous cell types or classes allows inference of cellular properties from much more heavily studied model organisms. The availability of densely sampled single cell or single nucleus RNA-seq datasets in human (described here) and mouse^26^ cortex using the same RNA-seq profiling platform allowed a direct comparison of transcriptomic cell types. The success of such a comparison is predicated on the idea of conserved transcriptional patterning. As a starting point, we asked whether the same types of genes discriminate human interneuron cell types as those reported for mouse interneuron types^52^. Indeed, we find the same sets of genes (mean = 21 genes/set) best discriminate human interneuron types (Fig.6A), including genes central to neuronal connectivity and signaling. Similar functional classes of genes also discriminate human and mouse excitatory neuron types (although with less conservation for classes of genes that discriminate non-neuronal cell types; Extended Data Fig.9A), indicating that shared expression patterns between species may facilitate matching cell types.

**Figure 6.**
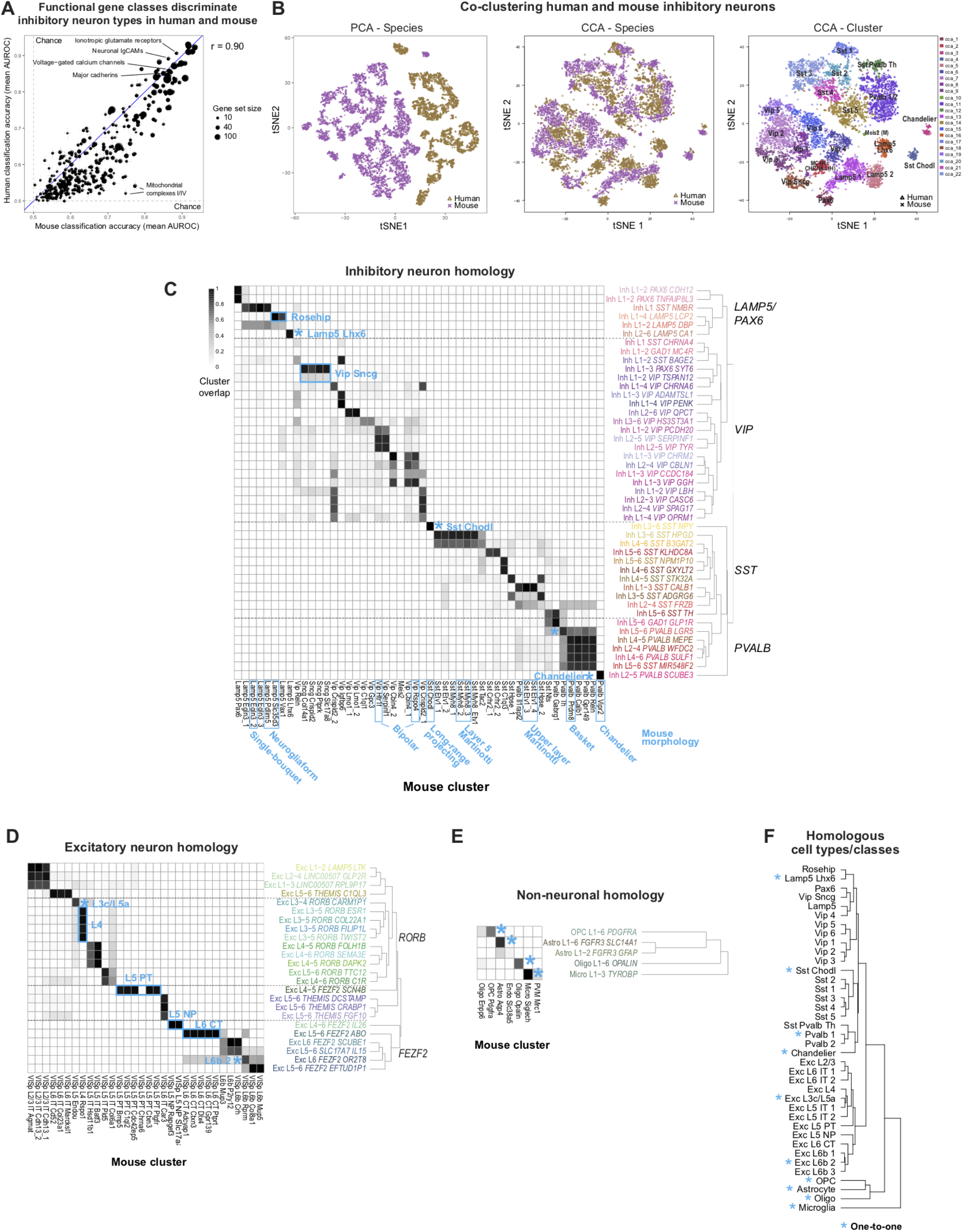
Evolutionary conservation of cell types between human and mouse. **(A)** neurons can be assigned to cell types based on expression patterns of functional gene families (n = 5 to 261 genes). Classification performance (average MetaNeighbor AUROC score across clusters) varies among functional classes of genes and is highly correlated (r = 0.90) between human and mouse. Error bars correspond to the standard deviation of average AUROC scores across ten sub-sampled iterations. **(B)** Human (gold) and mouse (purple) inhibitory neurons were aligned with principal components analysis (PCA; left) and canonical correlation analysis (CCA; middle), and the first 30 basis vectors were represented using t-SNE. Right: CCA clusters were identified by the Louvain algorithm using 30 nearest neighbors and annotated based on cluster labels from this study and mouse. Clusters labeled with (M) or (H) contain only mouse cells or human nuclei, respectively. **(C-E)** Human and mouse cell type homologies for inhibitory neurons (C), excitatory neurons (D), and non-neuronal cells (E) were predicted based on shared CCA cluster membership. Greyscale indicates, for each pair of human (rows) and mouse (columns) clusters, the minimum proportion of human nuclei or mouse cells that co-cluster using CCA. Note that rows and columns need not sum to one because clusters can partially overlap. One-to-one matches are indicated by an asterisk. Known morphologies are indicated for mouse inhibitory types and known projection targets are given for excitatory types (IT - intratelencephalic, PT - pyramidal tract/sub-cortical, NP - near-projecting, CT - corticothalamic). Note that human endothelial nuclei could not be aligned by CCA and were excluded from the analysis. **(F)** Hierarchical taxonomy of 34 neuronal and 4 nonneuronal homologous cell types and cell classes, including 10 cell types that match one-to-one between human and mouse.

Simply combining expression data for inhibitory neuron nuclei from human MTG and for cells from mouse V1 was not sufficient for identification of homologous cell types. PCA analysis resulted in samples clearly separated by species along the first principal component that explained almost 20% of expression variation (Fig.6B, Extended Data Fig.9B). Recent work has demonstrated the power of canonical correlation analysis (CCA) to align single cell RNA-seq data from human and mouse based on shared co-expression patterns^56^. Application of CCA and graph-based clustering to human and mouse cortical samples was much more successful (Fig.6B), and allowed matching of human and mouse types based on shared CCA cluster membership for inhibitory neurons (Fig.6C, Extended Data Fig.9E), excitatory neurons (Fig.6D, Extended Data Fig.9F) and non-neuronal cells (Fig.6E, Extended Data Fig.9G).

Remarkably, shared co-expression between mouse V1 and human MTG enabled the identification of homologous types at approximately half the resolution of the full human classification (38 types versus 75 types). Combining the CCA results allowed generation of a hierarchical taxonomy including 34 neuronal and 4 non-neuronal cell types and subclasses (Fig. 6F). A hybrid nomenclature from human and mouse^27^ was used to describe these homologous types. Ten cell types were matched one-to-one between species, whereas other types were matched at a subclass resolution. Transcriptomically distinct cell types more often had one-to-one matches, likely because more redundant marker genes compensated for divergent expression patterns, and we find even most rare types had homologous types in mouse and human.

This homology alignment enabled prediction of the anatomical, functional, and connectional properties of human cell types based on the much larger mouse literature for homologous cell types. For example, the human cluster Inh L2-5 *PVALB SCUBE3* described above matches one-to-one with the mouse chandelier (or axo-axonic) cell type *Pvalb Vipr2*, suggesting that this cell type selectively innervates the axon initial segment of excitatory neurons. Also, the human cluster Inh L3-6 *SST NPY* matches the mouse *Sst Chodl* type and is therefore predicted to have long-range projections and contribute to sleep regulation^26,57,58^. Many other anatomically defined interneuron types could be similarly inferred, including basket, Martinotti, bipolar, neurogliaform, and single-bouquet cells (Fig. 6C), although future experiments will be necessary to confirm these predictions.

The long-range projection targets of human glutamatergic neurons (e.g. intratelencephalic (IT), pyramidal tract (PT), and corticothalamic (CT)) that would otherwise be experimentally inaccessible can also be inferred based on their best transcriptomic match to mouse cell types; for example, the human Exc L4-5 *FEZF2 SCN4B* type corresponds to the PT sub-cortically projecting layer 5 pyramidal cells (Fig. 6D). The Exc L4-6 *FEZF2 IL26* matches two mouse layer 5 types (L5 NP *Slc17a8* and L5 NP *Rapgef3*) that lack long-range projections^26,59^. Finally, layer 6b (subplate) types can be identified by homology, and among human layer 6b types, Exc L6 *FEZF2 OR2T8* has much larger nuclei (Extended Data Fig. 1B) and corresponds to the mouse L6b *Rprm* type that selectively projects to thalamus rather than cortex.

Four of five human non-neuronal cell types matchedmouse cell types (Fig. 6E), while endothelial cells had such divergent global expression patterns between species that they could not be matched by CCA despite the expression of conserved canonical marker genes (e.g. *EMCN* and *NOSTRIN*). The mouse Oligo *Enpp6* cluster partially overlapped nuclei from human OPC and mature oligodendrocyte clusters and appears to represent an immature oligodendrocyte type^26,53^ that is rare or not present in adult human cortex. The morphologically distinct human layer 1 astrocyte type, Astro L1-2 *FGFR3 GFAP*, did not match any clusters from^26^, although a layer 1 enriched astrocyte with shared marker gene expression was previously reported in mouse^28^. Finally, while the majority of human microglia clustered with mouse microglia, two nuclei clustered with mouse perivascular macrophages (Extended Data Fig.9D), suggesting that this rare type was likely undersampled in human.

Only three mouse neuronal types and two human interneuron types lacked homologous types, although all three mouse types are very rare and may not have been sampled in human. The mouse *Meis2* inhibitory type, which is primarily restricted to white matter and has an embryonic origin outside of the ganglionic eminence^26^, may have been missed due to limited sampling of layer 6b and underlying white matter. Mouse Cajal-Retzius cells are glutamatergic neurons in layer 1. These cells are exceedingly rare (less than 0.1% of layer 1 neurons) in adult human cortex^60,61^ and were not expected to be sampled. Finally, the mouse layer 5 pyramidal tract type L5 PT *Chrna6*, a rare excitatory neuron type with strong projections to superior colliculus^59^, has no matching human cluster. However, 2 of 25 nuclei from the human pyramidal tract (PT)-like cluster Exc L4-5 *FEZF2 SCN4B* are more similar to this distinct mouse PT type than to other mouse PT types (Extended Data Fig.9C), suggesting this mismatch is also due to undersampling in human. Interestingly, both human interneuron types that lack closely matched mouse homologues (Inh L1 *SST CHRNA4* and Inh L1-2 *GAD1 MC4R*) are highly enriched in layer 1. Along with the phenotypic specialization of the layer 1 rosehip neuron^19^, it appears that layer 1 may be a hotspot of evolutionary change at the level of inhibitory cell types.

While many homologous subclasses had comparable diversity between species, some subclasses had expanded diversity in human or mouse. Human layer 4 excitatory neurons are more diverse than those of mouse (Fig. 6D), contributing to increased diversity of supragranular layers due to mixing into layer 3 as described above. Mouse layer 5 PT types are much more diverse than those in human, which may reflect either a true species difference or undersampling, as they make up <1% of layer 5 excitatory neurons in human MTG. Layer 6 CT types also show greater diversity in mouse V1 than human MTG; however, this difference may reflect an areal difference between a primary sensory area that has strong, reciprocal connnections with the thalamus and an area of association cortex. Indeed, we find increased diversity of cell types in human visual cortex that match mouse layer 6 CT types (data not shown).

### Divergent proportions of cell types

Alterations in the relative proportions of cell types could have profound consequences for cortical circuit function. snRNA-seq data predicted a significant species difference in the proportions of interneuron classes. Human MTG showed similar proportions of MGE-derived (44% *LHX6+* nuclei) and CGE-derived (50% *ADARB2+* nuclei) interneurons, whereas in mouse cortex roughly 70% of interneurons are MGE-derived and ~30% are CGE-derived^44,62^. To validate these differences, we applied multiplex FISH to quantify the proportions of CGE (*ADARB2+*) and MGE (*LHX6+*) interneurons in human MTG and mouse TEa (Fig. 7, Extended Data Fig. 10). Interneurons that co-expressed *ADARB2* and *LHX6*, corresponding to the human Inh L2-6 *LAMP5 CA1* and mouse *Lamp5 Lhx6* types (Figs. 1, 4), were considered separately. Consistent with the snRNA-seq data, we found similar proportions of MGE (50.2 ± 2.3%) and CGE (44.2 ± 2.4%) interneurons in human MTG, whereas we found more than twice as many MGE (67.8 ± 0.9%) than CGE (30.8 ± 1.2%) interneurons in mouse TEa. The increased proportion of CGE-derived interneurons in human was greatest in layer 4, whereas the decreased proportion of MGE interneurons in human was greatest in layers 4-6 (Fig. 7A). Interestingly, both the snRNA-seq data (6.1% of *GAD1+* cells) and *in situ* cell counts (5.6 ± 0.3% of *GAD1+* cells) confirmed a significant increase in the proportion of the Inh L2-6 *LAMP5 CA1* type in human MTG versus the *Lamp5 Lhx6* type in mouse TEa (1.4 ± 0.2% of *GAD1+* cells), most notably in layer 6 (Fig. 7A).

**Figure 7.**
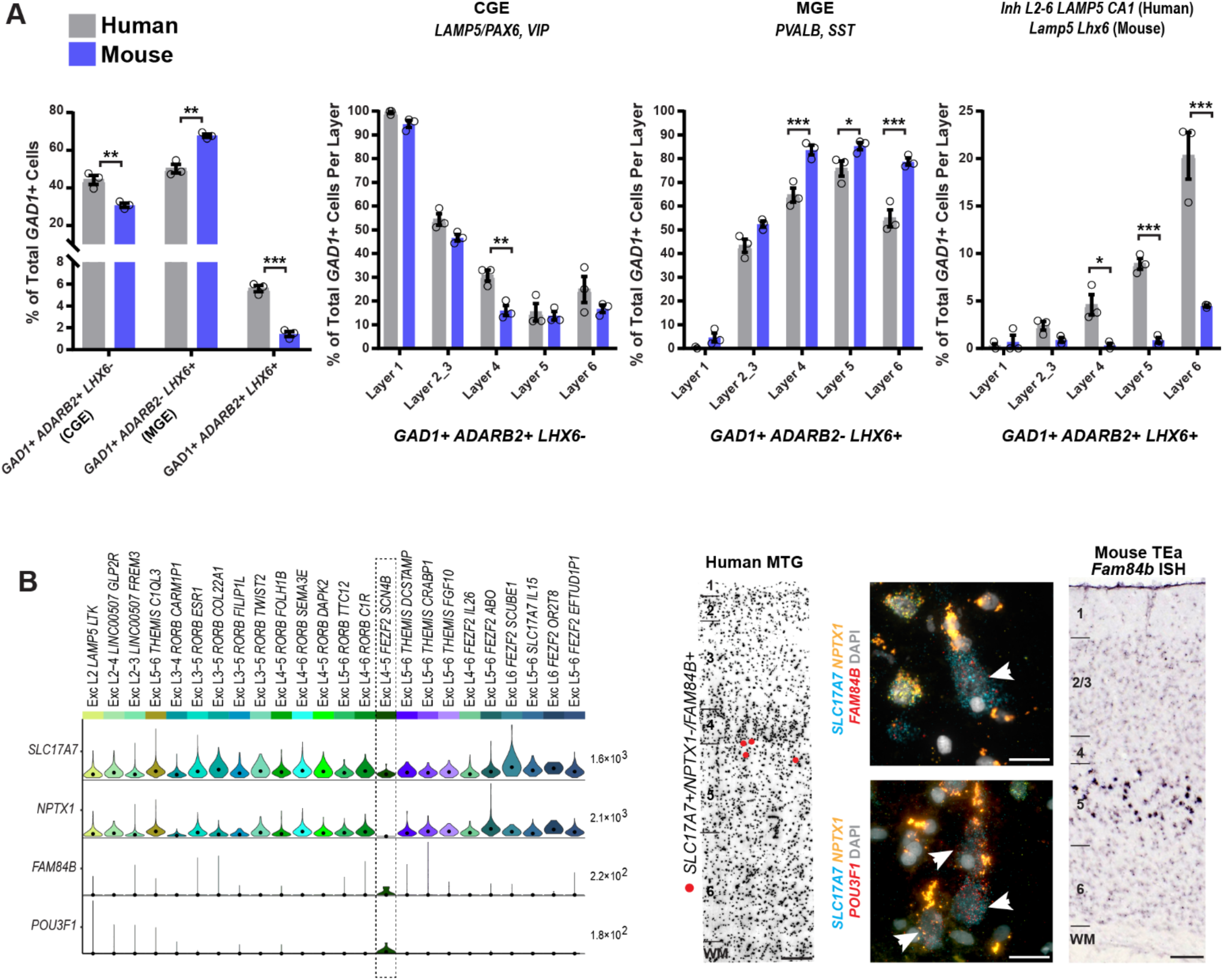
Frequency differences in cell classes and types between human and mouse. **(A)** Quantification of broad interneuron classes in human MTG and mouse temporal association area (TEa) based on counts of cells labeled using RNAscope multiplex fluorescent in situ hybridization (FISH). Sections were labeled with the gene panel *GAD1/Gad1*, *LHX6/Lhx6*, and *ADARB2/Adarb2* (human/mouse). Bar plots from left to right: (1) Percentage of each major cell class of total *GAD1+* cells. (2) Percentage of *GAD1+/ADARB2+/LHX6-* cells of total *GAD1+* cells per layer, representing *LAMP5/PAX6*, and *VIP* types. (3) Percentage of *GAD1+/ADARB2-/LHX6+* cells of total *GAD1+* cells per layer, representing all *PvALB* and *SST* types. (4) Percentage of *GAD1+/ADARB2+/LHX6+* cells of total *GAD1+* cells per layer, representing the Inh L2-6 *LAMP5 CA1* type (human) or Lamp5 Lhx6 type (mouse). Bars represent the mean, error bars the standard error of the mean, and circles show individual data points for each specimen (n=3 specimens for both human and mouse; t-test with Holm-Sidak correction for multiple comparisons, *p<0.05 **p<0.01, ***p<0.001). **(B)** Left to right: violin plot showing expression of specific markers of the putative pyramidal tract (PT) EXC L4-5 *FEZF2 SCN4B* cluster (black box) and *NPTX1*, a gene expressed by all non-PT excitatory neurons. Each row represents a gene, the black dots in each violin represent median gene expression within clusters, and the maximum expression value for each gene is shown on the right-hand side of each row. Gene expression values are shown on a linear scale. A representative inverted DAPI-stained cortical column (scale bar, 200μm) with red dots marking the position of cells positive for the genes *SLC17A7* and *FAM84B* and negative for *NPTX1* illustrates the relative abundance of the EXC L4-5 *FEZF2 SCN4B* type in human MTG. Representative examples of RNAscope multiplex FISH stained sections from human MTG showing *FAM84B* (top, white arrows, scale bar, 25μm) and *POU3F1*-expressing cells (bottom, white arrows, scale bar, 25μm). Expression of *Fam84b* in mouse TEa (scale bar, 75μm) is shown in the adjacent panel.

Another major predicted mismatch was seen for the sub-cortically projecting PT neurons, which comprise approximately 20% of layer 5 excitatory neurons in mouse but less than 1% in human based on single cell^26^ and single nucleus RNA-seq sampling. To directly compare the spatial distribution and abundance of PT types between species, we performed ISH for a pan-layer 5 PT marker (*Fam84b*)^26^ in mouse TEa and for markers of the homologous layer 5 PT type Exc L4-5 *FEZF2 SCN4B* in human MTG. In mouse TEa, *Fam84b* was expressed in many neurons in superficial layer 5 (Fig. 7B). To unambiguously label PT neurons in human MTG, we performed triple FISH with the pan-excitatory marker *SLC17A7*, the PT markers *FAM84B* or *POU3F1*, and *NPTX1*, which labels most SLC17A7-positive layer 5 neurons but not PT cells (Fig. 7B, Extended Data Fig. 11). In MTG, *SLC17A7+/NPTX1-* cells co-labeled with *FAM84B* or *POU3F1 were* sparsely distributed predominantly in superficial layer 5 and were large with a prominent, thick apical dendrite (Fig. 7B, Extended Data Fig. 11). Thus, PT cells have a similar distribution within layer 5 in human and mouse but are much less abundant in human, likely reflecting an evolutionary scaling constraint as discussed below.

### Divergent expression between homologous types

The identification of homologous or consensus cell types or classes allows direct analysis of the conservation and divergence of gene expression patterns across these types. For each pair of homologous cell types, we compared expression levels of 14,414 orthologous genes between human and mouse. Nuclear expression levels were estimated based on intronic reads to better compare human single nucleus and mouse single cell RNA-seq data. The Exc L3c/L5a type (Exc L3-4 *RORB CARM1P1* in human) has the most conserved expression (r = 0.78) of all types, and yet 12% of genes have highly divergent expression (defined as >10-fold difference), including many specific markers (orange dots, Fig. 8A) for this cell type. Microglia had the least conserved expression (r = 0.60), and more than 20% of genes were highly divergent (Fig.8B). Surprisingly, the Exc L3c/L5a consensus type shows a striking shift in layer position between human, where Exc L3-4 *RORB CARM1P1* is highly enriched in layer 3c of MTG, and mouse, where the homologous type L5 *Endou* is enriched in layer 5a of mouse V1 (Fig.8A). This laminar shift of a homologous cell type helps explain the reported expression shift of several genes from layer 5 in mouse to layer 3 in human^20^, including two genes (*BEND5* and *PRSS12*) expressed in Exc L3-4 *RORB CARM1P1* but not in layer 3 of mouse TEa.

**Figure 8.**
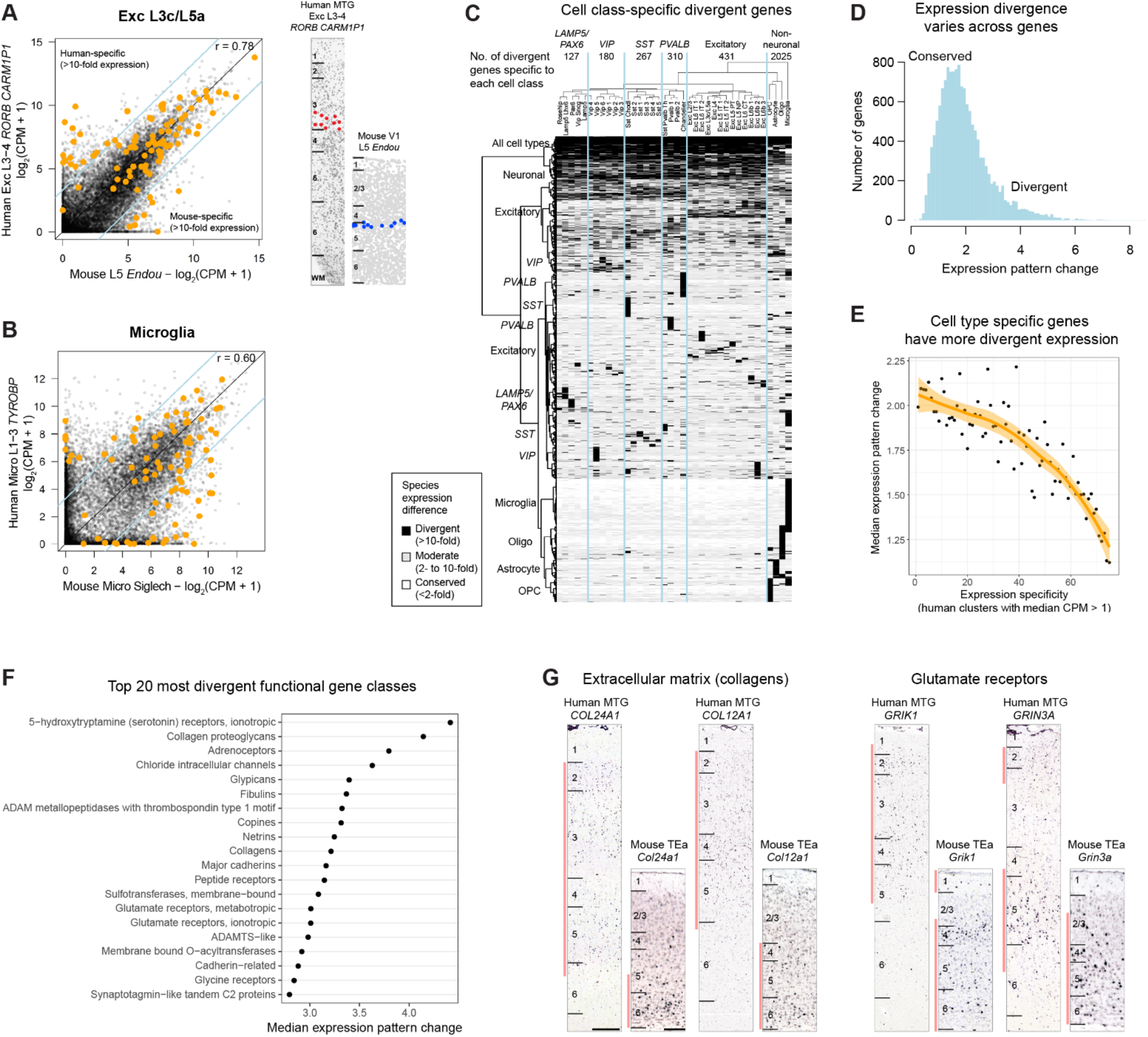
Divergent cell type expression between human and mouse. **(A)** Left: Comparison of expression levels of 14,414 orthologous genes between human and mouse for the most highly conserved one-to-one homologous type, Exc L3c/L5a. Genes outside of the blue lines have highly divergent expression (>10-fold change) between human and mouse. Approximately 100 genes (orange dots) are relatively specific markers in human and/or mouse. Right: ISH validation of layer distributions in human MTG and mouse primary visual cortex (data from Tasic et al., 2017). Cells are labeled based on expression of cluster marker genes in human (*RORB+/CNR1-/PRSS12+*) and mouse (*Scnn1a+/Hsd11b1*+). **(B)** Comparison of expression between human and mouse microglia, the least conserved homologous type. **(C)** Patterns of expression divergence between human and mouse for 8222 genes (57% of orthologous genes) with at least 10-fold expression change in one or more homologous cell types. Genes were hierarchically clustered and groups of genes that have similar patterns of expression divergence are labeled by the affected cell class. Top row: number of genes with expression divergence restricted to each broad class of cell types. **(D)** For each gene, the expression pattern change was quantified by the beta score (see Methods) of the absolute log fold change in expression between human and mouse. Genes with divergent patterns have large expression changes among a subset of homologous cell types. Genes with conserved patterns have similar expression levels in human and mouse or have a similar expression level change in all types. Pattern changes are approximately log-normally distributed, and a minority of genes have highly divergent patterns. **(E)** Genes expressed in fewer human cell types tended to have greater evolutionary divergence than more ubiquitously expressed genes. A loess curve and standard error was fit to median expression pattern changes across genes binned by numbers of clusters with expression (median CPM > 1). **(F)** Gene families with the most divergent expression patterns (highest median pattern change) include neurotransmitter receptors, ion channels, and cell adhesion molecules. **(G)** Genes estimated to have highly divergent expression patterns have different laminar expression validated by ISH in human and mouse. Red bars highlight layers with enriched expression. Scale bars: human (250μm), mouse (100μm).

Over half of all genes analyzed (8222, or 57%) had highly divergent expression in at least one of the 38 homologous types, and many genes had divergent expression restricted to a specific cell type or broad class (Fig. 8C). Non-neuronal cell types had the most highly divergent expression including 2025 genes with >10-fold species difference, supporting increased evolutionary divergence of non-neuronal expression patterns between human and mouse brain described previously^22^.

Most genes had divergent expression in a subset of types rather than all types, and this resulted in a shift in the cell type specificity or patterning of genes. These expression pattern changes were quantified as the beta score of log-fold differences across cell types (**Methods, Supplementary Table 2**), and scores were approximately log-normally distributed with a long tail of highly divergent genes (Fig. 8D). Cell type marker genes tended to be less conserved than more commonly expressed genes (Fig. 8E). In many cases, the most defining markers for cell types were not shared between human and mouse. For example, chandelier interneurons selectively express *Vipr2* in mouse but *COL15A1* and *NOG* in human (Fig. 4H). Interestingly, the functional classes of genes that best differentiate cell types within a species (Fig. 6A) are the same functional classes that show the most divergent expression patterns between species (Fig. 8F). In other words, the same gene families show cell type specificity in both species, but their patterning across cell types frequently differs.

The top 20 most divergent gene families between human and mouse (i.e. highest median pattern change) include neurotransmitter receptors (serotonin, adrenergic, glutamate, peptides, and glycine), ion channels (chloride), and cell adhesion molecules involved in axonal pathfinding (netrins and cadherins). Among the top 3% most divergent genes (see **Supplementary Table 2** for full list), the extracellular matrix collagens *COL24A1* and *COL12A1* and the glutamate receptor subunits *GRIK1* and *GRIN3A* were expressed in different cell types between species and were validated by ISH to have different laminar distributions in human MTG and mouse TEa (Fig. 8G). The cumulative effect of so many differences in the cellular patterning of genes with well characterized roles in neuronal signaling and connectivity is certain to cause many differences in human cortical circuit function.

## Discussion

Single cell transcriptomics provides a powerful tool to systematically characterize the cellular diversity of complex brain tissues, allowing a paradigm shift in neuroscience from the historical emphasis on cellular anatomy to a molecular classification of cell types and the genetic blueprints underlying the properties of each cell type. Echoing early anatomical studies^10^, recent studies of mouse neocortex have shown a great diversity of cell types^26,28^. Similar studies of human cortex^35,31,32^ have shown the same broad classes of cells but much less subtype diversity (Extended Data Fig. 4), likely resulting from technical differences, such as fewer nuclei sampled or reduced gene detection. A recent study showed a high degree of cellular diversity in human cortical layer 1^19^ by densely sampling high-quality postmortem human tissue with snRNA-seq and including intronic sequence to capture signal in nuclear transcripts^33^. The current study takes a similar dense sampling approach by sequencing approximately 16,000 single nuclei spanning all cortical layers of MTG, and defines 75 cell types representing non-neuronal (6), excitatory (24) and inhibitory (45) neuronal types. Importantly, robust cell typing could be achieved despite the increased biological and technical variability between human individuals. Nuclei from postmortem and acute surgically resected samples clustered together, and all clusters described contained nuclei from multiple individuals. Importantly, the ability to use these methods to study the fine cellular architecture of the human brain and to identify homologous cell types based on gene expression allows inference of cellular phenotypes across species as well. In particular, since so much knowledge has been accumulated about the cellular makeup of rodent cortex based on transcriptomics, physiology, anatomy and connectivity, this approach immediately allows strong predictions about such features as well as others that are not currently possible to measure in human such as developmental origins and long-range projection targets.

This molecular paradigm can help unify the field and increase the cellular resolution of many studies but has several consequences and challenges. Unambiguous definition of transcriptomic cell types *in situ* typically requires the detection of two or more markers with multiplexed molecular methods, demonstrating the need to further develop spatial transcriptomics methods^63^. Developing consistent nomenclature will also be challenging, particularly when marker genes are not conserved across species. Establishing cell type homologies across species can generate hypotheses about conserved and divergent cell features, and facilitates the larger, open access efforts to profile single cells across the brain underway in mouse, monkey, and human through the BRAIN Initiative^24^ and the Human Cell Atlas^25^. The current data are made publicly available with two new viewer applications to mine expression data across transcriptomic cell types in both human and mouse cortex (www.brain-map.org; viewer.cytosplore.org).

Interestingly, whereas excitatory neuron types are traditionally referred to as being confined to a single cortical layer, we find instead that many transcriptomically-defined excitatory types are represented in multiple layers. In part, this may reflect indistinct laminar boundaries in MTG; for example, von Economo^40^ noted intermixing of granule and pyramidal neurons in layer 4 along with blending of layer 4 pyramidal neurons into adjacent layers 3 and 5 in MTG. However, we find several types with broad spatial distributions across multiple layer boundaries, suggesting that indistinct laminar boundaries do not fully account for this lack of strict laminar segregation. Examination of the spatial distribution of excitatory neuron types in additional cortical areas will be necessary to determine if this is a particular feature of MTG or a more widespread phenomenon in human cortex.

The transcriptomic cellular organization and diversity in human MTG are surprisingly similar to those of mouse V1^26^, despite many differences in these data sets. First, mouse scRNA-seq was compared to human snRNA-seq, and to mitigate this, expression levels were estimated using intronic sequence that should be almost exclusively retained in the nucleus^33^. Second, young adult (~8-week-old) mice were compared to older (24-66 years) human specimens; however, prior transcriptomic studies demonstrated stable gene expression throughout adulthood in human^64,65^. Third, MTG in human was compared to V1 in mouse. This areal difference is expected to primarily affect comparison of excitatory neurons that vary more between regions than inhibitory neurons or glia^26^. Finally, scRNA-seq introduces significant biases due to differential survival of cell types during dissociation, necessitating the use of Cre-lines to enrich for under-sampled and rare cell types in mouse cortex^26^. In contrast, we found that snRNA-seq provides more unbiased sampling and estimates of cell type proportions. Despite these differences, the human and mouse cell type taxonomies could be matched at high resolution and reveal a “canonical” cellular architecture that is conserved between cortical areas and species. Beyond similarities in overall diversity and hierarchical organization, 10 cell types could be unambiguously mapped one-to-one between species, and 28 additional subclasses could be mapped at a higher level in the taxonomic tree. One-to-one matches were highly distinctive cell types, including several non-neuronal and neuronal types, such as chandelier cells. Comparison of absolute numbers of types between studies is challenging, but no major classes have missing homologous types other than exceedingly rare types that were likely undersampled in human, such as Cajal-Retzius cells.

A striking feature of cortical evolution is the relative expansion of the supragranular layers involved in cortico-cortical communication^18^. Consistent with this expansion, we find increased diversity of excitatory neurons in layers 2-4 in human compared to mouse. Layers 2 and 3 are dominated by three major types, but the most common layer 2/3 type exhibits considerable transcriptomic heterogeneity in the form of gene expression gradients, which would be expected to correlate with other cellular phenotypes. We also find expanded diversity of excitatory types in deep layer 3, along with a surprising increase in diversity in human layer 4 compared to mouse.

We observed several other evolutionary changes in cell type proportions and diversity that substantially alter the human cortical microcircuit. The relative proportions of major classes of GABAergic interneurons vary between human MTG and mouse V1, with human MTG having fewer *PVALB-* and *SST*-expressing interneurons and more *LAMP5/PAX6-* and VIP-expressing interneurons. Since these interneuron classes are derived from the MGE and CGE, respectively, in mouse, this difference is consistent with increased generation of CGE-derived interneurons in human^45^. Another major species difference is seen for human layer 5 excitatory neurons that are homologous to mouse sub-cortically projecting (PT) neurons. Both the frequency (<1% in human versus approximately 20% in mouse) and diversity (1 type in human versus 5 types in mouse)^26^ of PT neurons are markedly reduced in human, although reduced diversity may be an artifact of limited sampling in human. The sparsity of this type was confirmed *in situ* and was not a technical artifact of tissue processing. Rather, this sparsity likely reflects the 1200-fold expansion of human cortex relative to mouse compared to only 60-fold expansion of sub-cortical regions that are targets of these neurons^4,5^. If the number of PT neurons scales with the number of their sub-cortical projection targets, then the 20-fold greater expansion of cortical neurons would lead to a 20-fold dilution of PT neuron frequency as we observed. Indeed, the number of human corticospinal neurons, a subset of sub-cortically projecting neurons, has scaled linearly with the number of target neurons in the spinal cord, both increasing 40-fold compared to mouse^66,67,68^. Thus, this striking difference in cell type frequency may be a natural consequence of allometric scaling of the mammalian brain^69^.

Our results demonstrate striking species divergence of gene expression between homologous cell types, as observed in prior studies at the single gene^20^ or gross structural level^21^. We find more than half of all orthologous genes show a major (>10-fold) difference in expression in at least one of the 38 consensus cell types, and up to 20% of genes in any given cell type showing such major divergent expression. Several cell types, including endothelial cells, had such substantial expression divergence that they could not be matched across species using the methods employed here. These gene expression differences are likely to be functionally relevant, as divergent genes are associated with neuronal connectivity and signaling, signaling, including axon guidance genes, ion channels, and neuropeptide signaling. Surprisingly, serotonin receptors are the most divergent gene family, challenging the use of mouse models for the many neuropsychiatric disorders involving serotonin signaling^70^. Finally, the more selectively expressed a gene is in one species the less likely its pattern is to be conserved, and many well-known markers of specific cell types do not have conserved patterns.

Homologous cell types can have highly divergent features in concert with divergent gene expression. Here, we show that the interlaminar astrocyte, which has dramatic morphological specialization in primates including human, corresponds to one of two transcriptomic astrocyte types. A recent scRNA-seq analysis of mouse cortex also found 2 types, with one enriched in layer 1^28^. However, this mouse astrocyte type had less complex morphology and did not extend the long-range processes characteristic of interlaminar astrocytes. Thus, a 10-fold increase in size, the formation of a long process, and other phenotypic differences^17,55,54^ are evolutionary variations on a conserved genetically defined cell type. Similarly, a recent study identified the rosehip interneuron in human layer 1 19, which showed species differences in anatomy, physiology and marker gene profiles suggesting that it is a novel type of interneuron in human cortex. In fact, we now find that this rosehip type can be mapped to a mouse neurogliaform interneuron type. Thus, phenotypic differences large enough to define cell types with conventional criteria represent relatively minor variation on a conserved genetic blueprint for neurons as well.

Together these observations quantitatively frame the debate of whether human cortex is different from that of other mammals^9,10^, revealing a basic transcriptomic similarity of cell types punctuated by differences in proportions and gene expression between species that could greatly influence microcircuit function. The current results help to resolve the seeming paradox of conserved structure across mammals but failures in the use of mouse for pre-clinical studies^71,70^, and they highlight the need to analyze the human brain in addition to model organisms. The magnitude of differences between human and mouse suggest that similar profiling of more closely related non-human primates will be necessary to study many aspects of human brain structure and function. The enhanced resolution afforded by these molecular technologies also has great promise for accelerating a mechanistic understanding of brain evolution and disease.

## Acknowledgements

We would like to thank the Tissue Procurement, Tissue Processing, and Facilities teams at the Allen Institute for Brain Science for assistance with the transport and processing of postmortem and neurosurgical brain specimens. We thank the Technology team at the Allen Institute for assistance with data management. We gratefully acknowledge our collaborators at local hospitals (Swedish Medical Center, Harborview Medical Center/UW Medicine, and University of Washington Medical Center) for help with the coordination of human neurosurgical tissue collections. We thank Joe Davis and the San Diego Medical Examiner’s Office for assistance with postmortem tissue donations. We acknowledge the Molecular Biology, Histology, and Imaging teams at the Allen Institute for Brain Science for performing chromogenic *in situ* hybridization experiments. This work was funded by the Allen Institute for Brain Science, and by US National Institutes of Health grant 5 U01 MH1M812-02 to E.S.L. Funding from NWO-AES projects 12721: ‘Genes in Space’ and 12720: ‘VANPIRE’ (P.I. Anna Vilanova) for development of the Cytosplore MTG Viewer is gratefully acknowledged. We thank Baldur van Lew for scripting and narration of Cytosplore instructional and use case videos. The authors thank the Allen Institute founder, Paul G. Allen, for his vision, encouragement, and support.

## Author Contributions

E.S.L conceptualized and supervised the study. E.S.L. and R.Y. conceptualized the Human Cell Types Program. R.D.H and T.E.B. designed experiments. R.D.H., E.R.B., B. Long., J.L.C., B.P.L., S.I.S., K.B, J.G., D.H., S.L.D., M.M., S.P., E.R.T, N.V.S., and Z.M. contributed to nuclei isolation and/or validation experiments. T.E.B., J.A.M., O.P., Z.Y., O.F., J.G., S.S., and M.H. contributed to computational analyses. K.A.S. and B.T. managed the single-nucleus RNA-seq pipeline. L.T.G. developed data visualization tools. B.T. and H.Z. provided the mouse cortex transcriptomic cell type taxonomy for the cross-species comparative study. D.B., K.L., C.R, and M.T. performed single-nucleus RNA-seq. A. Bernard and J.P. managed establishment of single nucleus RNA-seq pipeline. A. Bernard and M.M contributed to the development and management of histological methods and data generation. K.B. performed immunohistochemistry experiments. R.D., N.D., T.C., J.N., A.O. processed postmortem brain tissues. A. Bernard and N.D. managed acquisition of postmortem and neurosurgical tissues. A. Beller, C.D.K, C.C., R.G.E., R.P.G., A.L.K, and J.G.O. contributed to neurosurgical tissue collections. B.A., M.K., and R.H.S. developed the semantic representation of clusters. J.E., T.H., A.M., and B. Lelieveldt developed the Cytosplore MTG Viewer. L.T.G., J.A.M., D.F., L.N, and A. Bernard contributed to the development of the RNA-Seq Data Navigator. S.R., A.S., and S.M.S. provided program management and/or regulatory compliance support. C.K. and A.R.J. provided institutional support and project oversight. E.S.L. and H.Z. directed the Allen Institute Cell Types Program. R.D.H., T.E.B., and E.S.L. wrote the paper with contributions from J.A.M and J.L.C., and in consultation with all authors.

## Methods

### Post-mortem tissue donors

Males and females 18 – 68 years of age with no known history of neuropsychiatric or neurological conditions (‘control’ cases) were considered for inclusion in this study (Extended Data Table 1). De-identified postmortem human brain tissue was collected after obtaining permission from decedent next-of-kin. The Western Institutional Review Board (WIRB) reviewed the use of de-identified postmortem brain tissue for research purposes and determined that, in accordance with federal regulation 45 CFR 46 and associated guidance, the use of and generation of data from de-identified specimens from deceased individuals did not constitute human subjects research requiring IRB review. Postmortem tissue collection was performed in accordance with the provisions of the Uniform Anatomical Gift Act described in Health and Safety Code §§ 7150, et seq., and other applicable state and federal laws and regulations. Routine serological screening for infectious disease (HIV, Hepatitis B, and Hepatitis C) was conducted using donor blood samples and only donors negative for all three tests were considered for inclusion in the study. Tissue RNA quality was assessed using an Agilent Bioanalyzer-generated RNA Integrity Number (RIN) and Agilent Bioanalyzer electropherograms for 18S/28S ratios. Specimens with RIN values ≥7.0 were considered for inclusion in the study (Extended Data Table 1).

### Processing of whole brain postmortem specimens

Whole postmortem brain specimens were transported to the Allen Institute on ice. Standard processing of whole brain specimens involved bisecting the brain through the midline and embedding of individual hemispheres in Cavex Impressional Alginate for slabbing. Coronal brain slabs were cut at 1cm intervals through each hemisphere and individual slabs were frozen in a slurry of dry ice and isopentane. Slabs were then vacuum sealed and stored at −80°C until the time of further use.

Middle temporal gyrus (MTG) was identified on and removed from frozen slabs of interest, and subdivided into smaller blocks for further sectioning. Individual tissue blocks were processed by thawing in PBS supplemented with 10mM DL-Dithiothreitol (DTT, Sigma Aldrich), mounting on a vibratome (Leica), and sectioning at 500μm in the coronal plane. Sections were placed in fluorescent Nissl staining solution (Neurotrace 500/525, ThermoFisher Scientific) prepared in PBS with 10mM DTT and 0.5% RNasin Plus RNase inhibitor (Promega) and stained for 5 min on ice. After staining, sections were visualized on a fluorescence dissecting microscope (Leica) and cortical layers were individually microdissected using a needle blade micro-knife (Fine Science Tools).

### Neurosurgical tissue donors

Tissue procurement from neurosurgical donors was performed outside of the supervision of the Allen Institute at local hospitals, and tissue was provided to the Allen Institute under the authority of the IRB of each participating hospital. A hospital-appointed case coordinator obtained informed consent from donors prior to surgery. Tissue specimens were de-identified prior to receipt by Allen Institute personnel. The specimens collected for this study were apparently non-pathological tissues removed during the normal course of surgery to access underlying pathological tissues. Tissue specimens collected were determined to be nonessential for diagnostic purposes by medical staff and would have otherwise been discarded.

### Processing of neurosurgical tissue samples

Neurosurgical tissue was transported to the Allen Institute in chilled, oxygenated artificial cerebrospinal fluid (ACSF) consisting of the following: 0.5 mM calcium chloride (dehydrate), 25 mM D-glucose, 20 mM HEPES, 10 mM magnesium sulfate, 1.2 mM sodium phosphate monobasic monohydrate, 92 mM N-methyl-d-glucamine chloride (NMDG-Cl), 2.5 mM potassium chloride, 30 mM sodium bicarbonate, 5 mM sodium L-ascorbate, 3 mM sodium pyruvate, and 2 mM thiourea. The osmolality of the solution was 295-305 mOsm/kg and the pH was 7.3. Slices were prepared using a Compresstome VF-200 or VF-300 vibratome (Precisionary Instruments). After sectioning, slices were recovered in ACSF containing 2 mM calcium chloride (dehydrate), 25 mM D-glucose, 20 mM HEPES, 2 mM magnesium sulfate, 1.2 mM sodium phosphate monobasic monohydrate, 2.5 mM potassium chloride, 30 mM sodium bicarbonate, 92 mM sodium chloride, 5 mM sodium L-ascorbate, 3 mM sodium pyruvate, and 2 mM thiourea at room temperature for at least 1 hour. After the recovery period, slices were transferred to RNase-free microcentrifuge tubes, snap frozen, and stored at −80°C until the time of use. Microdissection of cortical layers was carried out on tissue slices that were thawed and stained as described above for postmortem tissue.

### Nucleus sampling plan

We estimated that 16 cells were required to reliably discriminate two closely related Sst+ interneuron types reported by^27^. Monte Carlo simulations were used to estimate the sampling depth *N* needed to be 95% confident that at least 16 nuclei of frequency *f* have been selected from the population. Calculating *N* for a range of *f* revealed a simple linear approximation: *N* = 28 / *f*. Subtypes of mouse cortical layer 5 projection neurons can be rarer than 1% of the population^72^, so we targeted neuron types as rare as 0.2% of all cortical neurons. We initially sampled 14,000 neuronal nuclei distributed across cortical layers relative to the proportion of neurons reported in each layer^36^. We sampled approximately 1000 additional neuronal nuclei from layers with increased diversity observed based on RNA-seq data. We also targeted 1500 (10%) non-neuronal (NeuN-) nuclei and obtained approximately 1000 nuclei that passed QC, and we expected to capture types as rare as 3% of the non-neuronal population.

### Nucleus isolation and sorting

Microdissected tissue pieces were placed in into nuclei isolation medium containing 10mM Tris pH 8.0 (Ambion), 250mM sucrose, 25mM KCl (Ambion), 5mM MgCl2 (Ambion) 0.1% Triton-X 100 (Sigma Aldrich), 1% RNasin Plus, 1X protease inhibitor (Promega), and 0.1 mM DTT in 1ml dounce homogenizer (Wheaton). Tissue was homogenized using 10 strokes of the loose dounce pestle followed by 10 strokes of the tight pestle and the resulting homogenate was passed through 30μm cell strainer (Miltenyi Biotech) and centrifuged at 900xg for 10 min to pellet nuclei. Nuclei were resuspended in buffer containing 1X PBS (Ambion), 0.8% nuclease-free BSA (Omni-Pur, EMD Millipore), and 0.5% RNasin Plus. Mouse anti-NeuN conjugated to PE (EMD Millipore) was added to preparations at a dilution of 1:500 and samples were incubated for 30 min at 4°C. Control samples were incubated with mouse IgG1,k-PE Isotype control (BD Pharmingen). Samples were then centrifuged for 5 min at 400xg to pellet nuclei and pellets were resuspended in 1X PBS, 0.8% BSA, and 0.5% RNasin Plus. DAPI (4’, 6-diamidino-2-phenylindole, ThermoFisher Scientific) was applied to nuclei samples at a concentration of 0.1μg/ml.

Single nucleus sorting was carried out on either a BD FACSAria II SORP or BD FACSAria Fusion instrument (BD Biosciences) using a 130μm nozzle. A standard gating strategy was applied to all samples. First, nuclei were gated on their size and scatter properties and then on DAPI signal. Doublet discrimination gates were used to exclude nuclei aggregates. Lastly, nuclei were gated on NeuN signal (PE). Ten percent of nuclei were intentionally sorted as NeuN-negative and the remaining 90% of nuclei were NeuN-positive. Single nuclei were sorted into 8-well strip tubes containing 11.5μl of SMART-seq v4 collection buffer (Takara) supplemented with ERCC MIX1 spike-in synthetic RNAs at a final dilution of 1×10^-8^ (Ambion). Strip tubes containing sorted nuclei were briefly centrifuged and stored at −80°C until the time of further processing. Index sorting was carried out for most samples to allow properties of nuclei detected during sorting to be connected with the cell type identity revealed by subsequent snRNA-seq.

### RNA-sequencing

We used the SMART-Seq v4 Ultra Low Input RNA Kit for Sequencing (Takara #634894) per the manufacturer’s instructions for reverse transcription of RNA and subsequent cDNA amplification. Standard controls were processed alongside each batch of experimental samples. Control strips included: 2 wells without cells, 2 wells without cells or ERCCs (i.e. no template controls), and either 4 wells of 10 pg of Human Universal Reference Total RNA (Takara 636538) or 2 wells of 10 pg of Human Universal Reference and 2 wells of 10 pg Control RNA provided in the Clontech kit. cDNA was amplified with 21 PCR cycles after the reverse transcription step. AMPure XP Bead (Beckman Coulter A63881) purification was done using an Agilent Bravo NGS Option A instrument with a bead ratio of 1x, and purified cDNA was eluted in μl elution buffer provided by Takara. All samples were quantitated using PicoGreen®(ThermoFisher Scientific) on a Molecular Dynamics M2 SpectraMax instrument. cDNA libraries were examined on either an Agilent Bioanalyzer 2100 using High Sensitivity DNA chips or an Advanced Analytics Fragment Analyzer (96) using the High Sensitivity NGS Fragment Analysis Kit (1bp-6000bp). Purified cDNA was stored in 96-well plates at −20°C until library preparation.

The NexteraXT DNA Library Preparation (Illumina FC-131-1096) kit with NexteraXT Index Kit V2 Sets A-D (FC-131-2001, 2002, 2003, or 2004) was used for sequencing library preparation. NexteraXT DNA Library prep was done at either 0.5x volume manually or 0.4x volume on the Mantis instrument (Formulatrix). Three different cDNA input amounts were used in generating the libraries: 75pg, 100pg, and 125pg. AMPure XP bead purification was done using the Agilent Bravo NGS Option A instrument with a bead ratio of 0.9x and all samples were eluted in 22 μl of Resuspension Buffer (Illumina). Samples were quantitated using PicoGreen on a Molecular Bynamics M2 SpectraMax instrument. Sequencing libraries were assessed using either an Agilent Bioanalyzer 2100 with High Sensitivity DNA chips or an Advanced Analytics Fragment Analyzer with the High Sensitivity NGS Fragment Analysis Kit for sizing. Molarity was calculated for each sample using average size as reported by Bioanalyzer or Fragment Analyzer and pg/μl concentration as determined by PicoGreen. Samples were normalized to 2-10 nM with Nuclease-free Water (Ambion). Libraries were multiplexed at 96 samples/lane and sequenced on an Illumina HiSeq 2500 instrument using Illumina High Output V4 chemistry.

### RNA-seq gene expression quantification

Raw read (fastq) files were aligned to the GRCh38 human genome sequence (Genome Reference Consortium, 2011) with the RefSeq transcriptome version GRCh38.p2 (current as of 4/13/2015) and updated by removing duplicate Entrez gene entries from the gtf reference file for STAR processing. For alignment, Illumina sequencing adapters were clipped from the reads using the fastqMCF program^73^. After clipping, the paired-end reads were mapped using Spliced Transcripts Alignment to a Reference (STAR)^74^ using default settings. STAR uses and builds it own suffix array index which considerably accelerates the alignment step while improving on sensitivity and specificity, due to its identification of alternative splice junctions. Reads that did not map to the genome were then aligned to synthetic constructs (i.e. ERCC) sequences and the E. coli genome (version ASM584v2). The final results files included quantification of the mapped reads (raw exon and intron counts for the transcriptome-mapped reads). Also, part of the final results files are the percentages of reads mapped to the RefSeq transcriptome, to ERCC spike-in controls, and to E. coli. Quantification was performed using summerizeOverlaps from the R package GenomicAlignments^75^. Read alignments to the genome (exonic, intronic, and intergenic counts) were visualized as beeswarm plots using the R package *beeswarm*.

Expression levels were calculated as counts per million (CPM) of exonic plus intronic reads, and log2(CPM + 1) transformed values were used for a subset of analyses as described below.detection was calculated as the number of genes expressed in each sample with CPM > 0. CPM values reflected absolute transcript number and gene length, i.e. short and abundant transcripts may have the same apparent expression level as long but rarer transcripts. Intron retention varied across genes so no reliable estimates of effective gene lengths were available for expression normalization. Instead, absolute expression levels were estimated as fragments per kilobase per million (FPKM) using only exonic reads so that annotated transcript lengths could be used.

### Quality control of RNA-seq data

Nuclei were included for clustering analysis if they passed all of the following quality control (QC) thresholds:

- >30% cDNA longer than 400 base pairs
- >500,000 reads aligned to exonic or intronic sequence
- >40% of total reads aligned
- >50% unique reads
- TA nucleotide ratio > 0.7

After clustering (see below), clusters were identified as outliers if more than half of nuclei coexpressed markers of inhibitory (*GAD1, GAD2*) and excitatory (*SLC17A7*) neurons or were NeuN+ but did not express the pan-neuronal marker *SNAP25*. Median values of QC metrics listed above were calculated for each cluster and used to compute the median and inter-quartile range (IQR) of all cluster medians. Clusters were also identified as outliers if the cluster median QC metrics deviated by more than three times the IQRs from the median of all clusters.

Clusters were identified as donor-specific if they included fewer nuclei sampled from donors than expected by chance. For each cluster, the expected proportion of nuclei from each donor was calculated based on the laminar composition of the cluster and laminar sampling of the donor. For example, if 30% of layer 3 nuclei were sampled from a donor, then a layer 3-enriched cluster should contain approximately 30% of nuclei from this donor. In contrast, if only layer 5 were sampled from a donor, then the expected sampling from this donor for a layer 1-enriched cluster was zero. If the difference between the observed and expected sampling was greater than 50% of the number of nuclei in the cluster, then the cluster was flagged as donor-specific and excluded.

To confirm exclusion, clusters automatically flagged as outliers or donor-specific were manually inspected for expression of broad cell class marker genes, mitochondrial genes related to quality, and known activity-dependent genes.

### Clustering RNA-seq data

Nuclei and cells were grouped into transcriptomic cell types using an iterative clustering procedure based on community detection in a nearest neighbor graph as described in^33^. Briefly, intronic and exonic read counts were summed, and log2-transformed expression (CPM + 1) was centered and scaled across nuclei. X- and Y-chromosome were excluded to avoid nuclei clustering based on sex. Many mitochondrial genes had expression that was correlated with RNA-seq data quality, so nuclear and mitochondrial genes downloaded from Human MitoCarta2.0^76^ were excluded. Differentially expressed genes were selected while accounting for gene dropouts, and principal components analysis (PCA) was used to reduce dimensionality. Nearest-neighbor distances between nuclei were calculated using up to 20 principal components, Jaccard similarity coefficients were computed, and Louvain community detection was used to cluster this graph with 15 nearest neighbors. Marker genes were defined for all cluster pairs using two criteria: 1) significant differential expression (Benjamini-Hochberg false discovery rate < 0.05) using the R package *limma* and 2) either binary expression (CPM > 1 in >50% nuclei in one cluster and <10% in the second cluster) or >100-fold difference in expression. Pairs of clusters were merged if either cluster lacked at least one marker gene. Clustering was then applied iteratively to each sub-cluster until the occurrence of one of four stop criteria: 1) fewer than six nuclei (due to a minimum cluster size of three); 2) no significantly variable genes; 3) no significantly variable PCs; 4) no significant clusters.

To assess the robustness of clusters, the iterative clustering procedure described above was repeated 100 times for random subsets of 80% of nuclei. A co-clustering matrix was generated that represented the proportion of clustering iterations that each pair of nuclei were assigned to the same cluster. We defined consensus clusters by iteratively splitting the co-clustering matrix as described in^26^. We used the co-clustering matrix as the similarity matrix and clustered using either Louvain (>= 4000 nuclei) or Ward’s algorithm (< 4000 nuclei). We defined *Nk,l* as the average probabilities of nuclei within cluster *k* to co-cluster with nuclei within cluster l. We merged clusters *k* and *l* if *N*_k,l_ > max(*N*_k,k,_ N_l,l_) - 0.25 or if the sum of —log10(adjusted P-value) of differentially expressed genes between clusters *k* and *l* was less than 150. Finally, we refined cluster membership by reassigning each nucleus to the cluster to which it had maximal average co-clustering. We repeated this process until cluster membership converged.

Cluster names were defined using an automated strategy which combined molecular information (marker genes) and anatomical information (layer of dissection). Clusters were assigned a broad class of interneuron, excitatory neuron, microglia, astrocyte, oligodendrocyte precursor, oligodendrocyte, or endothelial cell based on maximal median cluster CPM of *GAD1, SLC17A7, TYROBP, AQP4, PDGFRA, OPALIN*, or *NOSTRIN*, respectively. Enriched layers were defined as the range of layers which contained at least 10% of the total cells from that cluster. Clusters in were then assigned a broad marker, defined by maximal median CPM of *PAX6, LAMP5, VIP, SST, PVALB, LINC00507, RORB, THEMIS, FEZF2, TYROBP, FGFR3, PDGFRA, OPALIN, or NOSTRIN*. Finally, clusters in all broad classes with more than one cluster (e.g., interneuron, excitatory neuron, and astrocyte) were assigned a gene showing the most specific expression in that cluster. These marker genes had the greatest difference in the proportion of expression (CPM > 1) with a cluster compared to all other clusters regardless of mean expression level.

### Scoring cluster marker genes

Many genes were expressed in the majority of nuclei in a subset of clusters. A marker score (beta) was defined for all genes to measure how binary expression was among clusters, independent of the number of clusters labeled (**Supplementary Table 2**). First, the proportion (*x_i_*) of nuclei in each cluster that expressed a gene above background level (CPM > 1) was calculated. Then, scores were defined as the squared differences in proportions normalized by the sum of absolute differences plus a small constant (ε) to avoid division by zero. Scores ranged from 0 to 1, and a perfectly binary marker had a score equal to 1.

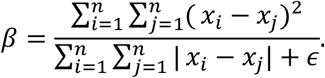

### Assigning core and intermediate cells

We defined core and intermediate cells as described^26^. Specifically, we used a nearest-centroid classifier, which assigns a cell to the cluster whose centroid has the highest Pearson’s correlation with the cell. Here, the cluster centroid is defined as the median expression of the 1200 marker genes with the highest beta score. To define core vs. intermediate cells, we performed 5-fold cross-validation 100 times. In each round, the cells were randomly partitioned into 5 groups, and cells in each group of 20% of the cells were classified by a nearest centroid classifier trained using the other 80% of the cells. A cell classified to the same cluster as its original cluster assignment more than 90 times was defined as a core cell, the others were designated intermediate cells. We define 14,204 core cells and 1,399 intermediate cells, which in most cases classify to only 2 clusters (1,345 out of 1,399, 96.1%). Most cells are defined as intermediate because they are confidently assigned to a different cluster from the one originally assigned (1,220 out of 1,399, 87.2%) rather than because they are not confidently assigned to any cluster.

### Cluster dendrograms

Clusters were arranged by transcriptomic similarity based on hierarchical clustering. First, the average expression level of the top 1200 marker genes (highest beta scores, as above) was calculated for each cluster. A correlation-based distance matrix 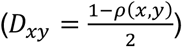 was calculated, and complete-linkage hierarchical clustering was performed using the “hclust” R function with default parameters. The resulting dendrogram branches were reordered to show inhibitory clusters followed by excitatory clusters, with larger clusters first, while retaining the tree structure. Note that this measure of cluster similarity is complementary to the co-clustering separation described above. For example, two clusters with similar gene expression patterns but a few binary marker genes may be close on the tree but highly distinct based on co-clustering.

### Mapping cell types to reported clusters

69 neuronal clusters in MTG were matched to 16 neuronal clusters reported by^31^ using nearest-centroid classifier of expression signatures. Specifically, single nucleus expression data was downloaded for 3042 single cells and 25,051 genes. 1359 marker genes (beta score > 0.4) of MTG clusters that had a matching gene in the Lake et al. dataset were selected, and the median expression for these genes was calculated for all MTG clusters. Next, Pearson’s correlations were calculated between each nucleus in the Lake et al. dataset and all 69 MTG clusters based on these 1359 genes. Nuclei were assigned to the cluster with the maximum correlation. A confusion matrix was generated to compare the cluster membership of nuclei reported by Lake et al. and assigned MTG cluster. The proportion of nuclei in each MTG cluster that were members of each of the 16 Lake et al. clusters were visualized as a dot plot with circle sizes proportional to frequency and colored by MTG cluster color.

### Colorimetric *in situ* hybridization

*In situ* hybridization (ISH) data for human and mouse cortex was from the Allen Human Brain Atlas and Allen Mouse Brain Atlas. All ISH data is publicly accessible at: www.brain-map.org. Data was generated using a semi-automated technology platform as described^77^, with modifications for postmortem human tissues as previously described^20^. Digoxigenin-labeled riboprobes were generated for each human gene such that they would have >50% overlap with the orthologous mouse gene in the Allen Mouse Brain Atlas^77^.

### GFAP immunohistochemistry

Tissue slices (350 μm) from neurosurgical specimens were fixed for 2-4 days in 4% paraformaldehyde in PBS at 4°C, washed in PBS, and cryoprotected in 30% sucrose. Cryoprotected slices were frozen and re-sectioned at 30 μm using a sliding microtome (Leica SM2000R). Free floating sections were mounted onto gelatin coated slides and dried overnight at 37 °C. Slides were washed in 1X tris buffered saline (TBS), followed by incubation in 3% hydrogen peroxide in 1X TBS. Slides were then heated in sodium citrate (pH 6.0) for 20 minutes at 98 °C. After cooling, slides were rinsed in MilliQ water followed by 1X TBS. Primary antibody (mouse anti-GFAP, EMD Millipore, #MAB360, clone GA5, 1:1500) was diluted in Renaissance Background Reducing Diluent (Biocare #PD905L). Slides were processed using a Biocare intelliPATH FLX Automated Slide Stainer. After primary antibody incubation, slides were incubated in Mouse Secondary Reagent (Biocare #IPSC5001G20), rinsed with 1X TBS, incubated in Universal HRP Tertiary Reagent (Biocare #IPT5002G20), rinsed in 1X TBS, and incubated in IP FLXDAB (Biocare Buffer #IPBF5009G20), and DAB chromogen (Biocare Chromogen #IPC5008G3). Slides were then rinsed in 1X TBS, incubated in DAB sparkle (Biocare #DSB830M), washed in MilliQ water, dehydrated through a series of graded alcohols, cleared with Formula 83, and coverslipped with DPX. Slides were imaged using an Aperio ScanScope XT slide scanner (Leica).

### Multiplex fluorescence *in situ* hybridization (FISH)

Genes were selected for multiplex FISH experiments that discriminated cell types and broader classes by visual inspection of differentially expressed genes that had relatively binary expression in the targeted types.

#### Single molecule FISH (smFISH)

Fresh-frozen human brain tissue from the MTG was sectioned at 10um onto Poly-L-lysine coated coverslips as described previously^78^, let dry for 10 min at room temperature, then fixed for 15 min at 4 C in 4% PFA. Sections were washed 3 × 10 min in PBS, then permeabilized and dehydrated with 100% isopropanol at room temperature for 3 min and allowed to dry. Sections were stored at −80 C until use. Frozen sections were rehydrated in 2XSSC (Sigma Aldrich 20XSSC, 15557036) for 5 min, then treated 2 × 5 min with 4%SDS (Sigma Aldrich, 724255) and 200mM boric acid (Sigma Aldrich, cat# B6768) pH 8.5 at room temperature. Sections were washed 3 times in 2X SSC, then once in TE pH 8 (Sigma Aldrich, 93283). Sections were heatshocked at 70 C for 10 min in TE pH 8, followed by 2XSSC wash at room temperature. Sections were then incubated in hybridization buffer (10% Formamide (v/v, Sigma Aldrich 4650), 10% Dextran Sulfate (w/v, Sigma Aldrich D8906), 200ug/mL BSA (Ambion AM2616), 2 mM Ribonucleoside vanadyl complex (New England Biolabs, S1402S), 1mg/ml tRNA (Sigma 10109541001) in 2XSSC) for 5 min at 38.5 C. Probes were diluted in hybridization buffer at a concentration of 250 nM and hybridized at 38.5 C for 2 h. Following hybridization, sections were washed 2 × 15 min at 38.5 C in wash buffer (2XSSC, 20% Formamide), and 1 × 15 min in wash buffer with 5 ug/ml DAPI (Sigma Aldrich, 32670). Sections are then imaged in Imaging buffer (20 mM Tris-HCl pH 8, 50 mM NaCl, 0.8% Glucose (Sigma Aldrich, G8270), 3 U/ml Glucose Oxidase (Sigma Aldrich, G2133), 90 U/ml Catalase (Sigma Aldrich, C3515). Following imaging, sections were incubated 3 × 10 min in stripping buffer (65% Formamide, 2X SSC) at 30 C to remove hybridization probes from the first round. Sections were then washed in 2X SSC for 3 × 5 min at room temperature prior to repeating the hybridization procedure.

#### RNAscope multiplex FISH

Human tissue specimens used for RNAscope multiplex FISH came from either neurosurgical resections or postmortem brain specimens. Mouse tissue for RNAscope experiments was from adult (P56 +/- 3 days) wildtype C57Bl/6J mice. All animal procedures were approved by the Institutional Animal Care and Use Committee at the Allen Institute for Brain Science (Protocol No. 1511). Mice were provided food and water ad libitum, maintained on a regular 12-h day/night cycle, and housed in cages with various enrichment materials added, including nesting materials, gnawing materials, and plastic shelters. Mice were anesthetized with 5% isoflurane and intracardially perfused with either 25 or 50 ml of ice cold, oxygenated artificial cerebral spinal fluid (0.5mM CaCl2, 25mM D-Glucose, 98mM HCl, 20mM HEPES, 10mM MgSO4, 1.25mM NaH2PO4, 3mM Myo-inositol, 12mM N-acetylcysteine, 96mM N-methyl-D-glucamine, 2.5mM KCl, 25mM NaHCO3, 5mM sodium L-Ascorbate, 3mM sodium pyruvate,0. 01mM Taurine, and 2mM Thiourea). The brain was then rapidly dissected, embedded in optimal cutting temperature (O.C.T.) medium, and frozen in a slurry of dry ice and ethanol. Tissues were stored at −80C until for later cryosectioning.

Fresh-frozen mouse or human tissues were sectioned at 14-16um onto Superfrost Plus glass slides (Fisher Scientific). Sections were dried for 20 minutes at −20C and then vacuum sealed and stored at −80C until use. The RNAscope multiplex fluorescent v1 kit was used according to the manufacturer’s instructions for fresh-frozen tissue sections (ACD Bio), with the following minor modifications: (1) fixation was performed for 60 minutes in 4% paraformaldehyde in 1X PBS at 4 °C, and (2) the protease treatment step was shortened to 15 min. Sections were imaged using either a 40X or 60X oil immersion lens on a Nikon TiE fluorescent microscope equipped with NIS-Elements Advanced Research imaging software (version 4.20).

#### RNAscope multiplex FISH with GFAP immunohistochemistry

Tissue sections were processed for RNAscope multiplex FISH detection of *ID3* (ACD Bio, #492181-C3, NM_002167.4) and *AQP4* (ACD Bio, #482441, NM_001650.5) exactly as described above. At the end of the RNAscope protocol, sections were fixed in 4% paraformaldehyde for 15 minutes at room temperature and then washed twice in 1X PBS for 5 minutes. Sections were incubated in blocking solution (10% normal donkey serum, 0.1% triton-x 100 in 1X PBS) for 30 minutes at room temperature and then incubated in primary antibody diluted 1:100 in blocking solution (mouse anti-GFAP, Sigma-Aldrich, #G3893, clone G-A-5) for 18 hours at 4C. Sections were then washed 3 times for 5 minutes each in 1X PBS, incubated with secondary antibody (goat anti-mouse IgG(H+L) Alexa Fluor 568 conjugate, ThermoFisher Scientific, #A-11004) for 30 minutes at room temperature, rinsed in 1X PBS 3 times for 5 minutes each, counterstained with DAPI (1ug/ml), and mounted with ProLong Gold mounting medium (ThermoFisher Scientific). Sections were imaged using either a 40X or 60X oil immersion lens on a Nikon TiE fluorescent microscope equipped with NIS-Elements Advanced Research imaging software (version 4.20).

### *In situ* validation of excitatory types

To validate excitatory neuron types, clusters were labeled with cell type specific combinatorial gene panels. For each gene panel, positive cells were manually called by visual assessment of RNA spots for each gene. The total number of positive cells was quantified for each section. Cells were counted on at least three sections derived from at least two donors for each probe combination. DAPI staining was used to determine the boundaries of cortical layers within each tissue section and the laminar position of each positive cell was recorded. The percentage of labeled cells per layer, expressed as a fraction of the total number of labeled cells summed across all layers, was calculated for each type. Probes used in these experiments were as follows (all from ACD Bio): *SLC17A7* (#415611, NM_020309.3), *RORB* (#446061, #446061-C2, NM_006914.3), *CNR1* (#591521-C2, NM_001160226.1), *PRSS12* (#493931-C3, NM_003619.3), *ALCAM* (#415731-C2, NM_001243283.1), *MET* (#431021, NM_001127500.1), *MME* (#410891-C2, NM_007289.2), *NTNG1* (#446101-C3, NM_001113226.1), *HS3ST4* (#506181, NM_006040.2), *CUX2* (#425581-C3, NM_015267.3), *PCP4* (#446111, NM_006198.2), *GRIN3A* (#534841-C3, NM_133445.2), *GRIK3* (#493981, NM_000831.3), *CRHR2* (#469621, NM_001883.4), *TPBG* (#405481, NM_006670.4), *POSTN* (#409181-C3, NM_006475.2), *SMYD1* (#493951-C2, NM_001330364.1)

### *In situ* validation of putative chandelier cells

Tissue sections were labeled with the gene panel *GAD1, PVALB*, and *NOG*, or *COL15A1*, specific markers of the Inh L2-5 *PVALB SCUBE3* putative chandelier cell cluster. Probes were as follows (all from ACD Bio): *GAD1* (#404031-C3, NM_000817.2), *PVALB* (#422181-C2, NM_002854.2), *NOG* (#416521, NM_005450.4), *COL15A1* (#484001, NM_001855.4). Counts were conducted on sections from 3 human tissue donors. For each donor, the total number of *GAD1+, PVALB+* and *NOG+* cells was summed across multiple sections. The laminar position of each cell, based on boundaries defined by assessing DAPI staining patterns in each tissue section, was recorded. The proportion of chandelier cells in each layer was calculated as a fraction of the total number of *GAD1+/PVALB+/NOG+* cells summed across all layers for each specimen.

### Cell counts of broad interneuron classes

Tissue sections were labeled with the RNAscope Multiplex Fluorescent kit (ACD Bio) as described above. For human tissue sections, the following probes (all from ACD Bio) were used: *GAD1* (#404031, NM_000817.2); *ADARB2* (#511651-C3, NM_018702.3); *LHX6* (#460051-C2, NM_014368.4). For mouse tissue sections, the following probes were used: *Gad1* (#400951, NM_008077.4); *Adarb2* (#519971-C3, NM_052977.5); *Lhx6* (#422791-C2, NM_001083127.1). The expression of each gene was assessed by manual examination of corresponding RNA spots. Cell counts were conducted on sections from 3 human tissue donors:2 neurosurgical and 1 postmortem. For mouse, 3 independent specimens were used. For both human and mouse, >500 total *GAD1+* cells per specimen were counted (Human, n=2706, 1553, and 3476 *GAD1+* cells per donor, respectively; Mouse, n=1897, 2587, and 708 *GAD1+* cells per specimen, resepectively). Expression of *ADARB2/Adarb2* and *LHX6/Lhx6* was manually assessed in each *GAD1+* cell and cells were scored as being positive or negative for each gene. At the same time, the laminar position of each *GAD1+* cell was recorded. Cell density,highlighted by DAPI staining, was used to determine laminar boundaries. The percentage of each cell class expressed as a fraction of total *GAD1+* cells and the percentage of each cell class per layer, expressed as a fraction of the total number of GAD1+ cells per layer, were calculated for each specimen. Statistical comparisons between human and mouse were done using unpaired two-tailed t-tests with Holm-Sidak correction for multiple comparisons.

### Imaging and quantification of smFISH expression

smFISH images were collected using an inverted microscope in an epifluorescence configuration (Zeiss Axio Observer.Z1) with a 63x oil immersion objective with numerical aperture 1.4. The sample was positioned in x, y and z with a motorized x, y stage with linear encoders and z piezo top-plate (Applied Scientific Instruments MS 2000-500) and z stacks with 300 nm plane spacing were collected in each color at each stage position through the entire z depth of the sample. Fluorescence emission was filtered using a high-speed filterwheel (Zeiss) directly below the dichroic turret and imaged onto a sCMOS camera (Hamamatsu ORCA Flash4.0) with a final pixel size of 100 nm. Images were collected after each round of hybridization using the same configuration of x, y tile locations, aligned manually before each acquisition based on DAPI fluorescence. smFISH signal was observed as diffraction-limited spots which were localized in 3D image stacks by finding local maxima after spatial bandpass filtering. These maxima were filtered for total intensity and radius to eliminate dim background and large, bright lipofuscin granules. Outlines of cells and cortical layers were manually annotated on images of GAD, SLC17A7 and DAPI as 2D polygons using FIJI. The number of mRNA molecules in each cell for each gene was then calculated and converted to densities (spots per 100um^2^).

Background expression of the excitatory neuron marker *SLC17A7* was defined as the 95th quantile of *SLC17A7* spot density among cells in cortical layer 1, since no excitatory cells should be present in layer 1. Excitatory neurons were defined as any cell with *SLC17A7* spot density greater than this threshold. To map excitatory cells to MTG reference clusters, spot counts were log-transformed and scaled so that the 90th quantile of expression for each gene in smFISH matched the maximum median cluster expression of that gene among the reference clusters. Reference clusters that could not be discriminated based on the smFISH panel of nine genes were merged and all comparisons between smFISH and RNA-seq cluster classes were performed using these cluster groups. Scaled spot densities for each cell were then compared to median expression levels of each reference cluster using Pearson correlation, and each cell was assigned to the cluster with the highest correlation. For cells that mapped to the Exc L2-3 *LINC00507 FREM3* cluster, *LAMP5* and *COL5A2* expression was plotted as a dot plot where the size and color of dots corresponded to probe spot density and the location corresponded to the *in situ* location.

### MetaNeighbor analysis

To compare the ability of different gene sets to distinguish cell types in mouse versus human cortex, we performed a modified supervised MetaNeighbor analysis^79^ independently for both species. First, we divided our data sets into two artificial experiments, selecting random groups of equal size up to a maximum of 10 cells per cluster for each experiment. We next ran MetaNeighbor separately for clusters from each broad class (GABAergic, glutamatergic, and non-neuronal) using the R function “run_MetaNeighbor” where “experiment_labels” are 1 or 2 corresponding to the two artificial experiments, “celltype_labels” are 2 for cells in the targeted cluster and 1 for cells in all other clusters of the same broad class, and “genesets” were all of the HGNC gene sets included in Table S3 of^52^. Mean AUROC scores for each gene set were then calculated by averaging the reported AUROC scores for a gene set across all clusters within a given broad class. This processes was repeated for 10 divisions of the human and mouse data into random experimental groups. Means and standard deviations of these mean AUROC scores for human and mouse GABAergic cell types are compared in Fig 5.

### Estimation of cell type homology

We aligned single nucleus and single cell RNA-seq data from human MTG and mouse primary visual cortex by applying canonical correlation analysis (CCA) as implemented in the Seurat R package^56^. We used log2-transformed CPM of intronic plus exonic reads for both datasets. Including exonic reads increased experimental differences due to measuring whole cell versus nuclear transcripts, but this was out-weighed by improved gene detection. We separated each of the datasets into three broad cell classes: GABAergic, glutamatergic, and non-neuronal, based on their assigned clusters, and selected up to 200 cells from each cluster. We included mouse non-neuronal cells from cell types that we had captured in our human survey, including astrocytes, oligodendrocyte precursors, oligodendrocytes, endothelial cells, and microglia. For each of these datasets, we selected the union of the top 2,000 genes with the highest dispersion for human and mouse and calculated 40 canonical correlates with diagonal CCA. Following this step, we removed 88 nuclei or cells for which the variance explained by CCA was less than half of the variance explained by PCA, and aligned the canonical basis vectors to allow integrated analysis. In particular, all human endothelial nuclei and over half of human microglial nuclei were removed along with mouse Cajal-Retzius cells.

We defined homologous cell types by clustering canonical correlates and identifying human and mouse samples that co-clustered. Initially, the first 10 canonical correlates were selected, and a weighted graph was constructed based on the Jaccard similarity of the 10 nearest neighbors of each sample. Louvain community detection was run to identify clusters that optimized the global modularity of the partitioned graph. For each pair of human and mouse clusters, the overlap was defined as the sum of the minimum proportion of samples in each cluster that overlapped within each CCA cluster. This approach identified pairs of human and mouse clusters that consistently co-clustered within one or more CCA clusters. Cluster overlaps varied from 0 to 1 and were visualized as a heatmap with human clusters as rows and mouse clusters as columns. Cell type homologies were identified as one-to-one, one-to-many, or many-to-many based on the pattern of overlap between clusters. A quality score was calculated for the homology mapping that rewarded overlaps greater than 0.6 (0.2 for non-neuronal clusters) and penalized for clusters lacking any overlaps. For each human cluster, the inverse of the sum of the number of overlapping mouse clusters was calculated, and this value was set to −1 if no overlapping clusters were found. The quality score was defined as the sum of the scores for the individual clusters and could range from −38 (no overlap) to 38 (all one-to-one matches). Including more canonical correlates or fewer nearest neighbors increased the number of cell types that could be discriminated within each species (increasing the quality score) but also resulted in more species-specific clusters (decreasing the quality score). A grid based search was used to select the number of canonical correlates and nearest neighbors that maximized the quality score. Detection of homologous cell types was confirmed by visual inspection (**Supplementary Table 1**).

### Quantification of expression divergence

For each pair of 38 homologous human and mouse cell types, the average expression of 14,414 orthologous genes was calculated as the average counts per million of intronic reads. Only intronic reads were used to better compare these single nucleus (human) and single cell (mouse) datasets. Average expression values were log2-transformed and scatter plots and Pearson’s correlations were calculated to compare human and mouse. Genes were ranked based on their cell type-specificity in human and mouse using a tau score defined in^80^, and the union of the top 50 markers in human and mouse were highlighted in the scatter plots. The fold difference in expression between human and mouse was calculated for all genes and homologous cell types and thresholded to identify large (>10-fold), moderate (2- to 10-fold), and small (<2-fold) differences. A heatmap was generated showing expression differences across cell types, and hierarchical clustering using Ward’s method was applied to group genes with similar patterns of expression change. For each of 6 major classes of cell types (*LAMP5/PAX6, VIP, SST, PVALB*, excitatory, non-neuronal), the number of genes was quantified that had >10-fold change in at least one cell type in that class and <10-fold change in all cell types in the other 5 classes. The expression pattern change of 14,414 genes was quantified as the beta score (see marker score methods above) of log2-expression differences across 38 homologous cell types (Supplementary Table 2). Genes with high scores have a large fold-change in expression in one or more (but not all) cell types. For each gene, the number of clusters with median expression (CPM) > 1 was compared to the median pattern change of those genes. A loess curve and standard error were fit using the R package *ggplot*. Finally, the median pattern change was calculated for the functional gene families used in the MetaNeighbor analysis described above.

### Data and Code Availability

Data and code used to produce figures will be available from https://github.com/AllenInstitute/MTG_celltypes. RNA-seq data from this study is publicly available and can be downloaded at http://celltypes.brain-map.org/, and data can be visualized and analyzed using two complementary viewers at http://celltypes.brain-map.org/rnaseq/human and https://viewer.cytosplore.org/.

---

**Extended Data Table 1.**
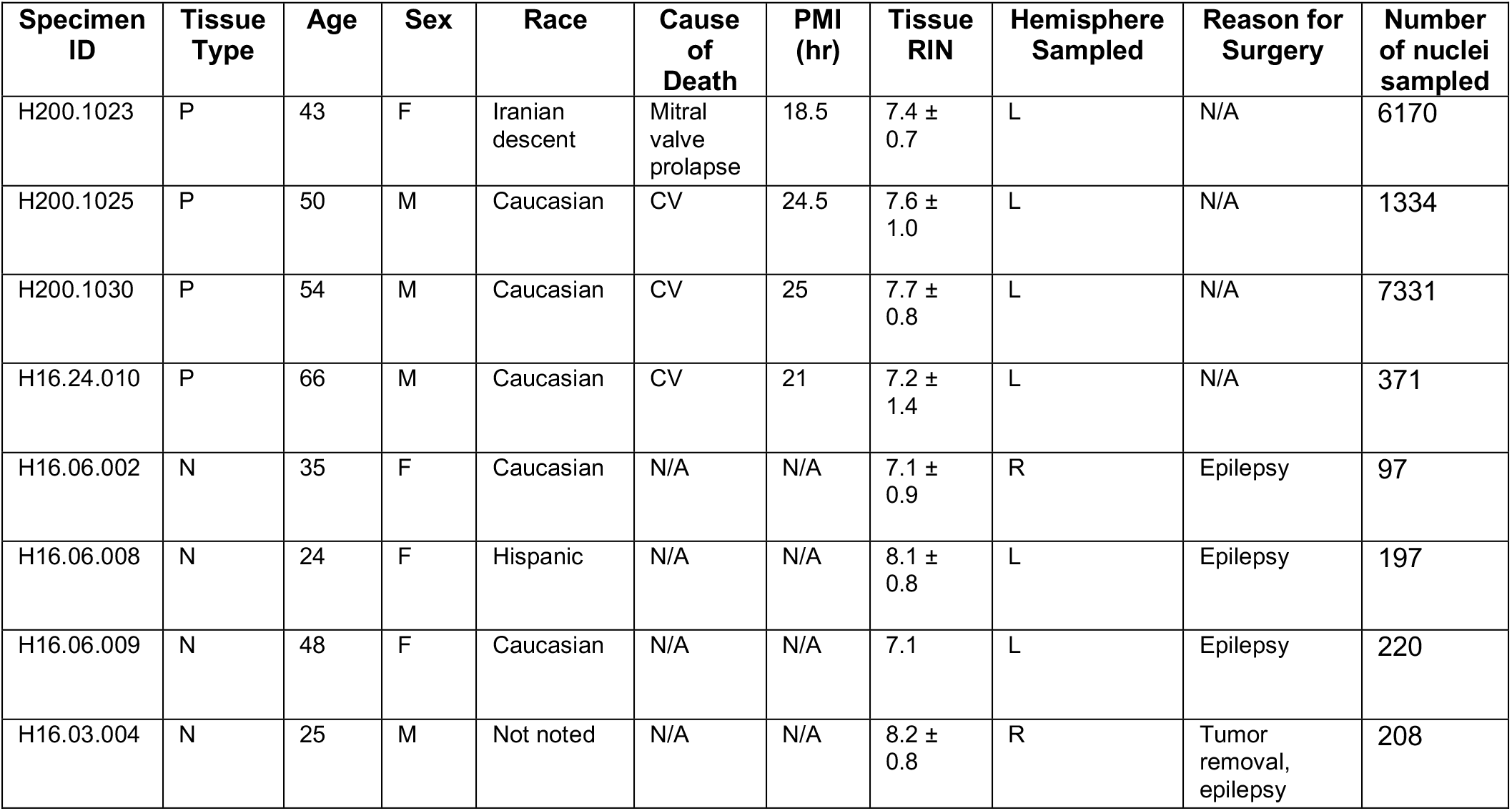
Summary of human tissue donor information. Tissue types - P, postmortem, N - neurosurgical. Cause of death - CV, cardiovascular, N/A, not applicable. PMI - postmortem interval. RIN - RNA Integrity Number.

| Specimen ID | Tissue Type | Age | Sex | Race | Cause of Death | PMI (hr) | Tissue RIN | Hemisphere Sampled | Reason for Surgery | Number of nuclei sampled |
| --- | --- | --- | --- | --- | --- | --- | --- | --- | --- | --- |
| H200.1023 | P | 43 | F | Iranian descent | Mitral valve prolapse | 18.5 | 7.4 ± 0.7 | L | N/A | 6170 |
| H200.1025 | P | 50 | M | Caucasian | CV | 24.5 | 7.6 ± 1.0 | L | N/A | 1334 |
| H200.1030 | P | 54 | M | Caucasian | CV | 25 | 7.7 ± 0.8 | L | N/A | 7331 |
| H16.24.010 | P | 66 | M | Caucasian | CV | 21 | 7.2 ± 1.4 | L | N/A | 371 |
| H16.06.002 | N | 35 | F | Caucasian | N/A | N/A | 7.1 ± 0.9 | R | Epilepsy | 97 |
| H16.06.008 | N | 24 | F | Hispanic | N/A | N/A | 8.1 ± 0.8 | L | Epilepsy | 197 |
| H16.06.009 | N | 48 | F | Caucasian | N/A | N/A | 7.1 | L | Epilepsy | 220 |
| H16.03.004 | N | 25 | M | Not noted | N/A | N/A | 8.2 ± 0.8 | R | Tumor removal, epilepsy | 208 |

**Extended Data Figure 1.**
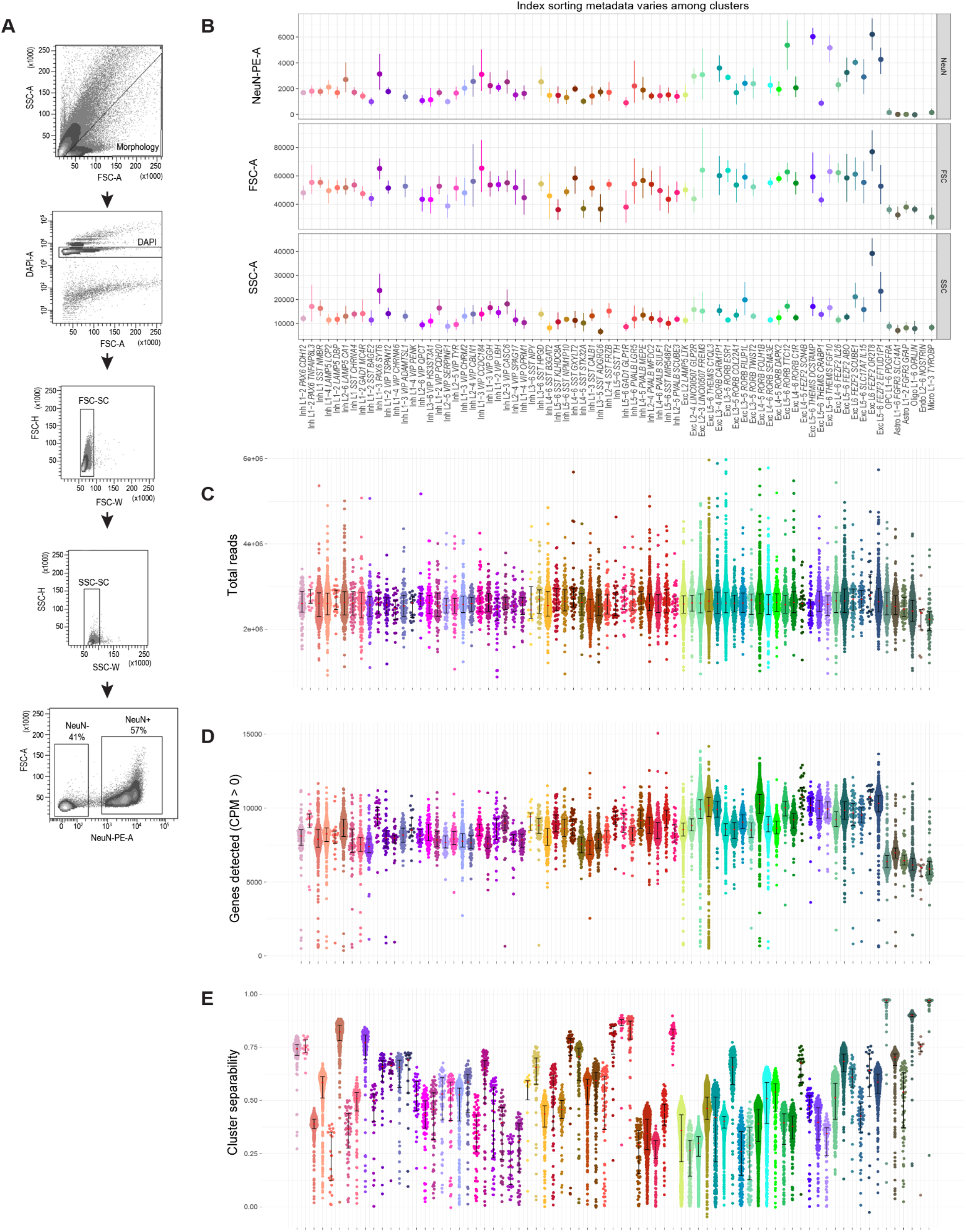
Nuclei metadata summarized by cluster. **(A)** FACS gating scheme for nuclei sorts. **(B)** FACS metadata for index sorted single nuclei shows significant variability in NeuN fluoresence intensity (NeuN-PE-A), size (forward-scatter area, FSC-A), and granularity (side-scatter area, SSC-A) across clusters. As expected, non-neuronal nuclei have almost no NeuN staining and are smaller (as inferred by lower FSC values). **(C-E)** Scatter plots plus median and interquartile interval of three QC metrics grouped and colored by cluster. **(C)** Median total reads were approximately 2.6 million for all cell types, although slightly lower for non-neuronal nuclei. **(D)** Median gene detection was highest among excitatory neuron types in layers 5 and 6 and a subset of types in layer 3, lower among inhibitory neuron types, and significantly lower among non-neuronal types. **(E)** Cluster separability varied substantially among cell types, with a subset of neuronal types and all non-neuronal types being highly discrete.

**Extended Data Figure 2.**
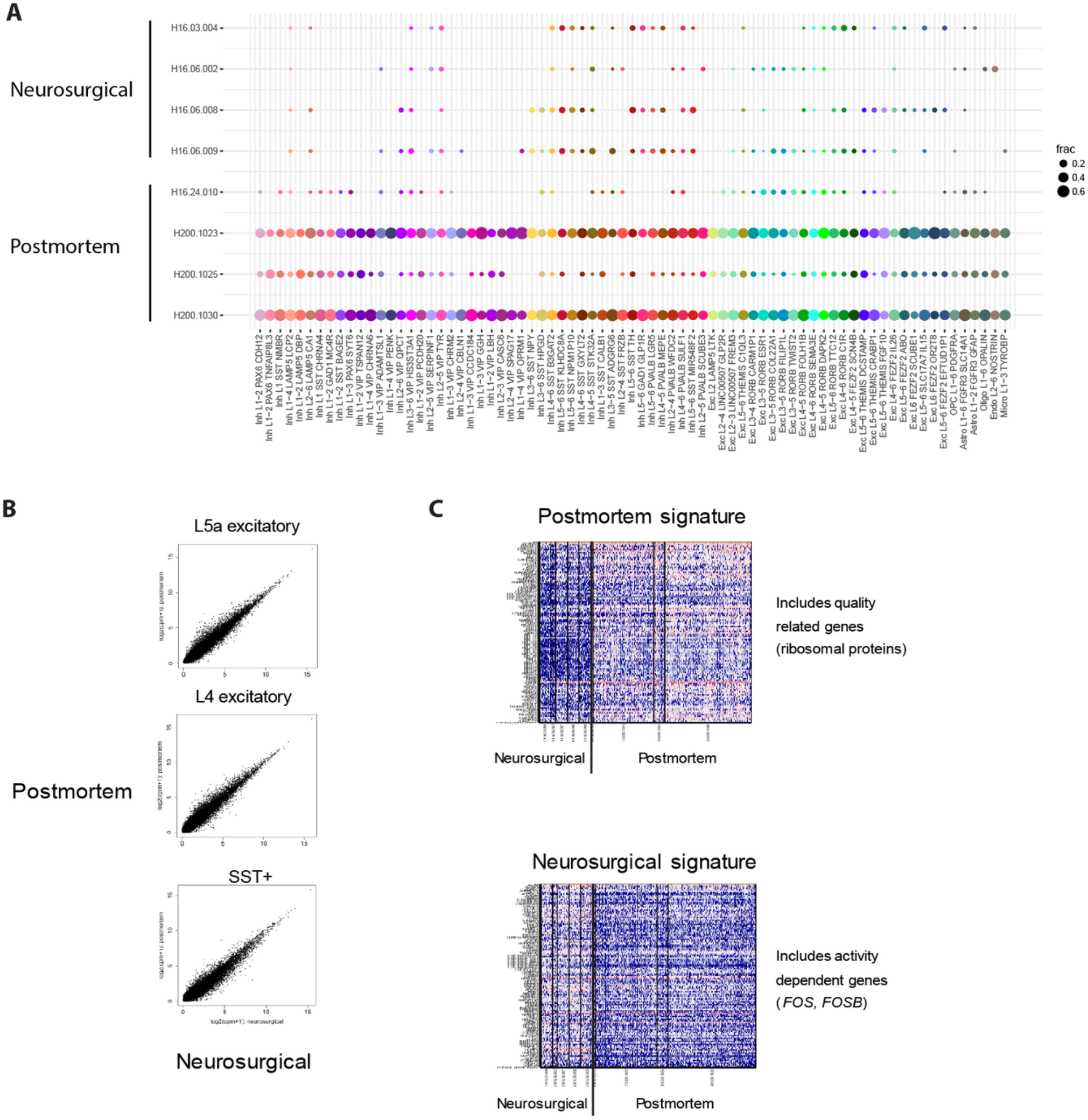
Small but consistent expression signature of donor tissue source. **(A)** Dot plot showing the proportion of nuclei isolated from neurosurgical and postmortem donors among human MTG clusters. Note that most nuclei from neurosurgical donors were isolated only from layer 5 so clusters enriched in other layers, such as layer 1 interneurons, have low representation of these donors. **(B)** Highly correlated expression between nuclei from postmortem and neurosurgical donors among two classes of excitatory neurons and one class of inhibitory neurons. Nuclei were pooled and compared within these broad classes due to the low sampling of individual clusters from neurosurgical donors. **(C)** Expression (log_10_(CPM + 1)) heatmaps of genes that are weakly but consistently up-regulated in nuclei from postmortem or neurosurgical donors including ribosomal genes and activity-dependent genes, respectively.

**Extended Data Figure 3.**
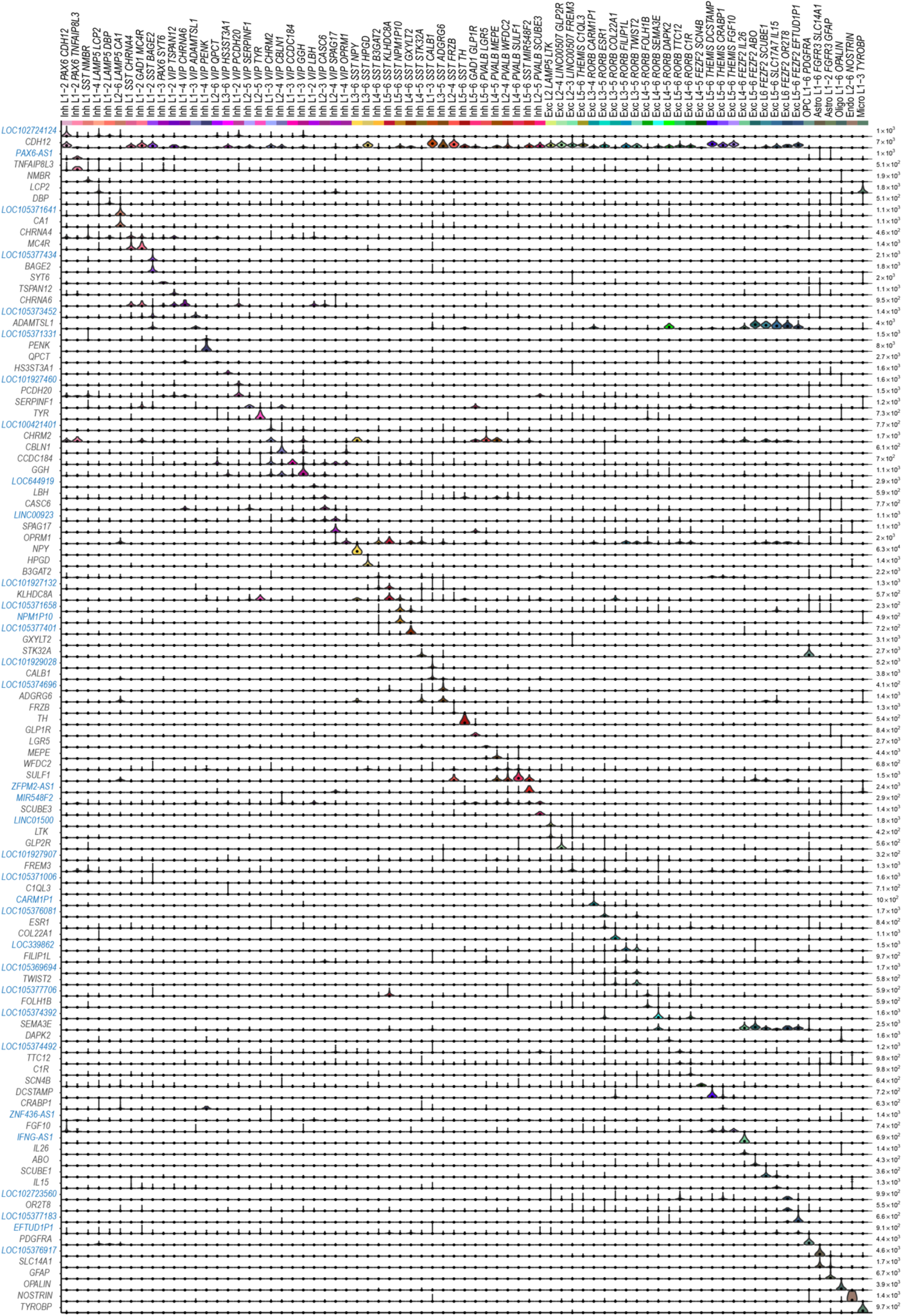
Expression of cell type specific markers. Violin plots of the best cell type markers include many non-coding genes (blue symbols): lncRNAs, antisense transcripts, and unnamed (LOC) genes. Expression values are on a linear scale and dots indicate median expression. Note that LOC genes were excluded from cluster names, and the best non-LOC marker genes were used instead.

**Extended Data Figure 4.**
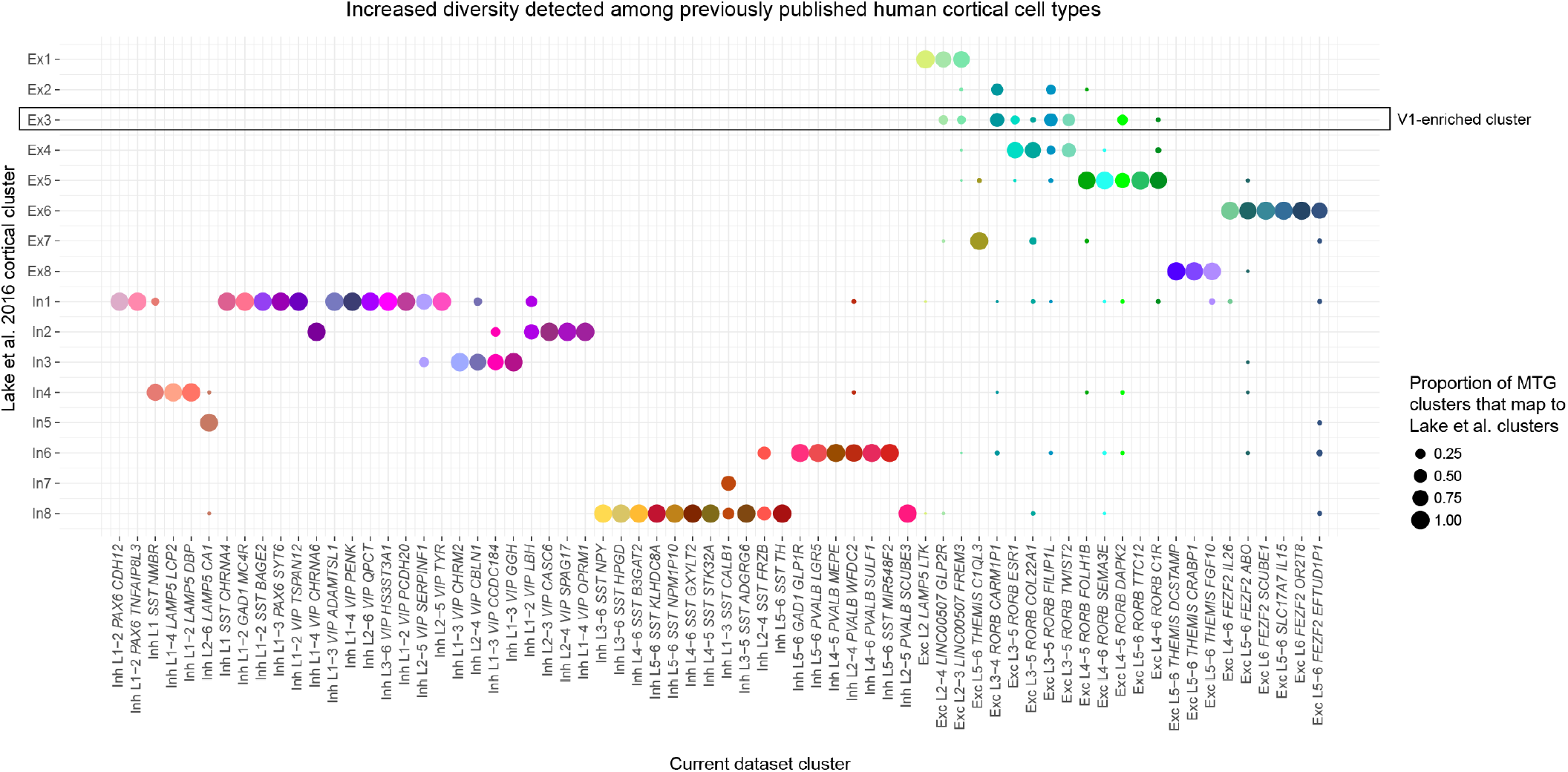
Matching MTG clusters to reported human cortical cell types. Dot plot showing the proportion of each MTG cluster that matches 16 clusters reported by^29^ based on a centroid expression classifier. Ex3 was highly enriched in visual cortex and not detected in temporal cortex by Lake et al.

**Extended Data Figure 5.**
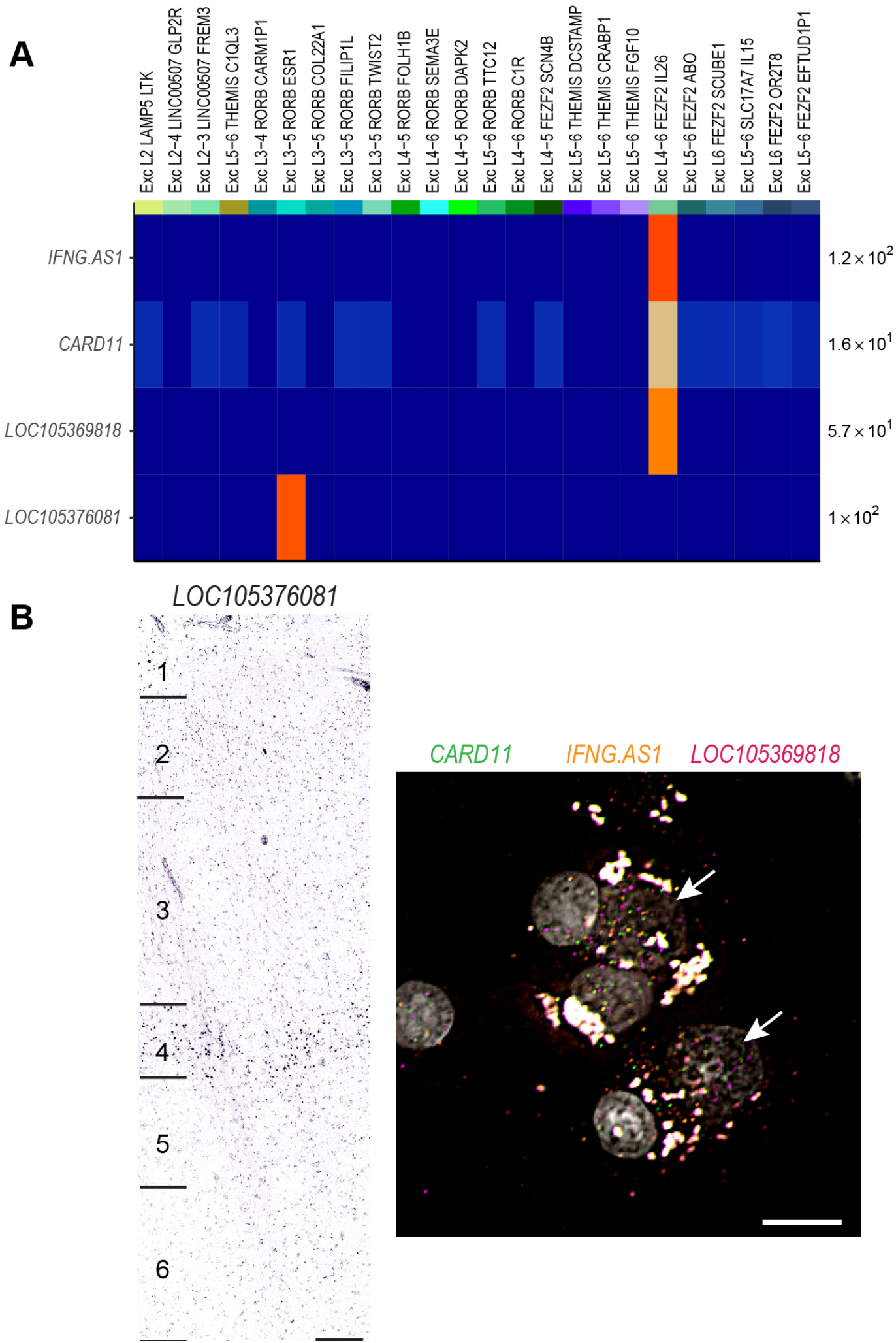
In situ validation of LOC and antisense transcripts as cell type specific markers. **(A)** Heatmap illustrating cell type specific expression of several LOC genes and one antisense transcript (iFNG-AS1). **(B)** Left - chromogenic in situ hybridization for LOC105376081, a specific marker of the Exc L3-5 *RORB ESR1* type shows expression of this gene predominantly in layer 4, consistent with the anatomical location of this cell type. Scale bar, 100um. Right - triple RNAscope FISH for markers of the Exc L4-6 *FEZF2 IL26* type. Coexpression of the protein coding gene *CARD11* with *IFNG-AS1*, an antisense transcript, and *LOC105369818* is apparent within several DAPI-labeled nuclei (white arrows). Scale bar, 15μm.

**Extended Data Figure 6.**
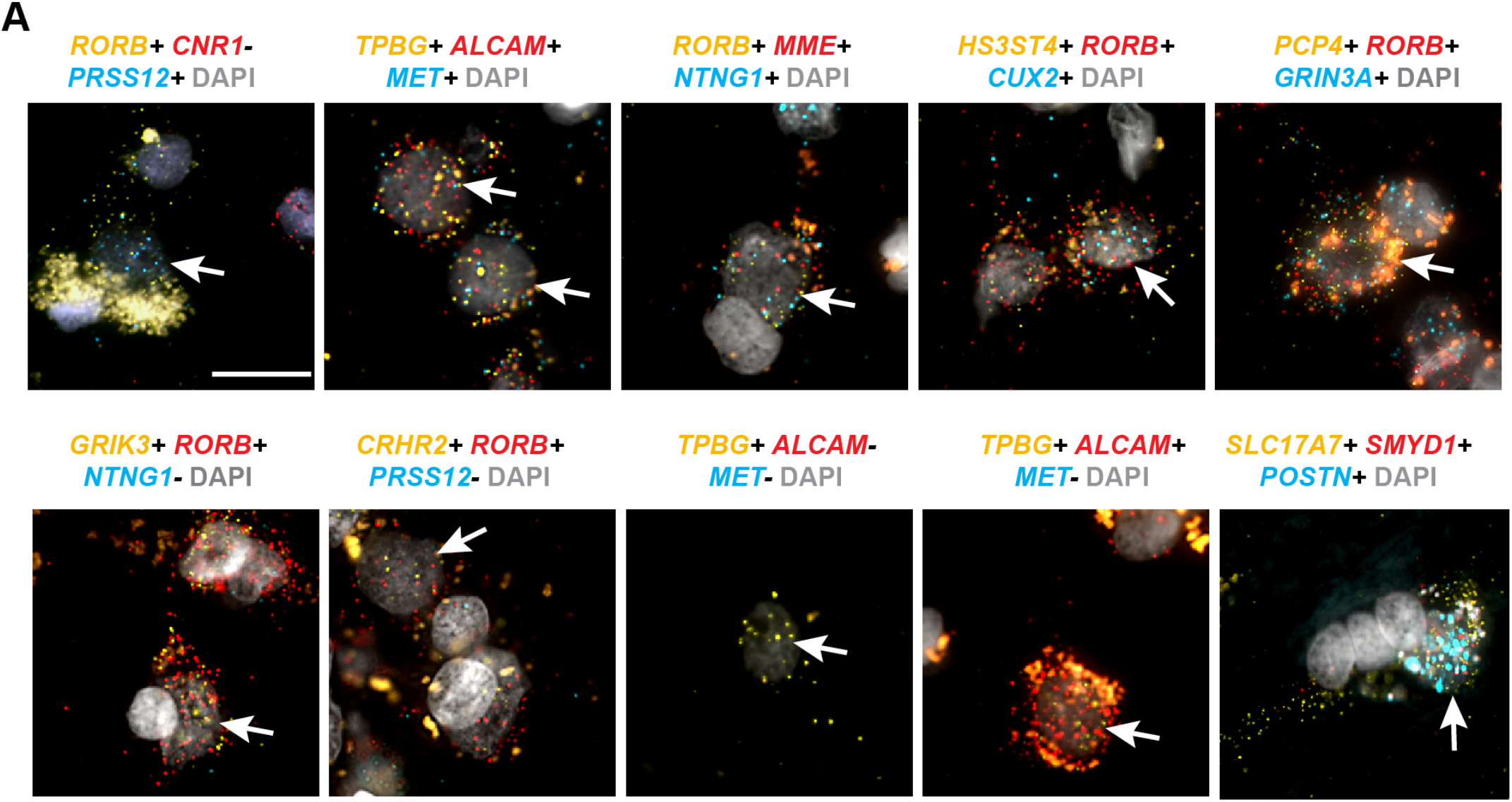
RNAscope multiplex FISH validation of 10 excitatory neuron types. Gene combinations probed are listed above each image. Labeled cells are indicated by white arrows. Scale bar, 20μm.

**Extended Data Figure 7.**
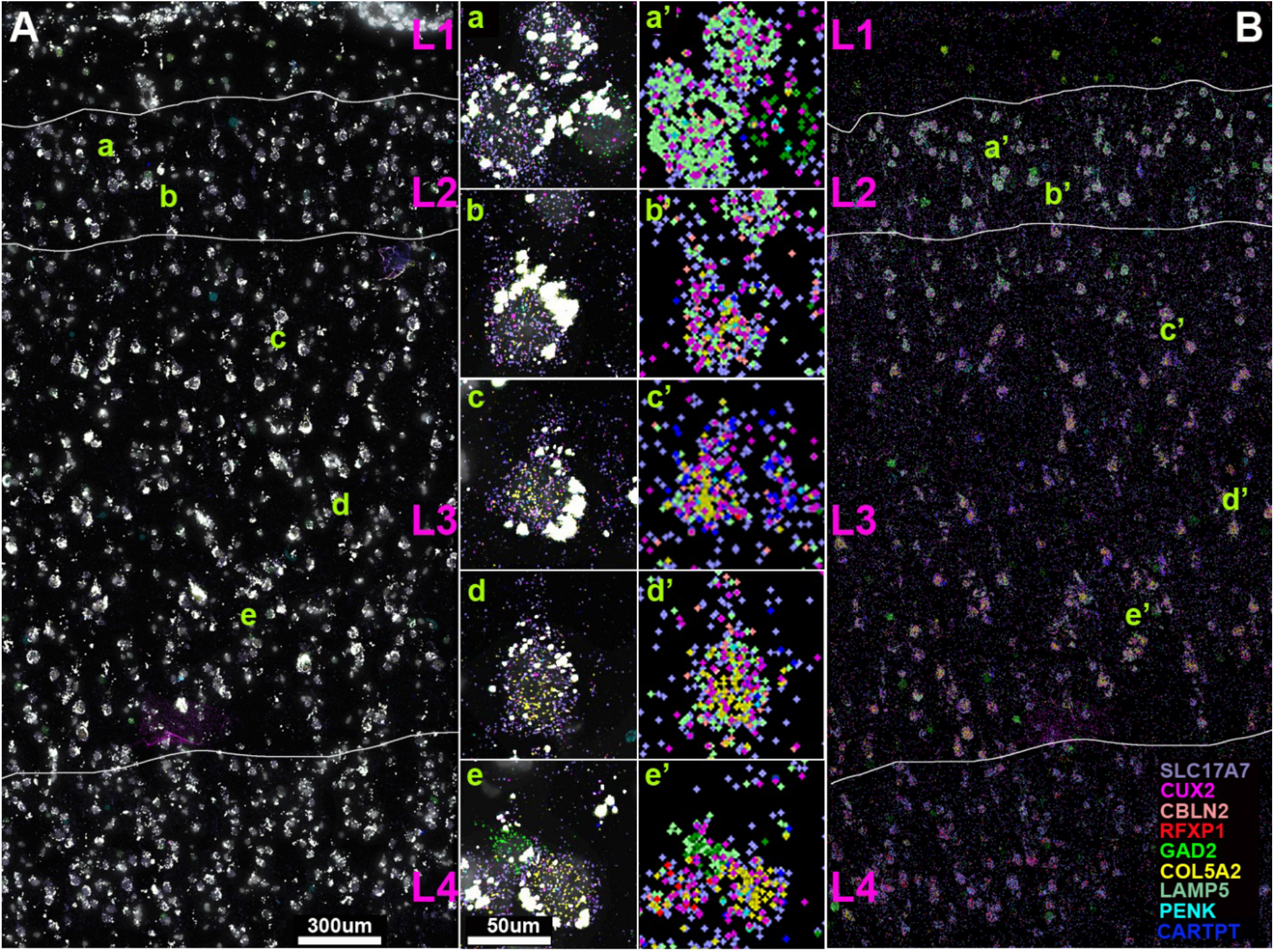
Single molecule (sm)FISH analysis of gene expression levels in human MTG layers 2 and 3. smFISH was performed with probes against *SLC17A7, CUX2, CBLN2, RFXP1, GAD2, COL5A2, LAMP5, PENK*, and *CARTPT* mRNA. **(A)** smFISH image (100x). Spots for each gene are pseudocolored as indicated in the bottom right legend. Layer demarcations are indicated in magenta. Scale bar = 300 um. B) Spot indications for each gene, pseudocolored as indicated in the bottom right legend, as in A. a,a’) Superficial layer 2 cells express *SLC17A7*(lavender), *CUX2* (magenta), and *LAMP5* (mint). b,b’) At deeper locations in layer 2, an example of an SLC17A7-expressing cell with *CUX2, LAMP5* and *COL5A2* expression. Note that *LAMP5* expression (mint) decreases in *CUX2/SLC17A7-*-expressing cells, while *COL5A2/CUX2*-expressing cells increase with depth along Layers 2 and 3 (see, c,c’; d,d’; e,e’).

**Extended Data Figure 8.**
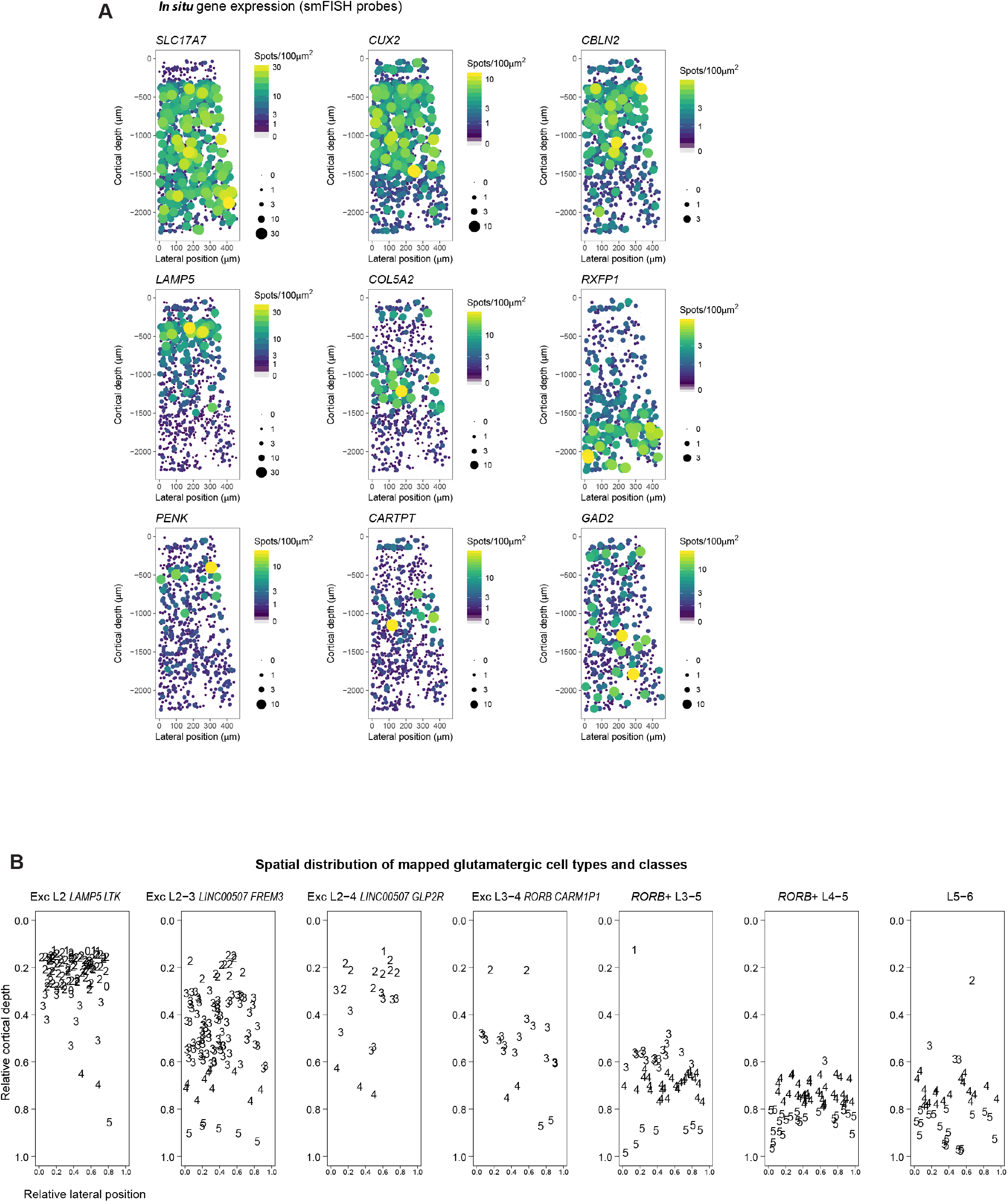
Laminar distribution of superficial excitatory neuron types validated by smFISH. **(A)** Probe density (spots per 100μm2) for 9 genes assayed across layers 1-4 (and partially layer 5) of human MTG. The cortical slice was approximately 0.5mm wide and 2mm deep. Points correspond to cellular locations *in situ* where the y-axis is the cortical depth from the pial surface and the x-axis is the lateral position. Point size and color correspond to probe density. Cells that lack probe expression are shown as small grey points. **(B)** *In situ* location of cells mapped to indicated cell types and classes (different panels) based on expression levels of 9 genes shown in (A). Numbers indicate qualitative calls of the layer to which each cell belongs based on cytoarchitecture. 0 indicates that the cell was not annotated.

**Extended Data Figure 9.**
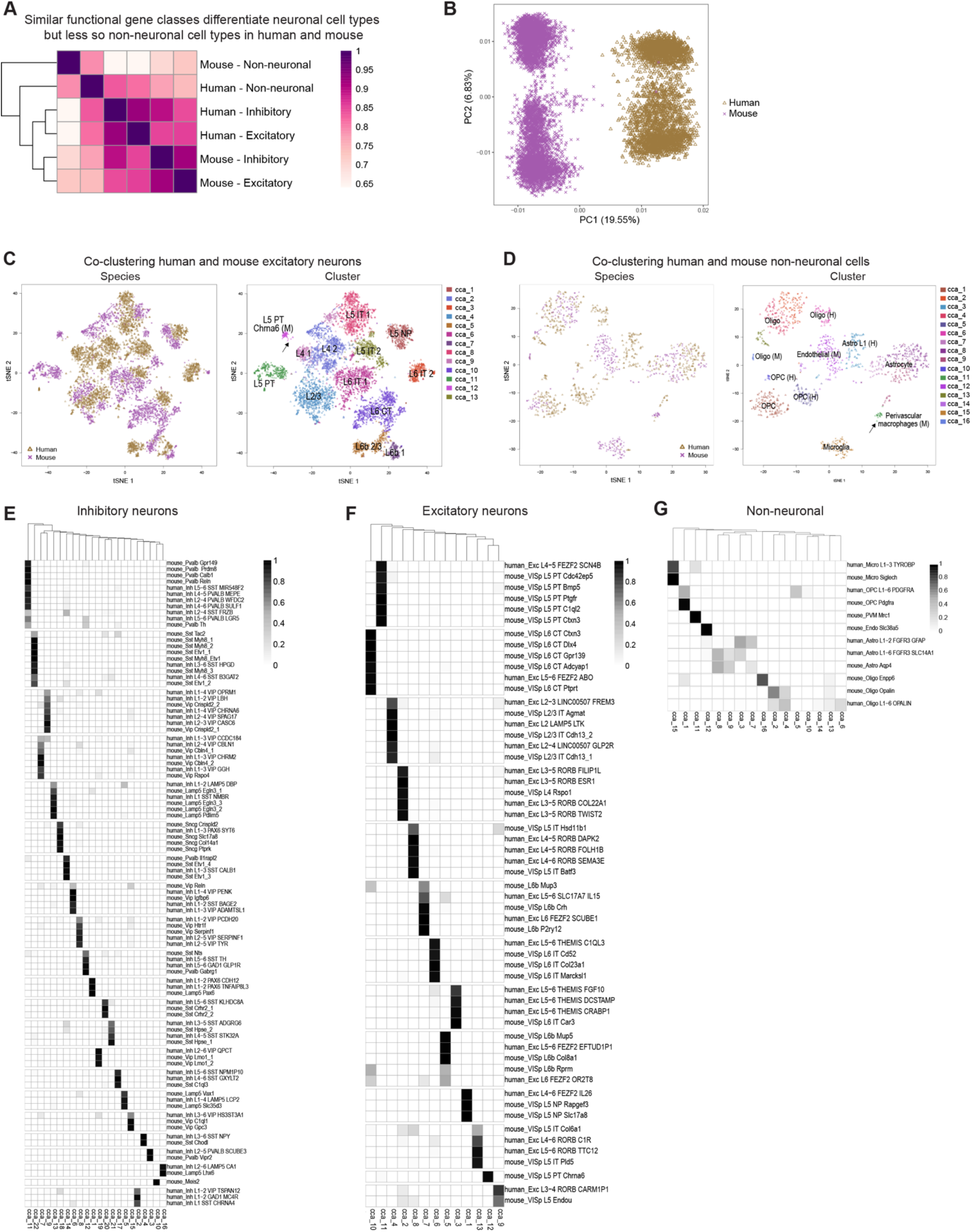
Quantifying human and mouse cell type homology. **(A)** Heatmap of Pearson’s correlations between average MetaNeighbor AUROC scores for three broad classes of human and mouse cortical cell types. Rows and columns are ordered by average-linkage hierarchical clustering. **(B)** Human (gold) and mouse (purple) GABAergic neurons projected on the first two principal components of a PCA combining expression data from both species. Almost 20% of expression differences are explained by species, while 6% are explained by major subclasses of interneurons. **(C)** t-SNE plots of first 30 basis vectors from a CCA of human and mouse glutamatergic neurons colored by species and CCA Cluster labeled with (M) contains only mouse cells. cluster. Arrow highlights two human nuclei that cluster with the mouse-specific (M) L5 PT Chrna6 cluster. **(D)** t-SNE plots of first 10 basis vectors from a CCA of human and mouse non-neuronal cells colored by species and CCA Clusters labeled with (M) or (H) contain only mouse cells or human nuclei, respectively. cluster. Human-specific (H) and mouse-specific (M) clusters are labeled. Arrow highlights two human nuclei that cluster with mouse perivascular macrophages. **(E-G)** Heatmaps showing the proportion of each human and mouse cluster (rows) that are members of each CCA cluster (columns) for GABAergic neurons (E), glutamatergic neurons (F), and non-neuronal cells (G). Rows and columns are hierarchically clustered, and most CCA clusters include human and mouse clusters that allows inference of homology between these clusters.

**Extended Data Figure 10.**
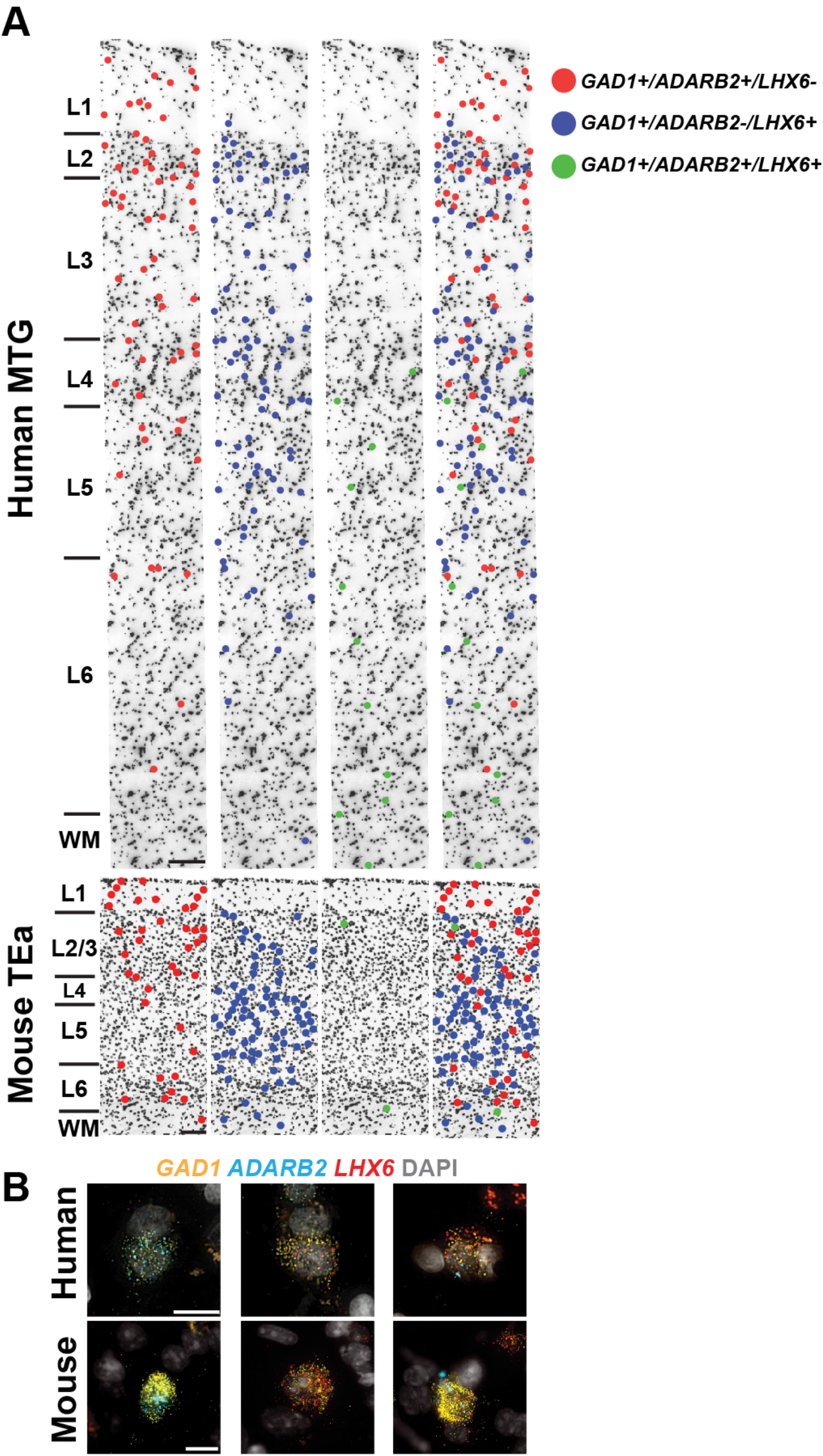
RNAscope multiplex FISH staining for broad interneuron class markers in human MTG and mouse temporal association area (TEa) **(A)** A representative inverted DAPI-stained cortical column illustrating the laminar positions of cells combinatorially labeled with broad interneuron class markers. Human MTG is shown in the top panels and mouse TEa is shown in the bottom panels. Left to right: red dots mark cells that are *GAD1+/Gad1+, ADARB2+/Adarb2+*, and *LHX6-/Lhx6-* (i.e. *ADARB2* branch interneurons); blue dots mark cells that are *GAD1+/Gad1+, ADARB2-/Adarb2-*, and *LHX6+/Lhx6+* (i.e. *LHX6* branch interneurons); green dots mark cells that are *GAD1+/Gad1+, ADARB2+/Adarb2+, LHX6+/Lhx6+* (i.e. Inh L2-6 *LAMP5 CA1* cells [human] or Lamp5 Lhx6 cells [mouse]); the far-right panel shows all the labeled cell classes overlaid onto the cortical column. **(B)** Representative images of cells labeled with the *GAD1, ADARB2*, and *LHX6* gene panel for human (top) and mouse (bottom). Left to right: cells double positive for *GAD1* and *ADARB2;* cells double positive for *GAD1* and *LHX6; GAD1, ADARB2*, and *LHX6* triple positive cells. Scale bars, 15μm (human), 10μm (mouse).

**Extended Data Figure 11.**
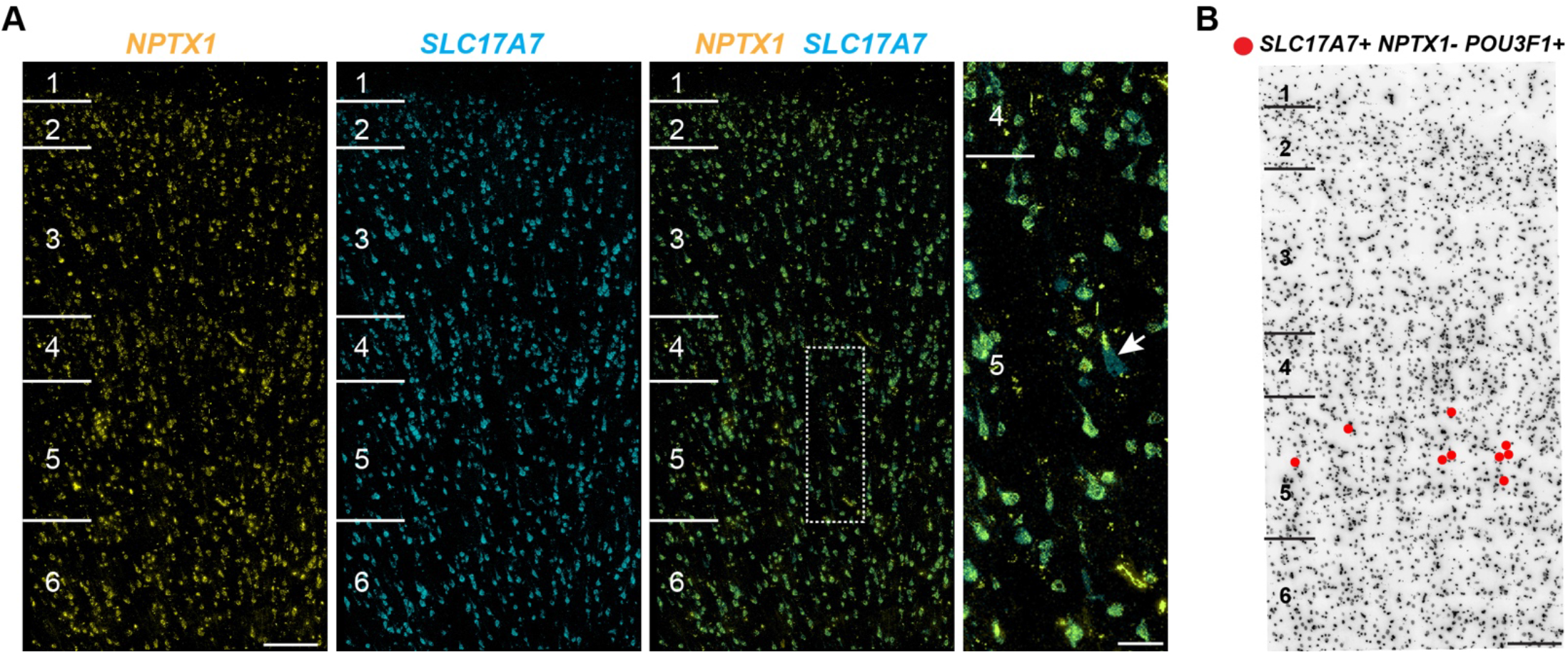
RNAscope multiplex FISH for markers of putative pyramidal tract (PT) neurons in human MTG. **(A)** FISH for *NPTX1*, a marker of non-PT excitatory types and *SLC17A7*, which is expressed in all excitatory neurons, shows that *NPTX1* labels most *SLC17A7+* cells across all cortical layers. The area indicated by the boxed region in the overlaid image of *NPTX1* and *SCL17A7* staining is shown at higher magnification in the adjacent panel to the right. One *SLC17A7+* cell, indicated with the white arrow, is *NPTX1-*, but the rest of the *SLC17A7+* cells in the field of view are *NPTX1+*. Scale bars, left (200um), right (50um). **(B)** A Representative inverted DAPI-stained cortical column overlaid with red dots that represent *SLC17A7+, NPTX1-*, and *POU3F1+* cells. *POU3F1* is a specific marker of the putative PT type (Exc L4-5 *FEZF2 SCN4B*). Scale bar, 200μm.

## Supplementary Data

### Semantic representation of cell type clusters

To provide for an unambiguous, rigorous, and informative semantic representation of the cell types defined using the single nucleus RNA sequencing gene expression clusters, we developed a strategy for defining provisional cell types (pCL) in an rdf representation. We previously proposed that these provisional cell types could be defined using a combination of the tissue anatomic structure from which the assayed specimen was derived (e.g. middle temporal gyrus - go_sc:part_of uberon:UBER0N_0002771), a set of marker genes whose combination is uniquely expressed in cells of the type (e.g. mtg_cluster_sc:selectively_expresses hugo:HGNC_8620 and mtg_cluster_sc:selectively_expresses hugo:HGNC_1751), and a supertype parent class represented in either the pCL or in the reference Cell Ontology (CL) (e.g. skos:broader :pCL78) [Bakken 2017, Aevermann 2018]. In addition, here we have also made the distinction between the experimental evidence supporting the existence of the provisional cell type class in the form of a single cell expression cluster ID (mtg_cluster_sc:evidence “Inh L1-2 PAX6 CDH12”) and the provisional cell type class itself (e.g. pCL1). We also included additional knowledge about the location of instances of each provisional cell type class within specific layers of the cerebral cortex, both in terms of any layer in which the cell body of the type is found to be localized in (mtg_cluster_sc:has_soma_location_in) and in terms of the layer that a cell of the type is preferentially enriched in (mtg_cluster_sc:enriched_in). Finally, we included the cell cluster size (i.e. the number of single nuclei included in the cell type cluster from this particular experiment) as a rough estimate of the relative cell type abundance in the specimen assayed. This rdf representation is amenable to query-based inferencing using SPARQL.

Aevermann BD, Novotny M, Bakken T, Miller JA, Diehl AD, Osumi-Sutherland D, Lasken RS, Lein ES, Scheuermann RH. Cell type discovery using single-cell transcriptomics: implications for ontological representation. Hum Mol Genet. 2018 May 1;27(R1):R40-R47. doi:10.1093/hmg/ddy100. PubMed PMID: 29590361; PubMed Central PMCID: PMC5946857.

Bakken T, Cowell L, Aevermann BD, Novotny M, Hodge R, Miller JA, Lee A, Chang I, McCorrison J, Pulendran B, Qian Y, Schork Nj, Lasken RS, Lein ES, Scheuermann RH. Cell type discovery and representation in the era of high-content single cell phenotyping. BMC Bioinformatics. 2017 Dec 21;18(Suppl 17):559. doi: 10.1186/s12859-017-1977-1. PubMed PMID: 29322913; PubMed Central PMCID: PMC5763450.

~~~
# ******************* Required IRcsourccs *******************
@prefix rdf:   <http://www.w3.org/1999/02/22-rdf-syntax-ns#>.
@prefix rdfs:   <http://www.w3.org/2000/01/rdf-schema#>.
@prefix xsd:   <http://www.w3.org/2001/XMLSchema#>.
@prefix owl:   <http://www.w3.org/2002/07/owl#>.
@prefix:       <http://www.jcvi.org/clext/mtgcluster#>.
@prefix mtg_cluster_sc: <http://www.jcvi.org/clext/mtgclusterschema#>.
@prefix go_sc:   <http://www.jcvi.org/clext/goschema#>.
@prefix cl:   <http://purl.obolibrary.org/obo/cl#>.
@prefix hugo:   <http://ncicb.nci.nih.gov/xml/owl/evs/hugo#>.
@prefix uberon:   <http://purl.obolibrary.org/obo/uberon#>.
@prefix skos:     <http://www.w3.org/2004/02/skos/core#>.
<http://www.jcvi.org/clext/mtgcluster>
 rdf:type owl:Ontology ;
owl:versionlnfo “$Id: mtg_cluster_v4.ttl, Ver 1.0, May 23, 2018, Mohamed Keshk. $”.
# ******************* MTG_Cluster_Annotation Concepts *******************
~~~

~~~
:pCL1
 a skos:Concept ;
 mtg_cluster_sc:id “pCL1” ;
 skos:broader :pCL78 ;
rdfs:label “PAX6|CDH12-expressing cerebral cortex MTG GABAergic interneuron” ;
 mtg_cluster_sc:evidence “Inh L1-2 PAX6 CDH12” ;
 go_sc:part_of uberon:UBERON_0002771 ;
 mtg_cluster_sc:enriched_in “cortical_layer1” ;
 mtg_cluster_sc:has_soma_location_in “cortical_layer1” ;
 mtg_cluster_sc:has_soma_location_in “cortical_layer2” ;
 mtg_cluster_sc:selectively_expresses hugo:HGNC_8620 ;
 mtg_cluster_sc:selectively_expresses hugo:HGNC_1751 ;
 mtg_cluster_sc:neuron_type “GABAergic” ;
 mtg_cluster_sc:cluster_size “90”^^xsd:int ;
.
~~~

~~~
:pCL2
 a skos:Concept ;
 mtg_cluster_sc:id “pCL2” ;
 skos:broader :pCL78 ;
 rdfs:label “PAX6|TNFAIP8L3-expressing cerebral cortex MTG GABAergic interneuron” ;
 mtg_cluster_sc:evidence “Inh L1-2 PAX6 TNFAIP8L3” ;
 go_sc:part_of uberon:UBER0N_0002771 ;
 mtg_cluster_sc:enriched_in “cortical_layer2” ;
 mtg_cluster_sc:has_soma_location_in “cortical_layer1” ;
 mtg_cluster_sc:has_soma_location_in “cortical_layer2” ;
 mtg_cluster_sc:selectively_expresses hugo:HGNC_8620 ;
 mtg_cluster_sc:selectively_expresses hugo:HGNC_20620 ;
 mtg_cluster_sc:neuron_type “GABAergic” ;
 mtg_cluster_sc:cluster_size “16”^^xsd:int ;
.
~~~

~~~
:pCL3
 a skos:Concept ;
 mtg_cluster_sc:id “pCL3” ;
 skos:broader :pCL78 ;
rdfs:label “SST|NMBR-expressing cerebral cortex MTG GABAergic interneuron” ;
 mtg_cluster_sc:evidence “Inh L1 SST NMBR” ;
 go_sc:part_of uberon:UBER0N_0002771 ;
 mtg_cluster_sc:enriched_in “cortical_layer1” ;
 mtg_cluster_sc:has_soma_location_in “cortical_layer1” ;
 mtg_cluster_sc:selectively_expresses hugo:HGNC_11329 ;
 mtg_cluster_sc:selectively_expresses hugo:HGNC_7843 ;
 mtg_cluster_sc:neuron_type “GABAergic” ;
 mtg_cluster_sc:cluster_size “283”^^xsd:int ;
.
~~~

~~~
:pCL4
 a skos:Concept ;
 mtg_cluster_sc:id “pCL4” ;
 skos:broader :pCL78 ;
rdfs:label “LAMP5|LCP2-expressing cerebral cortex MTG GABAergic interneuron” ;
 mtg_cluster_sc:evidence “Inh L1-4 LAMP5 LCP2” ;
 go_sc:part_of uberon:UBERON_0002771 ;
 mtg_cluster_sc:enriched_in “cortical_layer3” ;
 mtg_cluster_sc:has_soma_location_in “cortical_layer1” ;
 mtg_cluster_sc:has_soma_location_in “cortical_layer2” ;
 mtg_cluster_sc:has_soma_location_in “cortical_layer3” ;
 mtg_cluster_sc:has_soma_location_in “cortical_layer4” ;
 mtg_cluster_sc:selectively_expresses hugo:HGNC_16097 ;
 mtg_cluster_sc:selectively_expresses hugo:HGNC_6529 ;
 mtg_cluster_sc:neuron_type “GABAergic” ;
 mtg_cluster_sc:cluster_size “356”^^xsd:int ;
.
~~~

~~~
:pCL5
.
 a skos:Concept ;
 mtg_cluster_sc:id “pCL5” ;
 skos:broader :pCL78 ;
 rdfs:label “LAMP5|DBP-expressing cerebral cortex MTG GABAergic interneuron” ;
 mtg_cluster_sc:evidence “Inh L1-2 LAMP5 DBP” ;
 go_sc:part_of uberon:UBERON_0002771 ;
 mtg_cluster_sc:enriched_in “cortical_layer1” ;
 mtg_cluster_sc:has_soma_location_in “cortical_layer1” ;
 mtg_cluster_sc:has_soma_location_in “cortical_layer2” ;
 mtg_cluster_sc:selectively_expresses hugo:HGNC_16097 ;
mtg_cluster_sc:selectively_expresses hugo:HGNC_2697 ;
 mtg_cluster_sc:neuron_type “GABAergic” ;
 mtg_cluster_sc:cluster_size “21”^^xsd:int ;
.
~~~

~~~
:pCL6
 a skos:Concept ;
 mtg_cluster_sc:id “pCL6” ;
 skos:broader :pCL78 ;
 rdfs:label “LAMP5|CA1-expressing cerebral cortex MTG GABAergic interneuron” ;
 mtg_cluster_sc:evidence “Inh L2-6 LAMP5 CA1” ;
 go_sc:part_of uberon:UBERON_0002771 ;
 mtg_cluster_sc:enriched_in “cortical_layer5” ;
 mtg_cluster_sc:has_soma_location_in “cortical_layer2” ;
 mtg_cluster_sc:has_soma_location_in “cortical_layer3” ;
 mtg_cluster_sc:has_soma_location_in “cortical_layer4” ;
 mtg_cluster_sc:has_soma_location_in “cortical_layer5” ;
 mtg_cluster_sc:has_soma_location_in “cortical_layer6” ;
 mtg_cluster_sc:selectively_expresses hugo:HGNC_16097 ;
 mtg_cluster_sc:selectively_expresses hugo:HGNC_1368 ;
 mtg_cluster_sc:neuron_type “GABAergic” ;
 mtg_cluster_sc:cluster_size “256”^^xsd:int ;
.
~~~

~~~
:pCL7
 a skos:Concept ;
 mtg_cluster_sc:id “pCL7” ;
 skos:broader :pCL78 ;
rdfs:label “SST|CHRNA4-expressing cerebral cortex MTG GABAergic interneuron” ;
 mtg_cluster_sc:evidence “Inh L1 SST CHRNA4” ;
 go_sc:part_of uberon:UBERON_0002771 ;
 mtg_cluster_sc:enriched_in “cortical_layer1” ;
 mtg_cluster_sc:has_soma_location_in “cortical_layer1” ;
 mtg_cluster_sc:selectively_expresses hugo:HGNC_11329 ;
 mtg_cluster_sc:selectively_expresses hugo:HGNC_1958 ;
 mtg_cluster_sc:neuron_type “GABAergic” ;
 mtg_cluster_sc:cluster_size “52”^^xsd:int ;
.
~~~

~~~
:pCL8
 a skos:Concept ;
 mtg_cluster_sc:id “pCL8” ;
 skos:broader :pCL78 ;
 rdfs:label “GAD1|MC4R-expressing cerebral cortex MTG GABAergic interneuron” ;
 mtg_cluster_sc:evidence “Inh L1-2 GAD1 MC4R” ;
 go_sc:part_of uberon:UBER0N_0002771 ;
 mtg_cluster_sc:enriched_in “cortical_layer1” ;
 mtg_cluster_sc:has_soma_location_in “cortical_layer1” ;
 mtg_cluster_sc:has_soma_location_in “cortical_layer2” ;
 mtg_cluster_sc:selectively_expresses hugo:HGNC_4092 ;
 mtg_cluster_sc:selectively_expresses hugo:HGNC_6932 ;
 mtg_cluster_sc:neuron_type “GABAergic” ;
 mtg_cluster_sc:cluster_size “107”^^xsd:int ;
.
~~~

~~~
:pCL9
 a skos:Concept ;
 mtg_cluster_sc:id “pCL9” ;
 skos:broader :pCL78 ;
 rdfs:label “SST|BAGE2-expressing cerebral cortex MTG GABAergic interneuron” ;
 mtg_cluster_sc:evidence “Inh L1-2 SST BAGE2” ;
 go_sc:part_of uberon:UBER0N_0002771 ;
 mtg_cluster_sc:enriched_in “cortical_layer1” ;
 mtg_cluster_sc:has_soma_location_in “cortical_layer1” ;
 mtg_cluster_sc:has_soma_location_in “cortical_layer2” ;
 mtg_cluster_sc:selectively_expresses hugo:HGNC_11329 ;
 mtg_cluster_sc:selectively_expresses hugo:HGNC_15723 ;
 mtg_cluster_sc:neuron_type “GABAergic” ;
 mtg_cluster_sc:cluster_size “108”^^xsd:int ;
.
~~~

~~~
:pCL10
 a skos:Concept ;
 mtg_cluster_sc:id “pCL10” ;
 skos:broader :pCL80 ;
 rdfs:label “PAX6|SYT6-expressing cerebral cortex MTG GABAergic interneuron” ;
 mtg_cluster_sc:evidence “Inh L1 -3 PAX6 SYT6” ; go_sc:part_of uberon:UBERON_0002771 ;
 mtg_cluster_sc:enriched_in “cortical_layer2” ;
 mtg_cluster_sc:has_soma_location_in “cortical_layer1” ;
 mtg_cluster_sc:has_soma_location_in “cortical_layer2” ;
 mtg_cluster_sc:has_soma_location_in “cortical_layer3” ;
 mtg_cluster_sc:selectively_expresses hugo:HGNC_8620 ;
 mtg_cluster_sc:selectively_expresses hugo:HGNC_18638 ;
 mtg_cluster_sc:neuron_type “GABAergic” ;
 mtg_cluster_sc:cluster_size “29”^^xsd:int ;
.
~~~

~~~
:pCL11
 a skos:Concept ;
 mtg_cluster_sc:id “pCL11” ;
 skos:broader :pCL80 ;
 rdfs:label “TSPAN12-expressing cerebral cortex MTG GABAergic interneuron” ;
 mtg_cluster_sc:evidence “Inh L1-2 VIP TSPAN12” ; go_sc:part_of uberon:UBER0N_0002771 ;
 mtg_cluster_sc:enriched_in “cortical_layer1” ;
 mtg_cluster_sc:has_soma_location_in “cortical_layer1” ;
 mtg_cluster_sc:has_soma_location_in “cortical_layer2” ;
 mtg_cluster_sc:selectively_expresses hugo:HGNC_21641 ;
 mtg_cluster_sc:neuron_type “GABAergic” ;
 mtg_cluster_sc:cluster_size “42”^^xsd:int ;
.
~~~

~~~
:pCL12
 a skos:Concept ;
 mtg_cluster_sc:id “pCL12” ;
 skos:broader :pCL80 ;
 rdfs:label “CHRNA6-expressing cerebral cortex MTG GABAergic interneuron” ;
 mtg_cluster_sc:evidence “Inh L1-4 VIP CHRNA6” ;
 go_sc:part_of uberon:UBER0N_0002771 ;
 mtg_cluster_sc:enriched_in “cortical_layer3” ;
 mtg_cluster_sc:has_soma_location_in “cortical_layer1” ;
 mtg_cluster_sc:has_soma_location_in “cortical_layer2” ;
 mtg_cluster_sc:has_soma_location_in “cortical_layer3” ;
 mtg_cluster_sc:has_soma_location_in “cortical_layer4” ;
 mtg_cluster_sc:selectively_expresses hugo:HGNC_15963 ;
 mtg_cluster_sc:neuron_type “GABAergic” ;
 mtg_cluster_sc:cluster_size “25”^^xsd:int ;
.
~~~

~~~
:pCL13
 a skos:Concept ;
 mtg_cluster_sc:id “pCL13” ;
 skos:broader :pCL80 ;
 rdfs:label “ADAMTSL1-expressing cerebral cortex MTG GABAergic interneuron” ;
 mtg_cluster_sc:evidence “Inh L1 -3 VIP ADAMTSL1” ;
 go_sc:part_of uberon:UBERON_0002771 ;
 mtg_cluster_sc:enriched_in “cortical_layer3” ;
 mtg_cluster_sc:has_soma_location_in “cortical_layer1” ;
 mtg_cluster_sc:has_soma_location_in “cortical_layer2” ;
 mtg_cluster_sc:has_soma_location_in “cortical_layer3” ;
 mtg_cluster_sc:selectively_expresses hugo:HGNC_14632 ;
 mtg_cluster_sc:neuron_type “GABAergic” ;
 mtg_cluster_sc:cluster_size “72”^^xsd:int ;
.
~~~

~~~
:pCL14
 a skos:Concept ;
 mtg_cluster_sc:id “pCL14” ;
 skos:broader :pCL80 ;
 rdfs:label “PENK-expressing cerebral cortex MTG GABAergic interneuron” ;
 mtg_cluster_sc:evidence “Inh L1-4 VIP PENK” ;
 go_sc:part_of uberon:UBERON_0002771 ;
 mtg_cluster_sc:enriched_in “cortical_layer3” ;
 mtg_cluster_sc:has_soma_location_in “cortical_layer1” ;
 mtg_cluster_sc:has_soma_location_in “cortical_layer2” ;
 mtg_cluster_sc:has_soma_location_in “cortical_layer3” ;
 mtg_cluster_sc:has_soma_location_in “cortical_layer4” ;
 mtg_cluster_sc:selectively_expresses hugo:HGNC_8831 ;
 mtg_cluster_sc:neuron_type “GABAergic” ;
 mtg_cluster_sc:cluster_size “17”^^xsd:int ;
.
~~~

~~~
:pCL15
 a skos:Concept ;
 rdfs:label “QPCT-expressing cerebral cortex MTG GABAergic interneuron” ;
 mtg_cluster_sc:evidence “Inh L2-6 VIP QPCT” ;
 go_sc:part_of uberon:UBER0N_0002771 ;
 mtg_cluster_sc:enriched_in “cortical_layer4” ;
 mtg_cluster_sc:has_soma_location_in “cortical_layer2” ;
 mtg_cluster_sc:has_soma_location_in “cortical_layer3” ;
 mtg_cluster_sc:has_soma_location_in “cortical_layer4” ;
 mtg_cluster_sc:has_soma_location_in “cortical_layer5” ;
 mtg_cluster_sc:has_soma_location_in “cortical_layer6” ;
 mtg_cluster_sc:selectively_expresses hugo:HGNC_9753 ;
 mtg_cluster_sc:neuron_type “GABAergic” ;
 mtg_cluster_sc:cluster_size “37”^^xsd:int ;
.
~~~

~~~
:pCL16
 a skos:Concept ;
 mtg_cluster_sc:id “pCL16” ;
 skos:broader :pCL80 ;
rdfs:label “HS3ST3A1-expressing cerebral cortex MTG GABAergic interneuron” ;
 mtg_cluster_sc:evidence “Inh L3-6 VIP HS3ST3A1” ;
 go_sc:part_of uberon:UBER0N_0002771 ;
 mtg_cluster_sc:enriched_in “cortical_layer4” ;
 mtg_cluster_sc:has_soma_location_in “cortical_layer3” ;
 mtg_cluster_sc:has_soma_location_in “cortical_layer4” ;
 mtg_cluster_sc:has_soma_location_in “cortical_layer5” ;
 mtg_cluster_sc:has_soma_location_in “cortical_layer6” ;
 mtg_cluster_sc:selectively_expresses hugo:HGNC_5196 ;
 mtg_cluster_sc:neuron_type “GABAergic” ;
~~~

~~~
:pCL17 a skos:Concept ;
 mtg_cluster_sc:id “pCL17” ;
 skos:broader :pCL80 ;
 rdfs:label “PCDH20-expressing cerebral cortex MTG GABAergic interneuron” ;
 mtg_cluster_sc:evidence “Inh L1-2 VIP PCDH20” ;
 go_sc:part_of uberon:UBERON_0002771 ;
 mtg_cluster_sc:enriched_in “cortical_layer2” ;
 mtg_cluster_sc:has_soma_location_in “cortical_layer1” ;
 mtg_cluster_sc:has_soma_location_in “cortical_layer2” ;
 mtg_cluster_sc:selectively_expresses hugo:HGNC_14257 ;
 mtg_cluster_sc:neuron_type “GABAergic” ;
 mtg_cluster_sc:cluster_size “61”^^xsd:int ;
.
~~~

~~~
:pCL18
 a skos:Concept ;
 mtg_cluster_sc:id “pCL18” ;
 skos:broader :pCL80 ;
 rdfs:label “SERPINF1-expressing cerebral cortex MTG GABAergic interneuron” ;
 mtg_cluster_sc:evidence “Inh L2-5 VIP SERPINF1” ;
 go_sc:part_of uberon:UBERON_0002771 ;
 mtg_cluster_sc:enriched_in “cortical_layer4” ;
 mtg_cluster_sc:has_soma_location_in “cortical_layer2” ;
 mtg_cluster_sc:has_soma_location_in “cortical_layer3” ;
 mtg_cluster_sc:has_soma_location_in “cortical_layer4” ;
 mtg_cluster_sc:has_soma_location_in “cortical_layer5” ;
 mtg_cluster_sc:selectively_expresses hugo:HGNC_8824 ;
mtg_cluster_sc:neuron_type “GABAergic” ;
 mtg_cluster_sc:cluster_size “55”^^xsd:int ;
.
~~~

~~~
:pCL19
 a skos:Concept ;
 mtg_cluster_sc:id “pCL19” ;
 skos:broader :pCL80 ;
 rdfs:label “TYR-expressing cerebral cortex MTG GABAergic interneuron” ;
 mtg_cluster_sc:evidence “Inh L2-5 VIP TYR” ;
 go_sc:part_of uberon:UBER0N_0002771 ;
 mtg_cluster_sc:enriched_in “cortical_layer4” ;
 mtg_cluster_sc:has_soma_location_in “cortical_layer2” ;
 mtg_cluster_sc:has_soma_location_in “cortical_layer3” ;
 mtg_cluster_sc:has_soma_location_in “cortical_layer4” ;
 mtg_cluster_sc:has_soma_location_in “cortical_layer5” ;
 mtg_cluster_sc:selectively_expresses hugo:HGNC_12442 ;
 mtg_cluster_sc:neuron_type “GABAergic” ;
 mtg_cluster_sc:cluster_size “62”^^xsd:int ;
.
~~~

~~~
:pCL20
 a skos:Concept ;
 mtg_cluster_sc:id “pCL20” ;
 skos:broader :pCL80 ;
 rdfs:label “CHRM2-expressing cerebral cortex MTG GABAergic interneuron” ;
 mtg_cluster_sc:evidence “Inh L1 -3 VIP CHRM2” ;
 go_sc:part_of uberon:UBER0N_0002771 ;
 mtg_cluster_sc:enriched_in “cortical_layer2” ;
 mtg_cluster_sc:has_soma_location_in “cortical_layer1” ;
 mtg_cluster_sc:has_soma_location_in “cortical_layer2” ;
 mtg_cluster_sc:has_soma_location_in “cortical_layer3” ;
 mtg_cluster_sc:selectively_expresses hugo:HGNC_1951 ;
 mtg_cluster_sc:neuron_type “GABAergic” ;
 mtg_cluster_sc:cluster_size “175”^^xsd:int ;
.
~~~

~~~
:pCL21
 a skos:Concept ;
 mtg_cluster_sc:id “pCL21” ; skos:broader :pCL80 ;
 rdfs:label “CBLN1-expressing cerebral cortex MTG GABAergic interneuron” ;
 mtg_cluster_sc:evidence “Inh L2-4 VIP CBLN1” ;
 go_sc:part_of uberon:UBERON_0002771 ;
 mtg_cluster_sc:enriched_in “cortical_layer3” ;
 mtg_cluster_sc:has_soma_location_in “cortical_layer2” ;
 mtg_cluster_sc:has_soma_location_in “cortical_layer3” ;
 mtg_cluster_sc:has_soma_location_in “cortical_layer4” ;
 mtg_cluster_sc:selectively_expresses hugo:HGNC_1543 ;
 mtg_cluster_sc:neuron_type “GABAergic” ;
 mtg_cluster_sc:cluster_size “67”^^xsd:int ;
.
~~~

~~~
:pCL22
 a skos:Concept ;
 mtg_cluster_sc:id “pCL22” ; skos:broader :pCL80 ;
rdfs:label “CCDC184-expressing cerebral cortex MTG GABAergic interneuron” ;
 mtg_cluster_sc:evidence “Inh L1 -3 VIP CCDC184” ;
 go_sc:part_of uberon:UBERON_0002771 ;
 mtg_cluster_sc:enriched_in “cortical_layer2” ;
 mtg_cluster_sc:has_soma_location_in “cortical_layer1” ;
mtg_cluster_sc:has_soma_location_in “cortical_layer2” ;
 mtg_cluster_sc:has_soma_location_in “cortical_layer3” ;
 mtg_cluster_sc:selectively_expresses hugo:HGNC_33749 ;
 mtg_cluster_sc:neuron_type “GABAergic” ;
 mtg_cluster_sc:cluster_size “64”^^xsd:int ;
.
~~~

~~~
:pCL23
 a skos:Concept ;
 mtg_cluster_sc:id “pCL23” ;
 skos:broader :pCL80 ;
 rdfs:label “GGH-expressing cerebral cortex MTG GABAergic interneuron” ;
 mtg_cluster_sc:evidence “Inh L1 -3 VIP GGH” ;
 go_sc:part_of uberon:UBER0N_0002771 ;
 mtg_cluster_sc:enriched_in “cortical_layer2” ;
 mtg_cluster_sc:has_soma_location_in “cortical_layer1” ;
 mtg_cluster_sc:has_soma_location_in “cortical_layer2” ;
 mtg_cluster_sc:has_soma_location_in “cortical_layer3” ;
 mtg_cluster_sc:selectively_expresses hugo:HGNC_4248 ;
 mtg_cluster_sc:neuron_type “GABAergic” ;
 mtg_cluster_sc:cluster_size “68”^^xsd:int ;
.
~~~

~~~
:pCL24
 a skos:Concept ;
 mtg_cluster_sc:id “pCL24” ; skos:broader :pCL80 ;
 rdfs:label “LBH-expressing cerebral cortex MTG GABAergic interneuron” ;
 mtg_cluster_sc:evidence “Inh L1-2 VIP LBH” ;
 go_sc:part_of uberon:UBER0N_0002771 ;
 mtg_cluster_sc:enriched_in “cortical_layer2” ;
 mtg_cluster_sc:has_soma_location_in “cortical_layer1” ;
 mtg_cluster_sc:has_soma_location_in “cortical_layer2” ;
 mtg_cluster_sc:selectively_expresses hugo:HGNC_29532 ;
 mtg_cluster_sc:neuron_type “GABAergic” ;
 mtg_cluster_sc:cluster_size “47”^^xsd:int ;
.
~~~

~~~
:pCL25
 a skos:Concept ;
 mtg_cluster_sc:id “pCL25” ;
 skos:broader :pCL80 ;
 rdfs:label “CASC6-expressing cerebral cortex MTG GABAergic interneuron” ;
 mtg_cluster_sc:evidence “Inh L2-3 VIP CASC6” ;
 go_sc:part_of uberon:UBERON_0002771 ;
 mtg_cluster_sc:enriched_in “cortical_layer2” ;
 mtg_cluster_sc:has_soma_location_in “cortical_layer2” ;
 mtg_cluster_sc:has_soma_location_in “cortical_layer3” ;
 mtg_cluster_sc:selectively_expresses hugo:HGNC_49076 ;
 mtg_cluster_sc:neuron_type “GABAergic” ;
 mtg_cluster_sc:cluster_size “45”^^xsd:int ;
.
~~~

~~~
:pCL26
 a skos:Concept ;
 mtg_cluster_sc:id “pCL26” ;
 skos:broader :pCL80 ;
 rdfs:label “SPAG17-expressing cerebral cortex MTG GABAergic interneuron” ;
 mtg_cluster_sc:evidence “Inh L2-4 VIP SPAG17” ;
 go_sc:part_of uberon:UBERON_0002771 ;
 mtg_cluster_sc:enriched_in “cortical_layer3” ;
 mtg_cluster_sc:has_soma_location_in “cortical_layer2” ;
 mtg_cluster_sc:has_soma_location_in “cortical_layer3” ;
 mtg_cluster_sc:has_soma_location_in “cortical_layer4” ;
 mtg_cluster_sc:selectively_expresses hugo:HGNC_26620 ;
 mtg_cluster_sc:neuron_type “GABAergic” ;
 mtg_cluster_sc:cluster_size “33”^^xsd:int ;
.
~~~

~~~
:pCL27
 a skos:Concept ;
 mtg_cluster_sc:id “pCL27” ;
 skos:broader :pCL80 ;
 rdfs:label “0PRM1-expressing cerebral cortex MTG GABAergic interneuron” ;
 mtg_cluster_sc:evidence “Inh L1-4 VIP 0PRM1” ;
 go_sc:part_of uberon:UBER0N_0002771 ;
 mtg_cluster_sc:enriched_in “cortical_layer3” ;
 mtg_cluster_sc:has_soma_location_in “cortical_layer1” ;
 mtg_cluster_sc:has_soma_location_in “cortical_layer2” ;
 mtg_cluster_sc:has_soma_location_in “cortical_layer3” ;
 mtg_cluster_sc:has_soma_location_in “cortical_layer4” ;
 mtg_cluster_sc:selectively_expresses hugo:HGNC_8156 ;
 mtg_cluster_sc:neuron_type “GABAergic” ;
 mtg_cluster_sc:cluster_size “52”^^xsd:int ;
.
~~~

~~~
:pCL28
 a skos:Concept ;
 mtg_cluster_sc:id “pCL28” ; skos:broader :pCL81 ;
 rdfs:label “NPY-expressing cerebral cortex MTG GABAergic interneuron” ;
 mtg_cluster_sc:evidence “Inh L3-6 SST NPY” ;
 go_sc:part_of uberon:UBER0N_0002771 ;
 mtg_cluster_sc:enriched_in “cortical_layer5” ;
 mtg_cluster_sc:has_soma_location_in “cortical_layer3” ;
 mtg_cluster_sc:has_soma_location_in “cortical_layer4” ;
 mtg_cluster_sc:has_soma_location_in “cortical_layer5” ;
 mtg_cluster_sc:has_soma_location_in “cortical_layer6” ;
 mtg_cluster_sc:selectively_expresses hugo:HGNC_7955 ;
 mtg_cluster_sc:neuron_type “GABAergic” ;
 mtg_cluster_sc:cluster_size “15”^^xsd:int ;
.
~~~

~~~
:pCL29
 a skos:Concept ;
 mtg_cluster_sc:id “pCL29” ;
 skos:broader :pCL81 ;
 rdfs:label “HPGD-expressing cerebral cortex MTG GABAergic interneuron” ;
 mtg_cluster_sc:evidence “Inh L3-6 SST HPGD” ;
 go_sc:part_of uberon:UBERON_0002771 ;
 mtg_cluster_sc:enriched_in “cortical_layer5” ;
 mtg_cluster_sc:has_soma_location_in “cortical_layer3” ;
 mtg_cluster_sc:has_soma_location_in “cortical_layer4” ;
 mtg_cluster_sc:has_soma_location_in “cortical_layer5” ;
 mtg_cluster_sc:has_soma_location_in “cortical_layer6” ;
 mtg_cluster_sc:selectively_expresses hugo:HGNC_5154 ;
 mtg_cluster_sc:neuron_type “GABAergic” ;
 mtg_cluster_sc:cluster_size “60”^^xsd:int ;
.
~~~

~~~
:pCL30
 a skos:Concept ;
 mtg_cluster_sc:id “pCL30” ;
 skos:broader :pCL81 ;
 rdfs:label “B3GAT2-expressing cerebral cortex MTG GABAergic interneuron” ;
 mtg_cluster_sc:evidence “Inh L4-6 SST B3GAT2” ;
 go_sc:part_of uberon:UBER0N_0002771 ;
 mtg_cluster_sc:enriched_in “cortical_layer5” ;
 mtg_cluster_sc:has_soma_location_in “cortical_layer4” ;
 mtg_cluster_sc:has_soma_location_in “cortical_layer5” ;
 mtg_cluster_sc:has_soma_location_in “cortical_layer6” ;
 mtg_cluster_sc:selectively_expresses hugo:HGNC_922 ;
 mtg_cluster_sc:neuron_type “GABAergic” ;
 mtg_cluster_sc:cluster_size “182”^^xsd:int ;
.
~~~

~~~
:pCL31
 a skos:Concept ;
 mtg_cluster_sc:id “pCL31” ;
 skos:broader :pCL81 ;
 rdfs:label “KLHDC8A-expressing cerebral cortex MTG GABAergic interneuron” ;
 mtg_cluster_sc:evidence “Inh L5-6 SST KLHDC8A” ;
 go_sc:part_of uberon:UBER0N_0002771 ;
 mtg_cluster_sc:enriched_in “cortical_layer5” ;
 mtg_cluster_sc:has_soma_location_in “cortical_layer5” ;
 mtg_cluster_sc:has_soma_location_in “cortical_layer6” ;
 mtg_cluster_sc:selectively_expresses hugo:HGNC_25573 ;
 mtg_cluster_sc:neuron_type “GABAergic” ;
 mtg_cluster_sc:cluster_size “63”^^xsd:int ;
~~~

~~~
:pCL32
 a skos:Concept ;
 mtg_cluster_sc:id “pCL32” ; skos:broader :pCL81 ;
 rdfs:label “NPM1P10-expressing cerebral cortex MTG GABAergic interneuron” ;
 mtg_cluster_sc:evidence “Inh L5-6 SST NPM1P10” ;
 go_sc:part_of uberon:UBERON_0002771 ;
 mtg_cluster_sc:enriched_in “cortical_layer5” ;
 mtg_cluster_sc:has_soma_location_in “cortical_layer5” ;
 mtg_cluster_sc:has_soma_location_in “cortical_layer6” ;
 mtg_cluster_sc:selectively_expresses hugo:HGNC_7912 ;
 mtg_cluster_sc:neuron_type “GABAergic” ;
 mtg_cluster_sc:cluster_size “79”^^xsd:int ;
.
~~~

~~~
:pCL33
 a skos:Concept ;
 mtg_cluster_sc:id “pCL33” ;
 skos:broader :pCL81 ;
 rdfs:label “GXYLT2-expressing cerebral cortex MTG GABAergic interneuron” ;
 mtg_cluster_sc:evidence “Inh L4-6 SST GXYLT2” ;
 go_sc:part_of uberon:UBERON_0002771 ;
 mtg_cluster_sc:enriched_in “cortical_layer5” ;
 mtg_cluster_sc:has_soma_location_in “cortical_layer4” ;
 mtg_cluster_sc:has_soma_location_in “cortical_layer5” ;
 mtg_cluster_sc:has_soma_location_in “cortical_layer6” ;
 mtg_cluster_sc:selectively_expresses hugo:HGNC_33383 ;
 mtg_cluster_sc:neuron_type “GABAergic” ;
 mtg_cluster_sc:cluster_size “41”^^xsd:int ;
.
~~~

~~~
:pCL34
 a skos:Concept ;
 mtg_cluster_sc:id “pCL34” ;
 skos:broader :pCL81 ;
 rdfs:label “STK32A-expressing cerebral cortex MTG GABAergic interneuron” ;
 mtg_cluster_sc:evidence “Inh L4-5 SST STK32A” ;
 go_sc:part_of uberon:UBERON_0002771 ;
 mtg_cluster_sc:enriched_in “cortical_layer4” ;
 mtg_cluster_sc:has_soma_location_in “cortical_layer4” ;
 mtg_cluster_sc:has_soma_location_in “cortical_layer5” ;
 mtg_cluster_sc:selectively_expresses hugo:HGNC_28317 ;
 mtg_cluster_sc:neuron_type “GABAergic” ;
 mtg_cluster_sc:cluster_size “93”^^xsd:int ;
.
~~~

~~~
:pCL35
 a skos:Concept ;
 mtg_cluster_sc:id “pCL35” ;
 skos:broader :pCL81 ;
 rdfs:label “CALB1-expressing cerebral cortex MTG GABAergic interneuron” ;
 mtg_cluster_sc:evidence “Inh L1 -3 SST CALB1” ;
 go_sc:part_of uberon:UBERON_0002771 ;
 mtg_cluster_sc:enriched_in “cortical_layer2” ;
 mtg_cluster_sc:has_soma_location_in “cortical_layer1” ;
 mtg_cluster_sc:has_soma_location_in “cortical_layer2” ;
 mtg_cluster_sc:has_soma_location_in “cortical_layer3” ;
 mtg_cluster_sc:selectively_expresses hugo:HGNC_1434 ;
 mtg_cluster_sc:neuron_type “GABAergic” ;
 mtg_cluster_sc:cluster_size “279”^^xsd:int ;
.
~~~

~~~
:pCL36
 a skos:Concept ;
 mtg_cluster_sc:id “pCL36” ;
 skos:broader :pCL81 ;
 rdfs:label “ADGRG6-expressing cerebral cortex MTG GABAergic interneuron” ;
 mtg_cluster_sc:evidence “Inh L3-5 SST ADGRG6” ;
 go_sc:part_of uberon:UBER0N_0002771 ;
 mtg_cluster_sc:enriched_in “cortical_layer4” ;
 mtg_cluster_sc:has_soma_location_in “cortical_layer3” ;
 mtg_cluster_sc:has_soma_location_in “cortical_layer4” ;
 mtg_cluster_sc:has_soma_location_in “cortical_layer5” ;
 mtg_cluster_sc:selectively_expresses hugo:HGNC_13841 ;
 mtg_cluster_sc:neuron_type “GABAergic” ;
 mtg_cluster_sc:cluster_size “132”^^xsd:int ;
.
~~~

~~~
:pCL37
 a skos:Concept ;
 mtg_cluster_sc:id “pCL37” ;
 skos:broader :pCL81 ;
 rdfs:label “FRZB-expressing cerebral cortex MTG GABAergic interneuron” ;
 mtg_cluster_sc:evidence “Inh L2-4 SST FRZB” ;
 go_sc:part_of uberon:UBER0N_0002771 ;
 mtg_cluster_sc:enriched_in “cortical_layer3” ;
 mtg_cluster_sc:has_soma_location_in “cortical_layer2” ;
 mtg_cluster_sc:has_soma_location_in “cortical_layer3” ;
 mtg_cluster_sc:has_soma_location_in “cortical_layer4” ;
 mtg_cluster_sc:selectively_expresses hugo:HGNC_3959 ;
 mtg_cluster_sc:neuron_type “GABAergic” ;
 mtg_cluster_sc:cluster_size “64”^^xsd:int ;
.
~~~

~~~
:pCL38
 a skos:Concept ;
 mtg_cluster_sc:id “pCL38” ;
skos:broader :pCL81 ;
rdfs:label “TH-expressing-expressing cerebral cortex MTG GABAergic interneuron” ;
 mtg_cluster_sc:evidence “Inh L5-6 SST TH” ;
 go_sc:part_of uberon:UBERON_0002771 ;
 mtg_cluster_sc:enriched_in “cortical_layer5” ;
 mtg_cluster_sc:has_soma_location_in “cortical_layer5” ;
 mtg_cluster_sc:has_soma_location_in “cortical_layer6” ;
 mtg_cluster_sc:selectively_expresses hugo:HGNC_11782 ;
 mtg_cluster_sc:neuron_type “GABAergic” ;
 mtg_cluster_sc:cluster_size “27”^^xsd:int ;
.
~~~

~~~
:pCL39
 a skos:Concept ;
 mtg_cluster_sc:id “pCL39” ;
 skos:broader :pCL82 ;
 rdfs:label “GLP1R-expressing cerebral cortex MTG GABAergic interneuron” ;
 mtg_cluster_sc:evidence “Inh L5-6 GAD1 GLP1R” ;
 go_sc:part_of uberon:UBERON_0002771 ;
 mtg_cluster_sc:enriched_in “cortical_layer6” ;
 mtg_cluster_sc:has_soma_location_in “cortical_layer5” ;
 mtg_cluster_sc:has_soma_location_in “cortical_layer6” ;
 mtg_cluster_sc:selectively_expresses hugo:HGNC_4324 ;
 mtg_cluster_sc:neuron_type “GABAergic” ;
 mtg_cluster_sc:cluster_size “27”^^xsd:int ;
.
~~~

~~~
:pCL40
 a skos:Concept ;
 mtg_cluster_sc:id “pCL40” ;
 skos:broader :pCL82 ;
 rdfs:label “LGR5-expressing cerebral cortex MTG GABAergic interneuron” ;
 mtg_cluster_sc:evidence “Inh L5-6 PVALB LGR5” ;
 go_sc:part_of uberon:UBER0N_0002771 ;
 mtg_cluster_sc:enriched_in “cortical_layer5” ;
 mtg_cluster_sc:has_soma_location_in “cortical_layer5” ;
 mtg_cluster_sc:has_soma_location_in “cortical_layer6” ;
 mtg_cluster_sc:selectively_expresses hugo:HGNC_4504 ;
 mtg_cluster_sc:neuron_type “GABAergic” ;
 mtg_cluster_sc:cluster_size “52”^^xsd:int ;
.
~~~

~~~
:pCL41
 a skos:Concept ;
 mtg_cluster_sc:id “pCL41” ;
 skos:broader :pCL82 ;
 rdfs:label “MEPE-expressing cerebral cortex MTG GABAergic interneuron” ;
 mtg_cluster_sc:evidence “Inh L4-5 PVALB MEPE” ;
 go_sc:part_of uberon:UBER0N_0002771 ;
 mtg_cluster_sc:enriched_in “cortical_layer5” ;
 mtg_cluster_sc:has_soma_location_in “cortical_layer4” ;
 mtg_cluster_sc:has_soma_location_in “cortical_layer5” ;
 mtg_cluster_sc:selectively_expresses hugo:HGNC_13361 ;
 mtg_cluster_sc:neuron_type “GABAergic” ;
 mtg_cluster_sc:cluster_size “64”^^xsd:int ;
.
~~~

~~~
:pCL42
 a skos:Concept ;
 mtg_cluster_sc:id “pCL42” ;
 skos:broader :pCL82 ;
 rdfs:label “WFDC2-expressing cerebral cortex MTG GABAergic interneuron” ;
 mtg_cluster_sc:evidence “Inh L2-4 PVALB WFDC2” ;
 go_sc:part_of uberon:UBERON_0002771 ;
 mtg_cluster_sc:enriched_in “cortical_layer3” ;
 mtg_cluster_sc:has_soma_location_in “cortical_layer2” ;
 mtg_cluster_sc:has_soma_location_in “cortical_layer3” ;
 mtg_cluster_sc:has_soma_location_in “cortical_layer4” ;
 mtg_cluster_sc:selectively_expresses hugo:HGNC_15939 ;
 mtg_cluster_sc:neuron_type “GABAergic” ;
 mtg_cluster_sc:cluster_size “387”^^xsd:int ;
.
~~~

~~~
:pCL43
 a skos:Concept ;
 mtg_cluster_sc:id “pCL43” ; skos:broader :pCL82 ;
 rdfs:label “SULF1-expressing cerebral cortex MTG GABAergic interneuron” ;
 mtg_cluster_sc:evidence “Inh L4-6 PVALB SULF1” ;
 go_sc:part_of uberon:UBERON_0002771 ;
 mtg_cluster_sc:enriched_in “cortical_layer5” ;
 mtg_cluster_sc:has_soma_location_in “cortical_layer4” ;
 mtg_cluster_sc:has_soma_location_in “cortical_layer5” ;
 mtg_cluster_sc:has_soma_location_in “cortical_layer6” ;
 mtg_cluster_sc:selectively_expresses hugo:HGNC_20391 ;
 mtg_cluster_sc:neuron_type “GABAergic” ;
 mtg_cluster_sc:cluster_size “167”^^xsd:int ;
~~~

~~~
:pCL44 a skos:Concept ;
 mtg_cluster_sc:id “pCL44” ;
 skos:broader :pCL82 ;
 rdfs:label “SST|MIR548F2-expressing cerebral cortex MTG GABAergic interneuron” ;
 mtg_cluster_sc:evidence “Inh L5-6 SST MIR548F2” ;
 go_sc:part_of uberon:UBER0N_0002771 ;
 mtg_cluster_sc:enriched_in “cortical_layer5” ;
 mtg_cluster_sc:has_soma_location_in “cortical_layer5” ;
 mtg_cluster_sc:has_soma_location_in “cortical_layer6” ;
 mtg_cluster_sc:selectively_expresses hugo:HGNC_11329 ;
 mtg_cluster_sc:selectively_expresses hugo:HGNC_35306 ;
 mtg_cluster_sc:neuron_type “GABAergic” ;
 mtg_cluster_sc:cluster_size “80”^^xsd:int ;
.
~~~

~~~
:pCL45
 a skos:Concept ;
 mtg_cluster_sc:id “pCL45” ;
 skos:broader :pCL82 ;
 rdfs:label “SCUBE3-expressing cerebral cortex MTG GABAergic interneuron” ;
 mtg_cluster_sc:evidence “Inh L2-5 PVALB SCUBE3” ;
 go_sc:part_of uberon:UBER0N_0002771 ;
 mtg_cluster_sc:enriched_in “cortical_layer3” ;
 mtg_cluster_sc:has_soma_location_in “cortical_layer2” ;
 mtg_cluster_sc:has_soma_location_in “cortical_layer3” ;
 mtg_cluster_sc:has_soma_location_in “cortical_layer4” ;
 mtg_cluster_sc:has_soma_location_in “cortical_layer5” ;
 mtg_cluster_sc:selectively_expresses hugo:HGNC_13655 ;
 mtg_cluster_sc:neuron_type “GABAergic” ;
 mtg_cluster_sc:cluster_size “32”^^xsd:int ;
.
~~~

~~~
:pCL46
 a skos:Concept ;
 mtg_cluster_sc:id “pCL46” ;
 skos:broader :pCL77 ;
 rdfs:label “LAMP5|LTK-expressing cerebral cortex MTG Glutamatergic neuron” ;
 mtg_cluster_sc:evidence “Exc L2 LAMP5 LTK” ;
 go_sc:part_of uberon:UBERON_0002771 ;
 mtg_cluster_sc:enriched_in “cortical_layer2” ;
 mtg_cluster_sc:has_soma_location_in “cortical_layer2” ;
 mtg_cluster_sc:selectively_expresses hugo:HGNC_16097 ;
 mtg_cluster_sc:selectively_expresses hugo:HGNC_6721 ;
 mtg_cluster_sc:neuron_type “Glutamatergic” ;
 mtg_cluster_sc:cluster_size “812”^^xsd:int ;
.
~~~

~~~
:pCL47
 a skos:Concept ;
 mtg_cluster_sc:id “pCL47” ;
 skos:broader :pCL77 ;
 rdfs:label “LINC00507|GLP2R-expressing cerebral cortex MTG Glutamatergic neuron” ;
 mtg_cluster_sc:evidence “Exc L2-4 LINC00507 GLP2R” ;
 go_sc:part_of uberon:UBERON_0002771 ;
 mtg_cluster_sc:enriched_in “cortical_layer3” ;
 mtg_cluster_sc:has_soma_location_in “cortical_layer2” ;
 mtg_cluster_sc:has_soma_location_in “cortical_layer3” ;
 mtg_cluster_sc:has_soma_location_in “cortical_layer4” ;
 mtg_cluster_sc:selectively_expresses hugo:HGNC_43558 ;
 mtg_cluster_sc:selectively_expresses hugo:HGNC_4325 ;
 mtg_cluster_sc:neuron_type “Glutamatergic” ;
 mtg_cluster_sc:cluster_size “170”^^xsd:int ;
.
~~~

~~~
:pCL48
a skos:Concept ;
 mtg_cluster_sc:id “pCL48” ;
 skos:broader :pCL77 ;
 rdfs:label “LINC00507|FREM3-expressing cerebral cortex MTG Glutamatergic neuron” ;
 mtg_cluster_sc:evidence “Exc L2-3 LINC00507 FREM3” ;
 go_sc:part_of uberon:UBERON_0002771 ;
 mtg_cluster_sc:enriched_in “cortical_layer3” ;
 mtg_cluster_sc:has_soma_location_in “cortical_layer2” ;
 mtg_cluster_sc:has_soma_location_in “cortical_layer3” ;
 mtg_cluster_sc:selectively_expresses hugo:HGNC_43558 ;
 mtg_cluster_sc:selectively_expresses hugo:HGNC_25172 ;
 mtg_cluster_sc:neuron_type “Glutamatergic” ;
 mtg_cluster_sc:cluster_size “2284”^^xsd:int ;
.
~~~

~~~
:pCL49
 a skos:Concept ;
 mtg_cluster_sc:id “pCL49” ;
 skos:broader :pCL77 ;
 rdfs:label “THEMIS|C1QL3-expressing cerebral cortex MTG Glutamatergic neuron” ;
 mtg_cluster_sc:evidence “Exc L5-6 THEMIS C1QL3” ;
 go_sc:part_of uberon:UBERON_0002771 ;
 mtg_cluster_sc:enriched_in “cortical_layer6” ;
 mtg_cluster_sc:has_soma_location_in “cortical_layer5” ;
 mtg_cluster_sc:has_soma_location_in “cortical_layer6” ;
 mtg_cluster_sc:selectively_expresses hugo:HGNC_21569 ;
 mtg_cluster_sc:selectively_expresses hugo:HGNC_19359 ;
 mtg_cluster_sc:neuron_type “Glutamatergic” ;
 mtg_cluster_sc:cluster_size “1537”^^xsd:int ;
.
~~~

~~~
:pCL50
 a skos:Concept ;
 mtg_cluster_sc:id “pCL50” ;
 skos:broader :pCL77 ;
 rdfs:label “R0RB|CARM1P1-expressing cerebral cortex MTG Glutamatergic neuron” ;
 mtg_cluster_sc:evidence “Exc L3-4 R0RB CARM1P1” ;
 go_sc:part_of uberon:UBER0N_0002771 ;
 mtg_cluster_sc:enriched_in “cortical_layer3” ;
 mtg_cluster_sc:has_soma_location_in “cortical_layer3” ;
 mtg_cluster_sc:has_soma_location_in “cortical_layer4” ;
 mtg_cluster_sc:selectively_expresses hugo:HGNC_10259 ;
 mtg_cluster_sc:selectively_expresses hugo:HGNC_23392 ;
 mtg_cluster_sc:neuron_type “Glutamatergic” ;
 mtg_cluster_sc:cluster_size “280”^^xsd:int ;
.
~~~

~~~
:pCL51
 a skos:Concept ;
 mtg_cluster_sc:id “pCL51” ;
 skos:broader :pCL77 ;
 rdfs:label “R0RB|ESR1-expressing cerebral cortex MTG Glutamatergic neuron” ;
 mtg_cluster_sc:evidence “Exc L3-5 R0RB ESR1” ;
 go_sc:part_of uberon:UBER0N_0002771 ;
 mtg_cluster_sc:enriched_in “cortical_layer4” ;
 mtg_cluster_sc:has_soma_location_in “cortical_layer3” ;
 mtg_cluster_sc:has_soma_location_in “cortical_layer4” ;
 mtg_cluster_sc:has_soma_location_in “cortical_layer5” ;
 mtg_cluster_sc:selectively_expresses hugo:HGNC_10259 ;
 mtg_cluster_sc:selectively_expresses hugo:HGNC_3467 ;
 mtg_cluster_sc:neuron_type “Glutamatergic” ;
 mtg_cluster_sc:cluster_size “1428”^^xsd:int ;
.
~~~

~~~
:pCL52
 a skos:Concept ;
 mtg_cluster_sc:id “pCL52” ;
 skos:broader :pCL77 ;
 rdfs:label “RORB|COL22A1-expressing cerebral cortex MTG Glutamatergic neuron” ;
 mtg_cluster_sc:evidence “Exc L3-5 RORB COL22A1” ;
 go_sc:part_of uberon:UBERON_0002771 ;
 mtg_cluster_sc:enriched_in “cortical_layer4” ;
 mtg_cluster_sc:has_soma_location_in “cortical_layer3” ;
 mtg_cluster_sc:has_soma_location_in “cortical_layer4” ;
 mtg_cluster_sc:has_soma_location_in “cortical_layer5” ;
 mtg_cluster_sc:selectively_expresses hugo:HGNC_10259 ;
 mtg_cluster_sc:selectively_expresses hugo:HGNC_22989 ;
 mtg_cluster_sc:neuron_type “Glutamatergic” ;
 mtg_cluster_sc:cluster_size “160”^^xsd:int ;
.
~~~

~~~
:pCL53
 a skos:Concept ;
 mtg_cluster_sc:id “pCL53” ;
 skos:broader :pCL77 ;
 rdfs:label “RORB|FILIP1L-expressing cerebral cortex MTG Glutamatergic neuron” ;
 mtg_cluster_sc:evidence “Exc L3-5 RORB FILIP1L” ;
 go_sc:part_of uberon:UBERON_0002771 ;
 mtg_cluster_sc:enriched_in “cortical_layer4” ;
 mtg_cluster_sc:has_soma_location_in “cortical_layer3” ;
 mtg_cluster_sc:has_soma_location_in “cortical_layer4” ;
 mtg_cluster_sc:has_soma_location_in “cortical_layer5” ;
mtg_cluster_sc:selectively_expresses hugo:HGNC_10259 ;
 mtg_cluster_sc:selectively_expresses hugo:HGNC_24589 ;
 mtg_cluster_sc:neuron_type “Glutamatergic” ;
 mtg_cluster_sc:cluster_size “153”^^xsd:int ;
.
~~~

~~~
:pCL54
 a skos:Concept ;
 mtg_cluster_sc:id “pCL54” ;
 skos:broader :pCL77 ;
 rdfs:label “R0RB|TWIST2-expressing cerebral cortex MTG Glutamatergic neuron” ;
 mtg_cluster_sc:evidence “Exc L3-5 R0RB TWIST2” ;
 go_sc:part_of uberon:UBER0N_0002771 ;
 mtg_cluster_sc:enriched_in “cortical_layer4” ;
 mtg_cluster_sc:has_soma_location_in “cortical_layer3” ;
 mtg_cluster_sc:has_soma_location_in “cortical_layer4” ;
 mtg_cluster_sc:has_soma_location_in “cortical_layer5” ;
 mtg_cluster_sc:selectively_expresses hugo:HGNC_10259 ;
 mtg_cluster_sc:selectively_expresses hugo:HGNC_20670 ;
 mtg_cluster_sc:neuron_type “Glutamatergic” ;
 mtg_cluster_sc:cluster_size “93”^^xsd:int ;
.
~~~

~~~
:pCL55
 a skos:Concept ;
 mtg_cluster_sc:id “pCL55” ; skos:broader :pCL77 ;
 rdfs:label “R0RB|F0LH1B-expressing cerebral cortex MTG Glutamatergic neuron” ;
 mtg_cluster_sc:evidence “Exc L4-5 R0RB F0LH1B” ;
 go_sc:part_of uberon:UBER0N_0002771 ;
 mtg_cluster_sc:enriched_in “cortical_layer4” ;
mtg_cluster_sc:has_soma_location_in “cortical_layer4” ;
 mtg_cluster_sc:has_soma_location_in “cortical_layer5” ;
 mtg_cluster_sc:selectively_expresses hugo:HGNC_10259 ;
 mtg_cluster_sc:selectively_expresses hugo:HGNC_13636 ;
 mtg_cluster_sc:neuron_type “Glutamatergic” ;
 mtg_cluster_sc:cluster_size “870”^^xsd:int ;
.
~~~

~~~
:pCL56
 a skos:Concept ;
 mtg_cluster_sc:id “pCL56” ;
 skos:broader :pCL77 ;
 rdfs:label “RORB|SEMA3E-expressing cerebral cortex MTG Glutamatergic neuron” ;
 mtg_cluster_sc:evidence “Exc L4-6 RORB SEMA3E” ;
 go_sc:part_of uberon:UBERON_0002771 ;
 mtg_cluster_sc:enriched_in “cortical_layer5” ;
 mtg_cluster_sc:has_soma_location_in “cortical_layer4” ;
 mtg_cluster_sc:has_soma_location_in “cortical_layer5” ;
 mtg_cluster_sc:has_soma_location_in “cortical_layer6” ;
 mtg_cluster_sc:selectively_expresses hugo:HGNC_10259 ;
 mtg_cluster_sc:selectively_expresses hugo:HGNC_10727 ;
 mtg_cluster_sc:neuron_type “Glutamatergic” ;
 mtg_cluster_sc:cluster_size “777”^^xsd:int ;
.
~~~

~~~
:pCL57
 a skos:Concept ;
 mtg_cluster_sc:id “pCL57” ;
 skos:broader :pCL77 ;
rdfs:label “RORB|DAPK2-expressing cerebral cortex MTG Glutamatergic neuron” ;
 mtg_cluster_sc:evidence “Exc L4-5 RORB DAPK2” ;
 go_sc:part_of uberon:UBER0N_0002771 ;
 mtg_cluster_sc:enriched_in “cortical_layer4” ;
 mtg_cluster_sc:has_soma_location_in “cortical_layer4” ;
 mtg_cluster_sc:has_soma_location_in “cortical_layer5” ;
 mtg_cluster_sc:selectively_expresses hugo:HGNC_10259 ;
 mtg_cluster_sc:selectively_expresses hugo:HGNC_2675 ;
 mtg_cluster_sc:neuron_type “Glutamatergic” ;
 mtg_cluster_sc:cluster_size “173”^^xsd:int ;
.
~~~

~~~
:pCL58
 a skos:Concept ;
 mtg_cluster_sc:id “pCL58” ;
 skos:broader :pCL77 ;
 rdfs:label “R0RB|TTC12-expressing cerebral cortex MTG Glutamatergic neuron” ;
 mtg_cluster_sc:evidence “Exc L5-6 R0RB TTC12” ;
 go_sc:part_of uberon:UBER0N_0002771 ;
 mtg_cluster_sc:enriched_in “cortical_layer5” ;
 mtg_cluster_sc:has_soma_location_in “cortical_layer5” ;
 mtg_cluster_sc:has_soma_location_in “cortical_layer6” ;
 mtg_cluster_sc:selectively_expresses hugo:HGNC_10259 ;
 mtg_cluster_sc:selectively_expresses hugo:HGNC_23700 ;
 mtg_cluster_sc:neuron_type “Glutamatergic” ;
 mtg_cluster_sc:cluster_size “167”^^xsd:int ;
.
~~~

~~~
:pCL59
 a skos:Concept ;
 mtg_cluster_sc:id “pCL59” ;
 skos:broader :pCL77 ;
 rdfs:label “R0RB|C1R-expressing cerebral cortex MTG Glutamatergic neuron” ;
 mtg_cluster_sc:evidence “Exc L4-6 RORB C1R” ;
 go_sc:part_of uberon:UBERON_0002771 ;
 mtg_cluster_sc:enriched_in “cortical_layer5” ;
 mtg_cluster_sc:has_soma_location_in “cortical_layer4” ;
 mtg_cluster_sc:has_soma_location_in “cortical_layer5” ;
 mtg_cluster_sc:has_soma_location_in “cortical_layer6” ;
 mtg_cluster_sc:selectively_expresses hugo:HGNC_10259 ;
 mtg_cluster_sc:selectively_expresses hugo:HGNC_1246 ;
 mtg_cluster_sc:neuron_type “Glutamatergic” ;
 mtg_cluster_sc:cluster_size “160”^^xsd:int ;
.
~~~

~~~
:pCL60
 a skos:Concept ;
 mtg_cluster_sc:id “pCL60” ;
 skos:broader :pCL77 ;
 rdfs:label “FEZF2|SCN4B-expressing cerebral cortex MTG Glutamatergic neuron” ;
 mtg_cluster_sc:evidence “Exc L4-5 FEZF2 SCN4B” ;
 go_sc:part_of uberon:UBERON_0002771 ;
 mtg_cluster_sc:enriched_in “cortical_layer5” ;
 mtg_cluster_sc:has_soma_location_in “cortical_layer4” ;
 mtg_cluster_sc:has_soma_location_in “cortical_layer5” ;
 mtg_cluster_sc:selectively_expresses hugo:HGNC_13506 ;
 mtg_cluster_sc:selectively_expresses hugo:HGNC_10592 ;
 mtg_cluster_sc:neuron_type “Glutamatergic” ;
 mtg_cluster_sc:cluster_size “25”^^xsd:int ;
.
~~~

~~~
:pCL61
 a skos:Concept ;
 mtg_cluster_sc:id “pCL61” ;
skos:broader :pCL77 ;
rdfs:label “THEMIS|DCSTAMP-expressing cerebral cortex MTG Glutamatergic neuron” ;
 mtg_cluster_sc:evidence “Exc L5-6 THEMIS DCSTAMP” ;
 go_sc:part_of uberon:UBERON_0002771 ;
 mtg_cluster_sc:enriched_in “cortical_layer5” ;
 mtg_cluster_sc:has_soma_location_in “cortical_layer5” ;
 mtg_cluster_sc:has_soma_location_in “cortical_layer6” ;
 mtg_cluster_sc:selectively_expresses hugo:HGNC_21569 ;
 mtg_cluster_sc:selectively_expresses hugo:HGNC_18549 ;
 mtg_cluster_sc:neuron_type “Glutamatergic” ;
 mtg_cluster_sc:cluster_size “53”^^xsd:int ;
.
~~~

~~~
:pCL62
 a skos:Concept ;
 mtg_cluster_sc:id “pCL62” ;
 skos:broader :pCL77 ;
 rdfs:label “THEMIS|CRABP1-expressing cerebral cortex MTG Glutamatergic neuron” ;
 mtg_cluster_sc:evidence “Exc L5-6 THEMIS CRABP1” ; go_sc:part_of uberon:UBERON_0002771 ;
 mtg_cluster_sc:enriched_in “cortical_layer5” ;
 mtg_cluster_sc:has_soma_location_in “cortical_layer5” ;
 mtg_cluster_sc:has_soma_location_in “cortical_layer6” ;
 mtg_cluster_sc:selectively_expresses hugo:HGNC_21569 ;
 mtg_cluster_sc:selectively_expresses hugo:HGNC_2338 ;
 mtg_cluster_sc:neuron_type “Glutamatergic” ;
 mtg_cluster_sc:cluster_size “147”^^xsd:int ;
.
~~~

~~~
:pCL63 a skos:Concept ;
 mtg_cluster_sc:id “pCL63” ;
 skos:broader :pCL77 ;
 rdfs:label “THEMIS|FGF10-expressing cerebral cortex MTG Glutamatergic neuron” ;
 mtg_cluster_sc:evidence “Exc L5-6 THEMIS FGF10” ;
 go_sc:part_of uberon:UBER0N_0002771 ;
 mtg_cluster_sc:enriched_in “cortical_layer5” ;
 mtg_cluster_sc:has_soma_location_in “cortical_layer5” ;
 mtg_cluster_sc:has_soma_location_in “cortical_layer6” ;
 mtg_cluster_sc:selectively_expresses hugo:HGNC_21569 ;
 mtg_cluster_sc:selectively_expresses hugo:HGNC_3666 ;
 mtg_cluster_sc:neuron_type “Glutamatergic” ;
 mtg_cluster_sc:cluster_size “78”^^xsd:int ;
.
~~~

~~~
:pCL64
 a skos:Concept ;
 mtg_cluster_sc:id “pCL64” ;
 skos:broader :pCL77 ;
 rdfs:label “FEZF2|IL26-expressing cerebral cortex MTG Glutamatergic neuron” ;
 mtg_cluster_sc:evidence “Exc L4-6 FEZF2 IL26” ;
 go_sc:part_of uberon:UBER0N_0002771 ;
 mtg_cluster_sc:enriched_in “cortical_layer5” ;
 mtg_cluster_sc:has_soma_location_in “cortical_layer4” ;
 mtg_cluster_sc:has_soma_location_in “cortical_layer5” ;
 mtg_cluster_sc:has_soma_location_in “cortical_layer6” ;
 mtg_cluster_sc:selectively_expresses hugo:HGNC_13506 ;
 mtg_cluster_sc:selectively_expresses hugo:HGNC_17119 ;
 mtg_cluster_sc:neuron_type “Glutamatergic” ;
 mtg_cluster_sc:cluster_size “344”^^xsd:int ;
.
~~~

~~~
:pCL65
 a skos:Concept ;
 mtg_cluster_sc:id “pCL65” ; skos:broader :pCL77 ;
 rdfs:label “FEZF2|AB0-expressing cerebral cortex MTG Glutamatergic neuron” ;
 mtg_cluster_sc:evidence “Exc L5-6 FEZF2 AB0” ;
 go_sc:part_of uberon:UBER0N_0002771 ;
 mtg_cluster_sc:enriched_in “cortical_layer6” ;
 mtg_cluster_sc:has_soma_location_in “cortical_layer5” ;
 mtg_cluster_sc:has_soma_location_in “cortical_layer6” ;
 mtg_cluster_sc:selectively_expresses hugo:HGNC_13506 ;
 mtg_cluster_sc:selectively_expresses hugo:HGNC_79 ;
 mtg_cluster_sc:neuron_type “Glutamatergic” ;
 mtg_cluster_sc:cluster_size “373”^^xsd:int ;
.
~~~

~~~
:pCL66
 a skos:Concept ;
 mtg_cluster_sc:id “pCL66” ;
 skos:broader :pCL77 ;
 rdfs:label “FEZF2|SCUBE1-expressing cerebral cortex MTG Glutamatergic neuron” ;
 mtg_cluster_sc:evidence “Exc L6 FEZF2 SCUBE1” ;
 go_sc:part_of uberon:UBER0N_0002771 ;
 mtg_cluster_sc:enriched_in “cortical_layer6” ;
 mtg_cluster_sc:has_soma_location_in “cortical_layer6” ;
 mtg_cluster_sc:selectively_expresses hugo:HGNC_13506 ;
 mtg_cluster_sc:selectively_expresses hugo:HGNC_13441 ;
 mtg_cluster_sc:neuron_type “Glutamatergic” ;
 mtg_cluster_sc:cluster_size “52”^^xsd:int ;
.
~~~

~~~
:pCL67
 a skos:Concept ;
 mtg_cluster_sc:id “pCL67” ;
 skos:broader :pCL77 ;
 rdfs:label “IL15-expressing cerebral cortex MTG Glutamatergic neuron” ;
 mtg_cluster_sc:evidence “Exc L5-6 SLC17A7 IL15” ;
 go_sc:part_of uberon:UBERON_0002771 ;
 mtg_cluster_sc:enriched_in “cortical_layer6” ;
 mtg_cluster_sc:has_soma_location_in “cortical_layer5” ;
 mtg_cluster_sc:has_soma_location_in “cortical_layer6” ;
 mtg_cluster_sc:selectively_expresses hugo:HGNC_5977 ;
 mtg_cluster_sc:neuron_type “Glutamatergic” ;
 mtg_cluster_sc:cluster_size “56”^^xsd:int ;
.
~~~

~~~
:pCL68
 a skos:Concept ;
 mtg_cluster_sc:id “pCL68” ;
 skos:broader :pCL77 ;
 rdfs:label “FEZF2|OR2T8-expressing cerebral cortex MTG Glutamatergic neuron” ;
 mtg_cluster_sc:evidence “Exc L6 FEZF2 OR2T8” ;
 go_sc:part_of uberon:UBERON_0002771 ;
 mtg_cluster_sc:enriched_in “cortical_layer6” ;
 mtg_cluster_sc:has_soma_location_in “cortical_layer6” ;
 mtg_cluster_sc:selectively_expresses hugo:HGNC_13506 ;
 mtg_cluster_sc:selectively_expresses hugo:HGNC_15020 ;
 mtg_cluster_sc:neuron_type “Glutamatergic” ;
 mtg_cluster_sc:cluster_size “19”^^xsd:int ;
.
~~~

~~~
:pCL69
a skos:Concept ;
 mtg_cluster_sc:id “pCL69” ;
 skos:broader :pCL77 ;
 rdfs:label “FEZF2|EFTUD1P1-expressing cerebral cortex MTG Glutamatergic neuron” ;
 mtg_cluster_sc:evidence “Exc L5-6 FEZF2 EFTUD1P1” ;
 go_sc:part_of uberon:UBER0N_0002771 ;
 mtg_cluster_sc:enriched_in “cortical_layer6” ;
 mtg_cluster_sc:has_soma_location_in “cortical_layer5” ;
 mtg_cluster_sc:has_soma_location_in “cortical_layer6” ;
 mtg_cluster_sc:selectively_expresses hugo:HGNC_13506 ;
 mtg_cluster_sc:selectively_expresses hugo:HGNC_31739 ;
 mtg_cluster_sc:neuron_type “Glutamatergic” ;
 mtg_cluster_sc:cluster_size “314”^^xsd:int ;
.
~~~

~~~
:pCL70
 a skos:Concept ;
 mtg_cluster_sc:id “pCL70” ;
 skos:broader :pCL83 ;
 rdfs:label “PDGFRA-expressing MTG 0ligodendrocyte precursor cell” ;
 mtg_cluster_sc:evidence “0PC L1-6 PDGFRA” ;
 go_sc:part_of uberon:UBER0N_0002771 ;
 mtg_cluster_sc:enriched_in “cortical_layer4” ;
 mtg_cluster_sc:has_soma_location_in “cortical_layer1” ;
 mtg_cluster_sc:has_soma_location_in “cortical_layer2” ;
 mtg_cluster_sc:has_soma_location_in “cortical_layer3” ;
 mtg_cluster_sc:has_soma_location_in “cortical_layer4” ;
 mtg_cluster_sc:has_soma_location_in “cortical_layer5” ;
 mtg_cluster_sc:has_soma_location_in “cortical_layer6” ;
 mtg_cluster_sc:selectively_expresses hugo:HGNC_8803 ;
mtg_cluster_sc:cluster_size “238”^^xsd:int ;
.
~~~

~~~
:pCL71
 a skos:Concept ;
 mtg_cluster_sc:id “pCL71” ;
 skos:broader :pCL89 ;
 rdfs:label “SLC14A1-expressing MTG astrocyte” ;
 mtg_cluster_sc:evidence “Astro L1 -6 FGFR3 SLC14A1” ;
 go_sc:part_of uberon:UBERON_0002771 ;
 mtg_cluster_sc:enriched_in “cortical_layer3” ;
 mtg_cluster_sc:has_soma_location_in “cortical_layer1” ;
 mtg_cluster_sc:has_soma_location_in “cortical_layer2” ;
 mtg_cluster_sc:has_soma_location_in “cortical_layer3” ;
 mtg_cluster_sc:has_soma_location_in “cortical_layer4” ;
 mtg_cluster_sc:has_soma_location_in “cortical_layer5” ;
 mtg_cluster_sc:has_soma_location_in “cortical_layer6” ;
 mtg_cluster_sc:selectively_expresses hugo:HGNC_10918 ;
 mtg_cluster_sc:cluster_size “230”^^xsd:int ;
.
~~~

~~~
:pCL72
 a skos:Concept ;
 mtg_cluster_sc:id “pCL72” ;
 skos:broader :pCL89 ;
 rdfs:label “GFAP-expressing MTG astrocyte” ;
 mtg_cluster_sc:evidence “Astro L1-2 FGFR3 GFAP” ; go_sc:part_of uberon:UBERON_0002771 ;
 mtg_cluster_sc:enriched_in “cortical_layer2” ;
 mtg_cluster_sc:has_soma_location_in “cortical_layer1” ;
 mtg_cluster_sc:has_soma_location_in “cortical_layer2” ;
mtg_cluster_sc:selectively_expresses hugo:HGNC_4235 ;
 mtg_cluster_sc:cluster_size “61”^^xsd:int ;
.
~~~

~~~
:pCL73
 a skos:Concept ;
 mtg_cluster_sc:id “pCL73” ;
 skos:broader :pCL86 ;
 rdfs:label “OPALIN-expressing MTG Oligodendrocyte” ;
 mtg_cluster_sc:evidence “Oligo L1 -6 OPALIN” ;
 go_sc:part_of uberon:UBERON_0002771 ;
 mtg_cluster_sc:enriched_in “cortical_layer5” ;
 mtg_cluster_sc:has_soma_location_in “cortical_layer1” ;
 mtg_cluster_sc:has_soma_location_in “cortical_layer2” ;
 mtg_cluster_sc:has_soma_location_in “cortical_layer3” ;
 mtg_cluster_sc:has_soma_location_in “cortical_layer4” ;
 mtg_cluster_sc:has_soma_location_in “cortical_layer5” ;
 mtg_cluster_sc:has_soma_location_in “cortical_layer6” ;
 mtg_cluster_sc:selectively_expresses hugo:HGNC_20707 ;
 mtg_cluster_sc:cluster_size “313”^^xsd:int ;
.
~~~

~~~
:pCL74
 a skos:Concept ;
 mtg_cluster_sc:id “pCL74” ; skos:broader :pCL87 ;
 rdfs:label “NOSTRIN-expressing cerebral cortex MTG endothelial cell” ;
 mtg_cluster_sc:evidence “Endo L2-6 NOSTRIN” ;
 go_sc:part_of uberon:UBERON_0002771 ;
 mtg_cluster_sc:enriched_in “cortical_layer4” ;
 mtg_cluster_sc:has_soma_location_in “cortical_layer2” ;
 mtg_cluster_sc:has_soma_location_in “cortical_layer3” ;
 mtg_cluster_sc:has_soma_location_in “cortical_layer4” ;
 mtg_cluster_sc:has_soma_location_in “cortical_layer5” ;
 mtg_cluster_sc:has_soma_location_in “cortical_layer6” ;
 mtg_cluster_sc:selectively_expresses hugo:HGNC_20203 ;
 mtg_cluster_sc:cluster_size “9”^^xsd:int ;
.
~~~

~~~
:pCL75
 a skos:Concept ;
 mtg_cluster_sc:id “pCL75” ;
 skos:broader :pCL88 ;
 rdfs:label “TYR0BP-expressing MTG Microglial cell” ;
 mtg_cluster_sc:evidence “Micro L1-3 TYR0BP” ;
 go_sc:part_of uberon:UBER0N_0002771 ;
 mtg_cluster_sc:enriched_in “cortical_layer3” ;
 mtg_cluster_sc:has_soma_location_in “cortical_layer1” ;
 mtg_cluster_sc:has_soma_location_in “cortical_layer2” ;
 mtg_cluster_sc:has_soma_location_in “cortical_layer3” ;
 mtg_cluster_sc:selectively_expresses hugo:HGNC_12449 ;
 mtg_cluster_sc:cluster_size “63”^^xsd:int ;
.
~~~

~~~
:pCL76
 a skos:Concept ;
 mtg_cluster_sc:id “pCL76” ;
 skos:broader :pCL90 ;
 rdfs:label “GAD1-expressing cerebral cortex MTG GABAergic interneuron” ;
 go_sc:part_of uberon:UBER0N_0002771 ;
 mtg_cluster_sc:selectively_expresses hugo:HGNC_4092 ;
 mtg_cluster_sc:neuron_type “GABAergic” ;
.
~~~

~~~
:pCL77
 a skos:Concept ;
 mtg_cluster_sc:id “pCL77” ;
 skos:broader :pCL91 ;
 rdfs:label “SLC17A7-expressing MTG Glutamatergic neuron” ;
 go_sc:part_of uberon:UBER0N_0002771 ;
 mtg_cluster_sc:selectively_expresses hugo:HGNC_16704 ;
 mtg_cluster_sc:neuron_type “Glutamatergic” ;
~~~

~~~
:pCL78
 a skos:Concept ;
 mtg_cluster_sc:id “pCL78” ;
 skos:broader :pCL76 ;
 rdfs:label “ADARB2-expressing cerebral cortex MTG GABAergic interneuron” ;
 go_sc:part_of uberon:UBER0N_0002771 ;
 mtg_cluster_sc:selectively_expresses hugo:HGNC_227 ;
 mtg_cluster_sc:neuron_type “GABAergic” ;
.
~~~

~~~
:pCL79
 a skos:Concept ;
 mtg_cluster_sc:id “pCL79” ; skos:broader :pCL76 ;
 rdfs:label “LHX6-expressing cerebral cortex MTG GABAergic interneuron” ;
 go_sc:part_of uberon:UBER0N_0002771 ;
 mtg_cluster_sc:selectively_expresses hugo:HGNC_21735 ;
 mtg_cluster_sc:neuron_type “GABAergic” ;
.
~~~

~~~
:pCL80
a skos:Concept ;
 mtg_cluster_sc:id “pCL80” ;
 skos:broader :pCL78 ;
 rdfs:label “VIP-expressing cerebral cortex MTG GABAergic interneuron” ;
 go_sc:part_of uberon:UBERON_0002771 ;
 mtg_cluster_sc:selectively_expresses hugo:HGNC_12693 ;
 mtg_cluster_sc:neuron_type “GABAergic” ;
.
~~~

~~~
:pCL81
 a skos:Concept ;
 mtg_cluster_sc:id “pCL81” ;
 skos:broader :pCL79 ;
rdfs:label “SST-expressing cerebral cortex MTG GABAergic interneuron” ;
 go_sc:part_of uberon:UBERON_0002771 ;
 mtg_cluster_sc:selectively_expresses hugo:HGNC_11329 ;
 mtg_cluster_sc:neuron_type “GABAergic” ;
.
~~~

~~~
:pCL82
 a skos:Concept ;
 mtg_cluster_sc:id “pCL82” ; skos:broader :pCL79 ;
 rdfs:label “PVALB-expressing cerebral cortex MTG GABAergic interneuron” ;
 go_sc:part_of uberon:UBERON_0002771 ;
 mtg_cluster_sc:selectively_expresses hugo:HGNC_9704 ;
 mtg_cluster_sc:neuron_type “GABAergic” ;
.
~~~

~~~
:pCL83
 a skos:Concept ;
 mtg_cluster_sc:id “pCL83” ;
 skos:broader cl:CL_0002453 ;
 rdfs:label “MTG 0ligodendrocyte precursor cell” ;
 go_sc:part_of uberon:UBER0N_0002771 ;
.
~~~

~~~
:pCL84
 a skos:Concept ;
 mtg_cluster_sc:id “pCL84” ;
 skos:broader cl:CL_0002605 ;
 rdfs:label “MTG Astrocyte of the cerebral cortex” ;
 go_sc:part_of uberon:UBER0N_0002771 ;
~~~

~~~
:pCL86 a skos:Concept ;
 mtg_cluster_sc:id “pCL86” ;
 skos:broader cl:CL_0000128 ; rdfs:label “MTG 0ligodendrocyte” ;
 go_sc:part_of uberon:UBER0N_0002771 ;
.
~~~

~~~
:pCL87
 a skos:Concept ;
 mtg_cluster_sc:id “pCL87” ;
 skos:broader cl:CL_1001602 ; rdfs:label “cerebral cortex MTG endothelial cell” ;
 go_sc:part_of uberon:UBER0N_0002771 ;
~~~

~~~
:pCL88 a skos:Concept ;
 mtg_cluster_sc:id “pCL88” ;
 skos:broader cl:CL_0000129 ;
rdfs:label “MTG Microglial cell” ;
 go_sc:part_of uberon:UBERON_0002771 ;
.
~~~

~~~
:pCL89 a skos:Concept ;
 mtg_cluster_sc:id “pCL89” ;
 skos:broader :pCL84 ;
 rdfs:label “FGFR3-expressing MTG astrocyte” ;
 go_sc:part_of uberon:UBERON_0002771 ;
 mtg_cluster_sc:selectively_expresses hugo:HGNC_3690 ;
.
~~~

~~~
:pCL90
 a skos:Concept ;
 mtg_cluster_sc:id “pCL90” ;
 skos:broader cl:CL_0010011 ;
 rdfs:label “cerebral cortex MTG GABAergic interneuron” ;
 go_sc:part_of uberon:UBERON_0002771 ;
 mtg_cluster_sc:selectively_expresses hugo:HGNC_4092 ;
 mtg_cluster_sc:neuron_type “GABAergic” ;
.
~~~

~~~
:pCL91
 a skos:Concept ;
 mtg_cluster_sc:id “pCL91” ;
 skos:broader cl:CL_0000679 ;
 rdfs:label “MTG Glutamatergic neuron” ;
 go_sc:part_of uberon:UBERON_0002771 ;
 mtg_cluster_sc:selectively_expresses hugo:HGNC_16704 ;
 mtg_cluster_sc:neuron_type “Glutamatergic” ;
~~~

## References

1. Glasser, M. F. et al. A multi-modal parcellation of human cerebral cortex. Nature 536, 171- 178 (2016).

2. Nieuwenhuys, R. The myeloarchitectonic studies on the human cerebral cortex of the Vogt-Vogt school, and their significance for the interpretation of functional neuroimaging data. Brain Struct Funct 218, 303-52 (2013).

3. Essen, D. C. V., Glasser, M. F., Dierker, D. L., Harwell, J. & Coalson, T. Parcellations and Hemispheric Asymmetries of Human Cerebral Cortex Analyzed on Surface-Based Atlases. Cerebral Cortex 22, 2241-2262 (2011).

4. Azevedo, F. A. C. et al. Equal numbers of neuronal and nonneuronal cells make the human brain an isometrically scaled-up primate brain. The Journal of Comparative Neurology 513, 532-541 (2009).

5. Herculano-Houzel, S., Mota, B. & Lent, R. Cellular scaling rules for rodent brains. Proc Natl Acad Sci U S A 103, 12138-43 (2006).

6. Azevedo, F. A. et al. Equal numbers of neuronal and nonneuronal cells make the human brain an isometrically scaled-up primate brain. J Comp Neurol 513, 532-41 (2009).

7. Geschwind, D. H. & Rakic, P. Cortical evolution: judge the brain by its cover. Neuron 80, 633- 47 (2013).

8. DeFelipe, J. The evolution of the brain, the human nature of cortical circuits, and intellectual creativity. Front Neuroanat 5, 29 (2011).

9. Cajal, S. Ramón y. La Textura del Sistema Nerviosa del Hombre y los Vertebrados. (1904).

10. Nó, R. Lorente de. La corteza cerebral del ratón. Trab. Lab. Invest. Bio. (Madrid) 20, (1922).

11. Poorthuis, R. B. et al. Rapid Neuromodulation of Layer 1 Interneurons in Human Neocortex. Cell Rep 23, 951-958 (2018).

12. Eyal, G. et al. Unique membrane properties and enhanced signal processing in human neocortical neurons. Elife 5, (2016).

13. Szegedi, V. et al. Plasticity in Single Axon Glutamatergic Connection to GABAergic Interneurons Regulates Complex Events in the Human Neocortex. PLoS Biol 14, e2000237 (2016).

14. DeFelipe, J. Types of neurons, synaptic connections and chemical characteristics of cells immunoreactive for calbindin-D28K, parvalbumin and calretinin in the neocortex. J Chem Neuroanat 14, 1-19 (1997).

15. Benavides-Piccione, R., Ballesteros-Yáñez, I., DeFelipe, J. & Yuste, R. Cortical area and species differences in dendritic spine morphology. J Neurocytol 31, 337-46 (2002).

16. Gabbott, P. L. Subpial Fan Cell - A Class of Calretinin Neuron in Layer 1 of Adult Monkey Prefrontal Cortex. Front Neuroanat 10, 28 (2016).

17. Oberheim, N. A. et al. Uniquely hominid features of adult human astrocytes. J Neurosci 29, 3276-87 (2009).

18. Hill, R. S. & Walsh, C. A. Molecular insights into human brain evolution. Nature 437, 64-7 (2005).

19. Boldog, E. et al. Transcriptomic and morphophysiological evidence for a specialized human cortical GABAergic cell type. bioRxiv (2017). doi:10.1101/216085

20. Zeng, H. et al. Large-scale cellular-resolution gene profiling in human neocortex reveals species-specific molecular signatures. Cell 149, 483-96 (2012).

21. Bakken, T. E. et al. A comprehensive transcriptional map of primate brain development. Nature 535, 367-75 (2016).

22. Hawrylycz, M. et al. Canonical genetic signatures of the adult human brain. Nat Neurosci 18, 1832-44 (2015).

23. Miller, J. A. et al. Transcriptional landscape of the prenatal human brain. Nature 508, 199- 206 (2014).

24. Ecker, J. R. et al. The BRAIN Initiative Cell Census Consortium: Lessons Learned toward Generating a Comprehensive Brain Cell Atlas. Neuron 96, 542-557 (2017).

25. Regev, A. et al. The Human Cell Atlas. Elife 6, (2017).

26. Tasic, B. et al. Shared and distinct transcriptomic cell types across neocortical areas. bioRxiv (2017). doi:10.1101/229542

27. Tasic, B. et al. Adult mouse cortical cell taxonomy revealed by single cell transcriptomics.Nat Neurosci 19, 335–46 (2016).

28. Zeisel, A. et al. Brain structure. Cell types in the mouse cortex and hippocampus revealed by single-cell RNA-seq. Science 347, 1138–42 (2015).

29. Darmanis, S. et al. A survey of human brain transcriptome diversity at the single cell level. Proc Natl Acad Sci U S A 112, 7285–90 (2015).

30. Krishnaswami, S. R. et al. Using single nuclei for RNA-seq to capture the transcriptome of postmortem neurons. Nat Protoc 11, 499–524 (2016).

31. Lake, B. B. et al. Neuronal subtypes and diversity revealed by single-nucleus RNA sequencing of the human brain. Science 352, 1586–90 (2016).

32. Habib, N. et al. Massively parallel single-nucleus RNA-seq with DroNc-seq. Nat Methods 14, 955–958 (2017).

33. Bakken, T. E. et al. Equivalent high-resolution identification of neuronal cell types with singlenucleus and single-cell RNA-sequencing. bioRxiv (2017). doi:10.1101/239749

34. Lake, B. B. et al. A comparative strategy for single-nucleus and single-cell transcriptomes confirms accuracy in predicted cell-type expression from nuclear RNA. Sci Rep 7, 6031 (2017).

35. Lake, B. B. et al. Integrative single-cell analysis of transcriptional and epigenetic states in the human adult brain. Nat Biotechnol 36, 70–80 (2018).

36. DeFelipe, J., Alonso-Nanclares, L. & Arellano, J. I. Microstructure of the neocortex: comparative aspects. J Neurocytol 31, 299–316 (2002).

37. Bakken, T. et al. Cell type discovery and representation in the era of high-content single cell phenotyping. BMC Bioinformatics 18, 559 (2017).

38. Bakken, T. E. et al. Spatiotemporal dynamics of the postnatal developing primate brain transcriptome. Human Molecular Genetics 24, 4327–4339 (2015).

39. Werner, M. S. et al. Chromatin-enriched lncRNAs can act as cell-type specific activators of proximal gene transcription. Nat Struct Mol Biol 24, 596–603 (2017).

40. Von Economo, C. Cellular structure of the human cerebral cortex. (Karger Medical and Scientific Publishers, 2009).

41. Kalmbach, B. et al. h-channels contribute to divergent electrophysiological properties of supragranular pyramidal neurons in human versus mouse cerebral cortex. bioRxiv (2018). doi:10.1101/312298

42. Cytosplore: Interactive Immune Cell Phenotyping for Large Single-Cell Datasets. Computer Graphics Forum 35, (2016).

43. Hollt, T. et al. CyteGuide: Visual Guidance for Hierarchical Single-Cell Analysis. IEEE Trans Vis Comput Graph 24, 739–748 (2018).

44. Lee, S., Hjerling-Leffler, J., Zagha, E., Fishell, G. & Rudy, B. The largest group of superficial neocortical GABAergic interneurons expresses ionotropic serotonin receptors. J Neurosci 30, 16796-808 (2010).

45. Hansen, D. V. et al. Non-epithelial stem cells and cortical interneuron production in the human ganglionic eminences. Nat Neurosci 16, 1576-87 (2013).

46. Ma, T. et al. Subcortical origins of human and monkey neocortical interneurons. Nat Neurosci 16, 1588-97 (2013).

47. Tasic, B. et al. Shared and distinct transcriptomic cell types across neocortical areas. bioRxiv(2017). doi:10.1101/229542

48. Shah, B. P. et al. MC4R-expressing glutamatergic neurons in the paraventricular hypothalamus regulate feeding and are synaptically connected to the parabrachial nucleus.Proc Natl Acad Sci U S A 111, 13193-8 (2014).

49. Horstmann, A. et al. Common genetic variation near MC4R has a sex-specific impact on human brain structure and eating behavior. PLoS One 8, e74362 (2013).

50. Raghanti, M. A. et al. Neuropeptide Y-immunoreactive neurons in the cerebral cortex of humans and other haplorrhine primates. Am J Primatol 75, 415-24 (2013).

51. Xu, X., Roby, K. D. & Callaway, E. M. Immunochemical characterization of inhibitory mouse cortical neurons: three chemically distinct classes of inhibitory cells. J Comp Neurol 518, 389- 404 (2010).

52. Paul, A. et al. Transcriptional Architecture of Synaptic Communication Delineates GABAergic Neuron Identity. Cell 171, 522-539.e20 (2017).

53. Marques, S. et al. Oligodendrocyte heterogeneity in the mouse juvenile and adult central nervous system. Science 352, 1326-1329 (2016).

54. Zhang, Y. et al. Purification and Characterization of Progenitor and Mature Human Astrocytes Reveals Transcriptional and Functional Differences with Mouse. Neuron 89, 37-53 (2016).

55. Sosunov, A. A. et al. Phenotypic heterogeneity and plasticity of isocortical and hippocampal astrocytes in the human brain. J Neurosci 34, 2285-98 (2014).

56. Butler, A., Hoffman, P., Smibert, P., Papalexi, E. & Satija, R. Integrating single-cell transcriptomic data across different conditions, technologies, and species. Nat Biotechnol 36, 411-420 (2018).

57. Kilduff, T. S., Cauli, B. & Gerashchenko, D. Activation of cortical interneurons during sleep: an anatomical link to homeostatic sleep regulation? Trends Neurosci 34, 10-9 (2011).

58. He, M. et al. Strategies and Tools for Combinatorial Targeting of GABAergic Neurons in Mouse Cerebral Cortex. Neuron 92, 555 (2016).

59. Sorensen, S. A. et al. Correlated Gene Expression and Target Specificity Demonstrate Excitatory Projection Neuron Diversity. Cerebral Cortex 25, 433-449 (2013).

60. Belichenko, P. V., Vogt, W. D. M., Myklóssy, J. & Celio, M. R. Calretinin-positive Cajal-Retzius cells persist in the adult human neocortex. Neuroreport 6, 1869-74 (1995).

61. Glezer, I. I., Hof, P. R. & Morgane, P. J. Calretinin-immunoreactive neurons in the primary visual cortex of dolphin and human brains. Brain Res 595, 181–8 (1992).

62. Miyoshi, G. et al. Genetic fate mapping reveals that the caudal ganglionic eminence produces a large and diverse population of superficial cortical interneurons. J Neurosci 30, 1582–94 (2010).

63. Lein, E., Borm, L. E. & Linnarsson, S. The promise of spatial transcriptomics for neuroscience in the era of molecular cell typing. Science 358, 64–69 (2017).

64. Colantuoni, C. et al. Temporal dynamics and genetic control of transcription in the human prefrontal cortex. Nature 478, 519–23 (2011).

65. Kang, H. J. et al. Spatio-temporal transcriptome of the human brain. Nature 478, 483–489 (2011).

66. Bahney, J. & Bartheld, C. S. von. The Cellular Composition and Glia-Neuron Ratio in the Spinal Cord of a Human and a Nonhuman Primate: Comparison with Other Species and Brain Regions. The Anatomical Record 301, 697–710 (2017).

67. Bjugn, R. The use of the optical disector to estimate the number of neurons, glial and endothelial cells in the spinal cord of the mouse-with a comparative note on the rat spinal cord. Brain Res 627, 25–33 (1993).

68. Lassek, A. M. & Rassmussen, G. L. The human pyramidal tract: A fiber and numerical analysis. Archives of Neurology & Psychiatry 42, 872–876 (1939).

69. Finlay, B. & Darlington, R. Linked regularities in the development and evolution of mammalian brains. Science 268, 1578–1584 (1995).

70. Markou, A., Chiamulera, C., Geyer, M. A., Tricklebank, M. & Steckler, T. Removing obstacles in neuroscience drug discovery: the future path for animal models. Neuropsychopharmacology 34, 74–89 (2009).

71. Nestler, E. J. & Hyman, S. E. Animal models of neuropsychiatric disorders. Nature Neuroscience 13, 1161–1169 (2010).

72. Sorensen, S. A. et al. Correlated Gene Expression and Target Specificity Demonstrate Excitatory Projection Neuron Diversity. Cerebral Cortex 25, 433–449 (2013).

73. Aronesty, E. Comparison of Sequencing Utility Programs. The Open Bioinformatics Journal 7, 1–8 (2013).

74. Dobin, A. et al. STAR: ultrafast universal RNA-seq aligner. Bioinformatics 29, 15–21 (2012).

75. Lawrence, M. et al. Software for Computing and Annotating Genomic Ranges. PLoS Computational Biology 9, e1003118 (2013).

76. Calvo, S. E., Clauser, K. R. & Mootha, V. K. MitoCarta2.0: an updated inventory of mammalian mitochondrial proteins. Nucleic Acids Res 44, D1251–7 (2016).

77. Lein, E. S. et al. Genome-wide atlas of gene expression in the adult mouse brain. Nature 445, 168–76 (2007).

78. Lyubimova, A. et al. Single-molecule mRNA detection and counting in mammalian tissue.Nat Protoc 8, 1743–58 (2013).

79. Crow, M., Paul, A., Ballouz, S., Huang, Z. J. & Gillis, J. Characterizing the replicability of cell types defined by single cell RNA-sequencing data using MetaNeighbor. Nat Commun 9, 884 (2018).

80. Yanai, I. et al. Genome-wide midrange transcription profiles reveal expression level relationships in human tissue specification. Bioinformatics 21, 650-9 (2005).

